# Robust Hi-C chromatin loop maps in human neurogenesis and brain tissues at high-resolution

**DOI:** 10.1101/744540

**Authors:** Leina Lu, Xiaoxiao Liu, Wei-Kai Huang, Paola Giusti-Rodríguez, Jian Cui, Shanshan Zhang, Wanying Xu, Zhexing Wen, Shufeng Ma, Jonathan D Rosen, Zheng Xu, Cynthia Bartels, Riki Kawaguchi, Ming Hu, Peter Scacheri, Zhili Rong, Yun Li, Patrick F Sullivan, Hongjun Song, Guo-li Ming, Yan Li, Fulai Jin

## Abstract

Genome-wide mapping of chromatin interactions at high resolution remains experimentally and computationally challenging. Here we used a low-input “easy Hi-C” (eHi-C) protocol to map the 3D genome architecture in neurogenesis and brain tissues, and also developed an improved Hi-C bias-correction pipeline (*HiCorr*) enabling better identification of enhancer loops or aggregates at sub-TAD level. We compared ultra-deep 3D genome maps from 10 human tissue- or cell types, with a focus on stem cells and neural development. We found several large loci in skin-derived human iPSC lines showing recurrent 3D compartmental memory of somatic heterochromatin. Chromatin loop interactions, but not genome compartments, are hallmarks of neural differentiation. Interestingly, we observed many cell type- or differentiation-specific enhancer aggregates spanning large neighborhoods, supporting a phase-separation mechanism that stabilizes enhancer contacts during development. Finally, we demonstrated that chromatin loop outperforms eQTL in explaining neurological GWAS results, revealing a unique value of high-resolution 3D genome maps in elucidating the disease etiology.

**Highlights:**

- Low input “easy Hi-C” protocol compatible with 50-100K cells
- Improved Hi-C bias correction allows direct observation and accurate identification of sub-TAD chromatin loops and enhancer aggregates
- Recurrent architectural memory of somatic heterochromatin at compartment level in skin-derived hiPSCs
- Chromatin loop, but not genome compartment, marks neural differentiation
- Chromatin loop outperforms eQTL in defining brain GWAS target genes

## Main

Hi-C has transformed our understanding of mammalian genome organization^1,2^. In the past decade, with increasing sequencing depth, a hierarchy of 3D genome structures, such as compartment A/B^1^, topological domains or topological associated domains (TADs)^3,4^ were revealed. More recently, kilobase resolution Hi-C analysis was achieved with sequencing depth at billion-read scale^5,6^. At this resolution, it is possible to discern specific chromatin loops between *cis*-regulatory elements. The information inherent in the 3D genome, especially chromatin loops, is critical for understanding the genetics of complex diseases^5,7–9^, such as the GWAS of cognitive traits and psychiatric disorders^10,11^.

However, kilobase-resolution Hi-C analysis is challenging both experimentally and computationally, especially when the amount of starting material is small. Experimentally, it is important to develop low-input Hi-C protocols that can deliver high quality libraries for ultra-deep sequencing. Computationally, mapping chromatin interactions with Hi-C at high-resolution suffers from the difficulty to correct the data biases, which leads to the low reproducibility or coverage in loop calling^12^. For example, the commonly used genome-wide loop caller HICCUPS yields ~10^4^ CTCF loops^6^ that only explain a small number of GWAS hits; Several recent Hi-C studies called SNP/gene interactions with locus-focused algorithms^11,13,14^, but these algorithms are not suitable for unbiased genome-wide loop calling and usually have strong biases towards selected loci and a high false positive rate. Alternatively, other studies using targeted capture Hi-C, ChIA-PET, HiChIP, *etc*.^15–20^ reported many more promoter-enhancer interactions, even though those methods are incomprehensive, biased due to target selection, and sometimes require even more biomaterials than Hi-C. Currently there is not a consensus whether Hi-C is a viable option to map promoter-enhancer loops at sub-TAD level for transcription regulation and human disease studies.

To address these challenges, we developed a new genome-wide Hi-C bias-correction pipeline that substantially improved the mapping of enhancer-promoter chromatin loops at fragment resolution. We also developed a genome-wide all-to-all version of 4C^21^ protocol named “easy Hi-C” (eHi-C), which yields high complexity Hi-C libraries with fewer than 0.1 million cells as the starting material. With these new toolsets, we mapped chromatin loops in 10 (e)Hi-C datasets and revealed new insights into the transcriptional regulation and the genetics of human diseases.

### Design and performance of eHi-C

In Hi-C, 5’ overhangs are created after restrictive DNA digestion (*e.g.* with *HindIII*) so that ligation junctions can be labeled with biotinylated nucleotides and eventually enriched in a pull-down step with streptavidin beads. However, this biotin-dependent strategy has intrinsic limitations that prevent the use of Hi-C if only low cell inputs are possible, because the efficiency of biotin incorporation is low^22^, and the recovery rate of biotin-labeled DNA from the pull-down procedure can be variable.

We therefore developed eHi-C to circumvent the limitations of Hi-C by using a biotin-free strategy to enrich ligation products (**Fig. 1a**). The eHi-C protocol is essentially a genome-wide “all-to-all” version of 4C^21^, and only involves a series of enzymatic reactions. eHi-C is also closely similar to ELP, another biotin-free genome-wide method developed several years ago for fission yeast 3D genome analysis^23^. However, ELP does not remove contamination from several species of non-junction DNA, and < 4% of ELP reads represent proximity ligation events^23^. Our eHi-C protocol has several key improvements, which allows the generation of high-yield libraries from small amounts of input tissue (**Extended Data Fig. 1**, more discussion in **Supplementary Methods**). We tested low-input eHi-C with 0.1 million cells and found that the resulting DNA libraries had an equivalent complexity as published conventional Hi-C libraries with 10 million cells; the yield of *cis*-contacts from eHi-C libraries is also better than most of the published *HindIII*-based Hi-C libraries (**Extended Data Table 2-3** and **Extended Data Fig. 1g-h**). At low resolution, the contact heatmaps from Hi-C and eHi-C data are identical showing the same component A/B^1^ and TAD^3,4^ structures (**Extended Data Fig. 1i-j**). Finally, since eHi-C has a distinct error source and bias structure from conventional Hi-C due to protocol differences (**Supplementary Methods** and **Extended Data Fig. 2**), we have adjusted our data filtering and normalization method to unify the high-resolution analysis of both Hi-C and eHi-C data (see later discussion).

**Figure 1.**
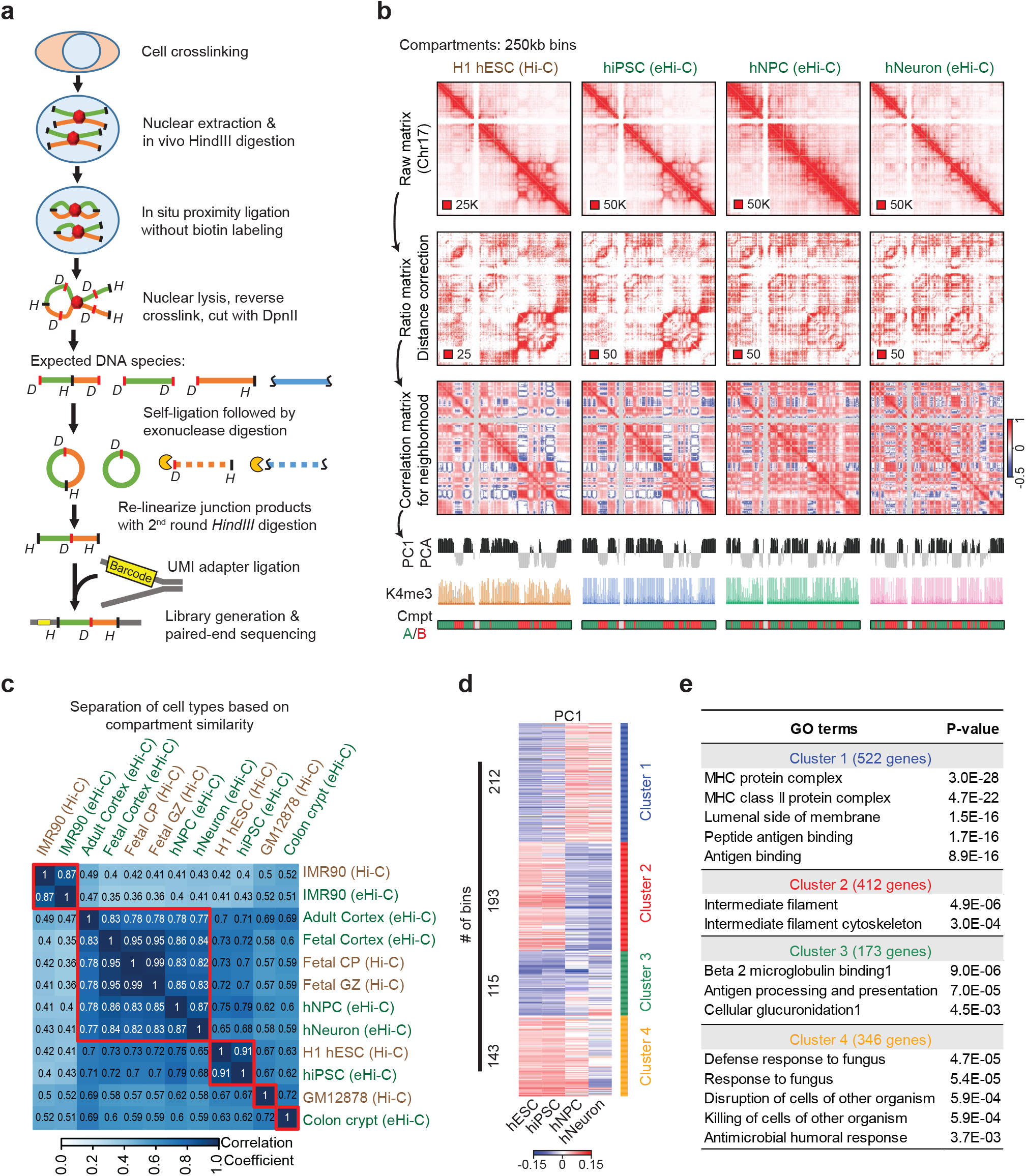
Mapping 3D genome with eHi-C. **a**, The scheme of eHi-C. **b**, Compartment level comparison of Hi-C or eHi-C data at 250kb resolution. Correlation matrices were created after two steps of data transformations showing the 3D genome neighborhood; PC1 from principal component analysis (PCA) were used to call compartment A/B. **c**, Heatmap showing the neighborhood similarity between 5 Hi-C and 7 eHi-C datasets (including a low-depth IMR90 eHi-C dataset). The correlation coefficient is computed by comparing the correlation matrices from different samples. **d**, Clustering the 663 bins (DCRs) based on PC1 values representing compartment switching. **e**, GO analysis of genes in each DCR clusters.

### Billion-read scale 3D genome datasets in 10 cell- or tissue-types

We have successfully performed eHi-C in multiple cell- and tissue types. Theoretically, the best Hi-C analysis resolution is determined by the restrictive endonuclease used (~2 kb for 6-base cutters, and ~200 bp for 4-base cutters). However, due to the lack of sequencing depth, high-resolution analysis at kilobase-scale is only achievable within 1-2 Mb. We estimated that for 6-base cutters, ~200 million mid-range (within 2 Mb) *cis*- contacts are required for fragment-level analysis (5-10 kb resolution); usually this translates into ~ 1-2 billion total non-redundant read pairs (**Supplementary Methods**).

Five of our eHi-C datasets meet this sequencing depth requirement, including hiPSC and derived hNPC and hNeuron and two postmortem brain tissues (fetal cerebrum and adult anterior temporal cortex) (**Extended Data Table 4**). The hNPCs and hNeurons were derived from hiPSCs using a previous established forebrain neuron-specific differentiation protocol^24,25^ (**Extended Data Fig. 3**). We also generated or obtained billion read-scale conventional Hi-C data for the H1 human embryonic stem cell (hESC), IMR90 (skin fibroblast)^5^, GM12878 (B-Lymphocyte line)^6,26^, and two developing human cerebral cortex samples (cortical plate, fetal CP; and germinal zone, fetal GZ)^11^ (**Extended Data Table 4,7**). Altogether, we have sufficient sequencing depth for fragment-resolution analysis in 10 tissue- or cell- types.

### Compartment analysis lacks the specificity to identify neural genes

Genome compartmentalization is known to associate with cell identity and gene regulation^1,27–29^. The analysis defined compartment A/B with the first principal component values (PC1)^1^, which represents the euchromatin/heterochromatin neighborhoods. As expected, hiPSC and hESC have very similar correlation matrices despite the difference in the Hi-C protocol; neural differentiation causes significant changes of genome compartments, consistent with previous reports^30,31^ (**Fig. 1b**). Clustering analysis further showed a highly tissue- or cell-type specific genome compartmentalization. Notably, all brain/neuron related samples clustered together, and the three fetal brain samples (two Hi-C and one eHi-C) formed the tightest sub-cluster (**Fig. 1c**). These results demonstrate the consistency between eHi-C and Hi-C at low resolution.

We identified 663 differentially compartmentalized regions (DCRs) associated with neural differentiation (**Fig. 1d**, **Supplementary Data 1**). Presumably, these neurogenesis DCRs are relevant to neural gene activation. However, although we observed a consistent correlation between H3K27ac occupancy and PC1 values (**Extended Data Fig. 4**), gene ontology analysis failed to identify neuron related terms in the neurogenesis DCRs (**Fig. 1e**). This may be due to: (1) neurogenesis drives large changes in compartment beyond neuronal genes; (2) neuronal gene activation may occur without changing the genome compartments. We conclude that high-resolution chromatin loop analyses are required for accurate dissection of the connection between neurogenesis and genome architecture.

### H3K9me3 marks a compartment barrier in reprograming human skin fibroblast

Unexpectedly, we also observed a few visible ultra-large loci with compartmental differences between hiPSC and hESC (reprogramming DCRs) in chr6, chr21 and chr22 (**Fig. 2a**, **Extended Data Fig. 5-6**). This is surprising because a previous Hi-C study in mouse cells did not find compartment-level differences between mESC and miPSCs derived from pre-B, MEF, macrophage and neuron stem cells^31^. We reanalyzed these published mouse Hi-C datasets and indeed confirmed the lack of large reprogramming DCRs in miPSCs (**Extended Data Fig. 7**). We also mapped genome compartments in several other hiPSC lines and found that these large reprogramming DCRs are recurrent only in skin-derived hiPSCs. Specifically, we found DCR^chr6^ recurrent in 2 of 3 skin-hiPSCs, DCR^chr21^ and DCR^chr22^ recurrent in all skin-hiPSCs (**Extended Data Fig. 6**).

**Figure 2.**
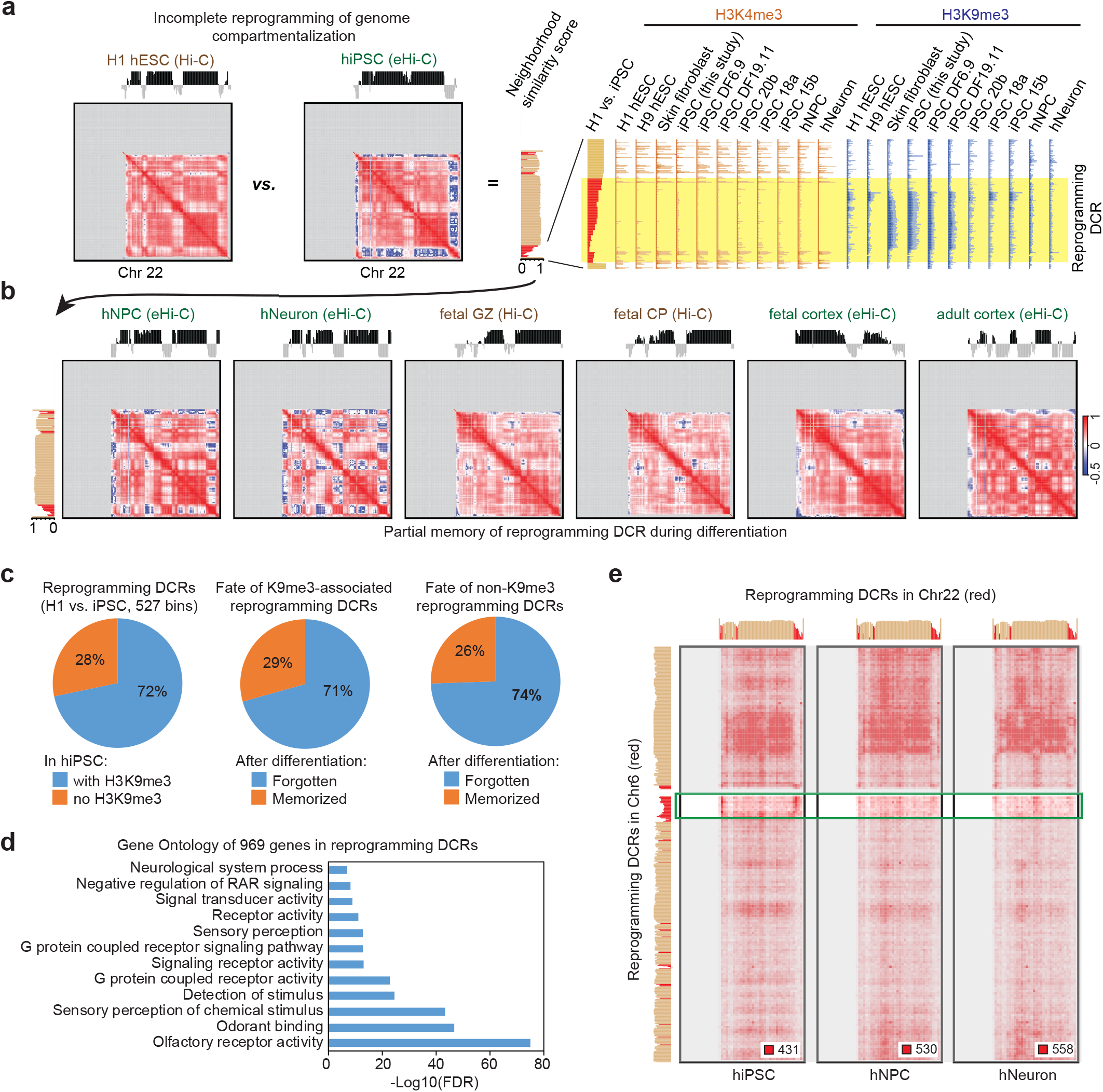
H3K9me3 marks an architectural barrier of induced pluripotency. **a**, A large reprograming DCR in chr22 identified by comparing H1 hESC and hiPSC. ChIP-seq tracks at a major DCR show the incomplete removal of H3K9me3 mark. **b**, Comparison of neighborhood profiles during neuronal differentiation and in brain tissues shows partial memory of reprogramming DCRs. **c**, Pie charts summarizing the H3K9me3 binding, and the fate of reprograming DCRs. **d**, Olfactory receptor genes are enriched in reprogramming DCRs. **e**, *Trans*- interaction signal (1Mb resolution) can be observed between reprogramming DCRs in hiPSCs but not after differentiation.

With a more sensitive method, we also identified many smaller reprogramming DCRs in skin-derived hiPSCs (**Supplementary Method**), and found that most reprogramming DCRs are associated with H3K9me3 in hiPSCs and enriched with olfactory receptor genes (**Fig. 2c-d**). The H3K9me3 mark at reprograming DCRs appears to originate from skin fibroblasts, which are also observed in several other hiPSC genomes (**Fig. 2a**, and **Extended Data Fig. 5a**). These results are consistent with the concept that H3K9me3 is an epigenetic barrier for human iPSC reprogramming^32^. Additionally, even though H3K9me3 is known to be part of compartment B heterochromatin, we observed from the correlation matrices that bins in the large reprogramming DCRs only correlate with themselves, not any other bins, regardless in compartment A or B (**Fig. 2a**, and **Extended Data Fig. 5a**). Furthermore, we also observed inter-chromosomal interactions between DCR^chr6^ and DCR^chr22^ in hiPSCs but not hNPC or hNeuron (**Fig. 2e**), suggesting that the large reprogramming DCRs formed an isolated neighborhood distinct from compartment A/B. Taken together, our results demonstrated a particularly strong 3D genome memory associated with H3K9me3 in human skin fibroblast reprogramming.

We further investigated the fate of reprogramming DCRs during neural differentiation. Most reprogramming DCRs lost (“forgot”) their iPSC-specific architecture and acquired a similar neighborhood profile to hESCs and brain tissues (**Fig. 2c**, example in **Extended Data Fig. 5**). Interestingly, some reprogramming DCRs still partially retain architectural “memory” despite losing their H3K9me3 mark in hNPCs and hNeurons (such as DCR^chr22^ in **Fig. 2a-b**). Such architectural memory may not have a major impact on neurogenesis since the hiPSCs successfully differentiated with no obvious defects. However, it remains a question whether the memory of DCRs causes subtle differences in neuron function, or whether it affects other cell lineages. It will be also relevant to test if cell-of-origin affects the functions of hiPSC-derived neurons, and to test whether DCR memory contributes to those differences.

### An improved Hi-C normalization method for chromatin loop identification

Identifying chromatin loops, especially the enhancer-promoter interactions at sub-TAD level, remains a major bioinformatic challenge in Hi-C analysis, as it is increasingly difficult to correct biases when the resolution increases to single fragment level^12^. We previously developed a method to explicitly correct fragment size, distance, GC content and mappability biases, and to estimate the expected frequency between any two fragments^5,33^. Using joint function, this method can correct the interaction effects between parameters (e.g. the interaction between fragment size and distance). However, this explicit method does not correct biases from unknown sources. Alternative strategies, such as VC normalization^1^, ICE^34^ and Knight-Ruiz (KR) matrix-balancing algorithms^6^, correct both known and unknown biases by normalizing a “visibility” factor (usually the total read counts) for each locus, with or without iterations. However, these implicit methods assume all biases are hidden in the visibility factor, and the visibility between different loci are independent; these assumptions are questionable at high-resolution within short to mid-ranges (more discussion in **Supplementary Methods**). For example, implicit methods do not correct the biases from distance or the size constraint Hi-C or eHi-C libraries (**Extended Data Fig. 2f**).

We developed a new strategy that corrects the implicit “visibility” factor after normalizing all aforementioned known biases, that consequently has the advantages of both explicit and implicit methods. The new method still estimates expected values for every fragment pair and uses observed / expected ratios to determine chromatin interactions (**Fig. 3a**, **Supplementary Methods**). Importantly, we computed the “visibility” only using the *trans*-reads. This is because normalizing *cis*- visibility has the risk of over-correction since many *cis*-reads come from chromatin loops (**Fig. 3b**): we found that *cis-* visibility is higher at histone marked loci and repetitive elements. The latter is possibly due to the widespread contribution of transposable elements to transcriptional regulatory sequences in the mammalian genome^35^(**Fig. 3c-e**). From the ratio heatmaps, we can directly observe discrete chromatin loops without the interference from local DNA packaging signal along the diagonal. Compared to other normalization methods (including matrix balancing methods), our approach significantly improves the peak calling and does not have the over-correction problem at the short range (**Fig. 3f**, compare the last column with other columns).

**Figure 3.**
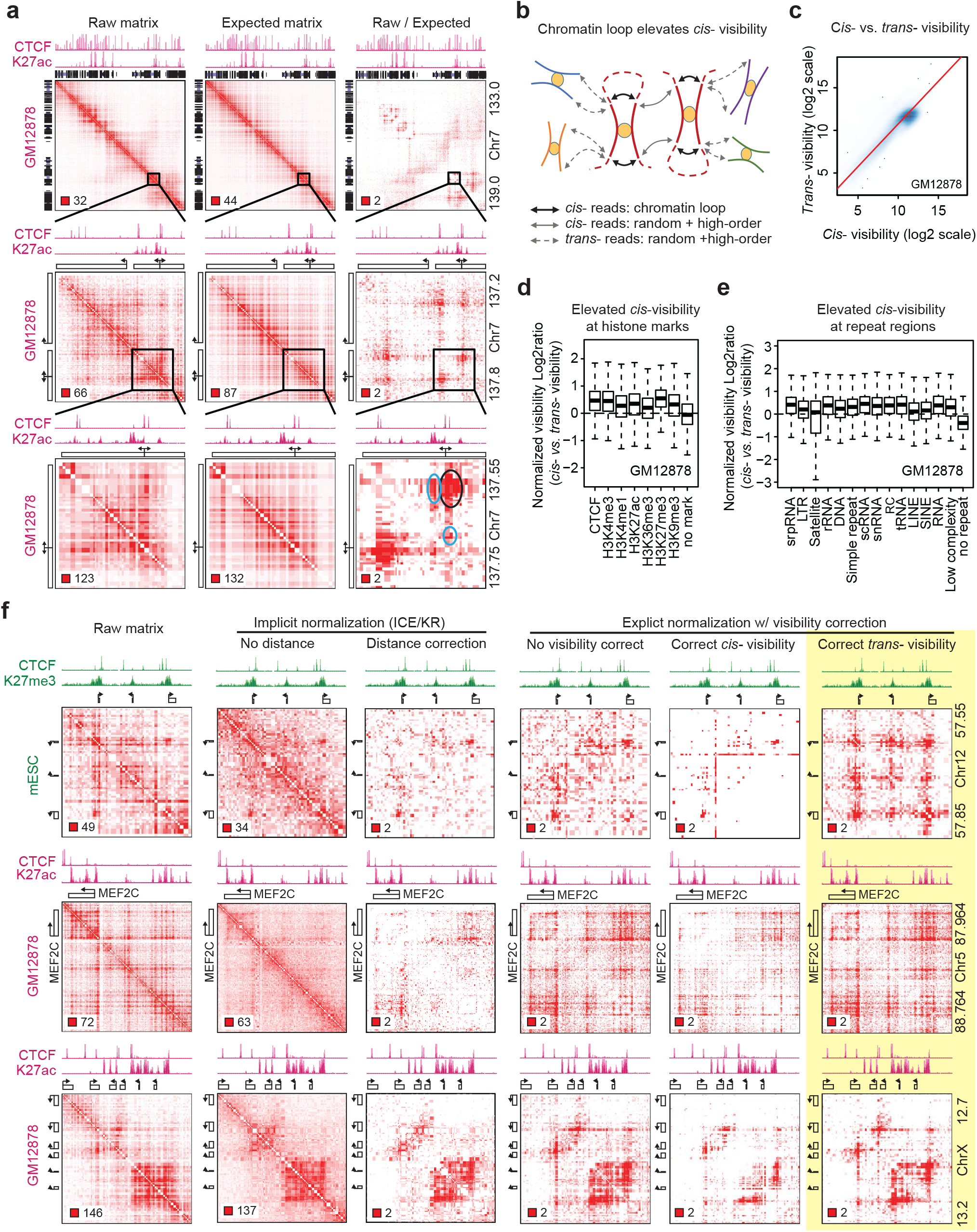
Improving the chromatin looping identification from Hi-C data. **a**, Raw, expected, and ratio heatmaps at different zooming scales. Note that chromatin loop is not obvious before correction. **b**, Chromatin loops contributes to *cis* but not *trans* Hi-C reads of a restrictive fragment, leading to an elevated *cis*/*trans* visibility ratio. **c**, Scatter plot of all fragments in GM12878 Hi-C data showing a skew towards higher *cis*- than *trans*- visibility. **d-e**, Epigenetically marked regions and repeat elements have a higher *cis*/*trans* visibility ratio. **f**, Comparing the results of different visibility correction methods.

### Identifying chromatin loops and enhancer aggregates from Hi-C

The improved bias-correction allows convenient identification of chromatin loops as red pixels from ratio heatmaps (**Supplementary Methods**). With sequencing depth at 150~200 million mid-range contacts, the new loop caller identified 60~150K loop pixels in different samples with a high reproducibility at 40~60% between biological replicates (**Extended Data Fig. 8**); inadequate sequencing depth appears to be the major reason for non-reproduced loops (**Extended Data Fig. 8b**). As a “global enrichment” strategy, our loop caller does not make prior assumptions about the distance, shape, size, or density of chromatin loops^12^, therefore identifies more enhancer/promoter chromatin loops than alternative methods (**Extended Data Fig. 8e-g**, more discussion in **Supplementary Methods**).

For example, **Fig. 4a** shows an example of a GM12878-specific enhancer aggregate, revealing discrete loop peaks with various shapes and sizes in the ratio heatmap. Four major enhancers (size ranging from 10kb to 30kb) appear to mediate these chromatin interactions, since the identical CTCF binding sites in H1 and IMR90 are not sufficient to create these interactions (**Fig. 4a**). This example is reminiscent of a “phase separation” model in which individual enhancers in a super-enhancer interact with each other via the condensation of transcription factors and cofactors^36^. However, this enhancer aggregate encompasses >150 kilobase, well beyond the size of a super-enhancer. When any of the four enhancers was repressed by dCas9-mediated enhancer silencing^37^, we observed the loss of enhancer mark on all enhancers (**Extended Data Fig. 9**), and the downregulation of two GM12878-specific genes (*LINC00158* and *MIR155HG*) in this enhancer aggregate (**Fig. 4b-c**), suggesting that all clustered enhancers in this example function in a coordinated fashion. Interestingly, the expression of two nearby genes (*MRPL39* and *JAM2*) are also GM12878-specific and dependent on the enhancer aggregate, possibly through mechanisms that do not require direct chromatin interactions^38^.

**Figure 4.**
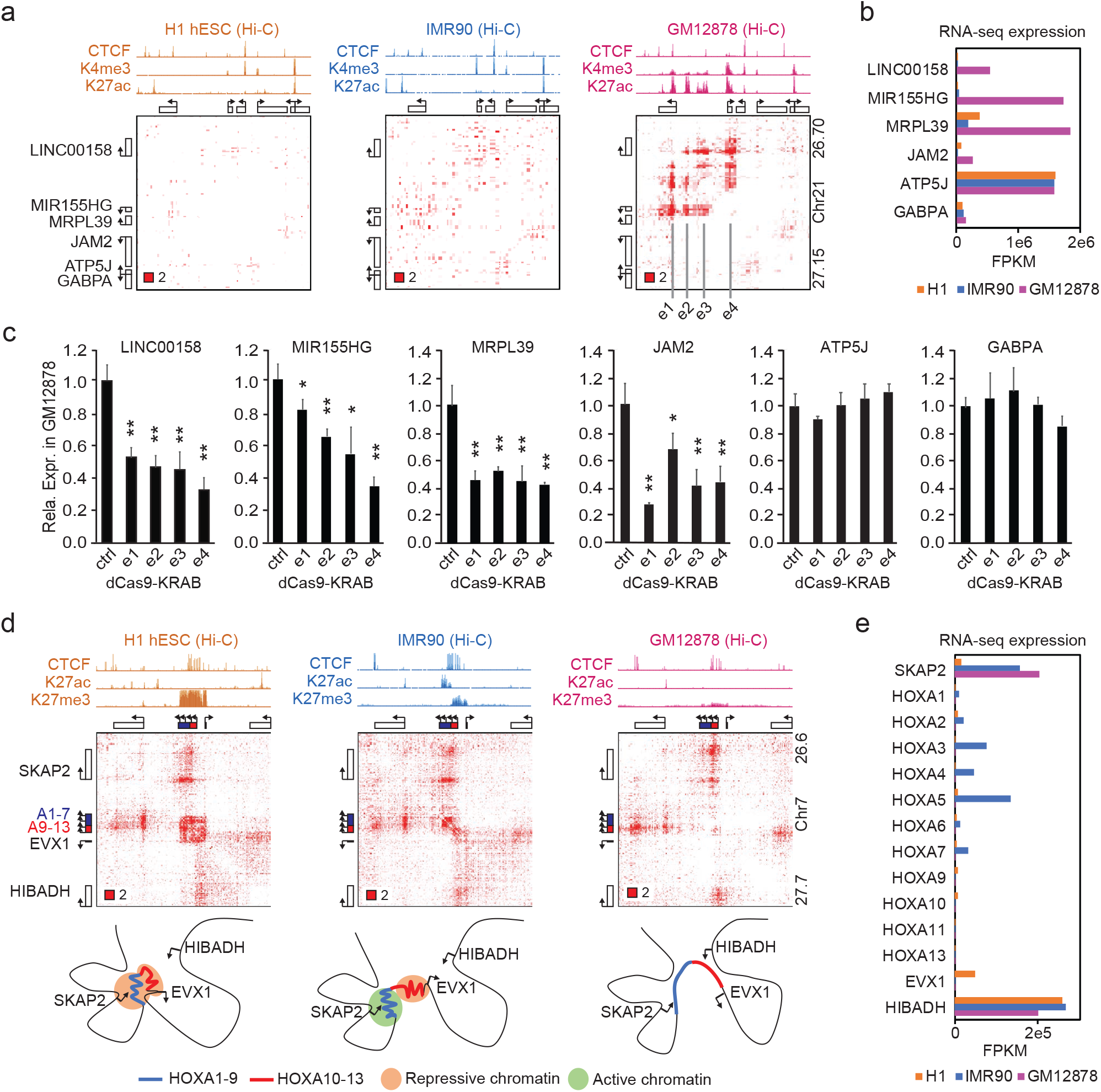
Cell type-specific chromatin loops or enhancer aggregates. **a-b**, The bias corrected Hi-C heatmaps at a GM12878-specific enhancer aggregate (**a**), and the transcription levels of the six genes in this region (**b**). **c**, The expression levels of every gene when the four enhancers indicated in (**a**) are repressed using CRISPRi; data are representative from > 3 independent experiments. Error bar: *s.d*. of 3 PCR replicates; * p < 0.05, ** p < 0.01 in *t* test. **d**, Architecture of HoxA gene cluster in H1, IMR90 and GM12878 cells. **e**, Expression of HoxA genes in these three cell types.

With the removal of the local DNA packaging signal, we can also distinguish chromatin compaction events as red pixel domains. The best example is the Polycomb group (PcG) associated chromatin domain at HOXA gene family^39–41^. The normalization dimmed the up- and downstream TAD signal and allowed direct observation of the ESC-specific repressive chromatin domain at HOXA genes, which splits or dissolves when it loses some or all the H3K27me3 mark in IMR90 and GM12878 cells (**Fig. 4d-e**).

### Enhancer loops and aggregates mark neuronal differentiation but not gene activation

We are particularly interested in identifying enhancer aggregates associated with neural differentiation, since they may represent a 3D genome signature for the neuronal lineage. To do this, we first identified 323,700 loop pixels in total from hiPSC, hNPC and hNeuron cells, each with ~140K pixels (**Fig. 5a**). The overlap between hNeurons and hNPCs is greater than their overlap with hiPSCs (**Fig. 5a**). Insulators (with CTCF), promoters (with H3K4me3), and enhancers (with H3K27ac) are clearly top contributors to chromatin loops. Interestingly, the numbers of enhancer- or promoter interactions are higher in hNPCs and hNeurons than in hiPSCs (**Fig. 5b**). The genes involved in hNPC and hNeuron chromatin loops are strongly associated with neuronal differentiation functions (**Fig. 5d**). This is in contrast to the results from dynamic compartment analyses (**Fig. 1d-e**), confirming that chromatin loops, but not high-order genome compartmentalization, are the hallmarks of neuronal differentiation.

**Figure 5.**
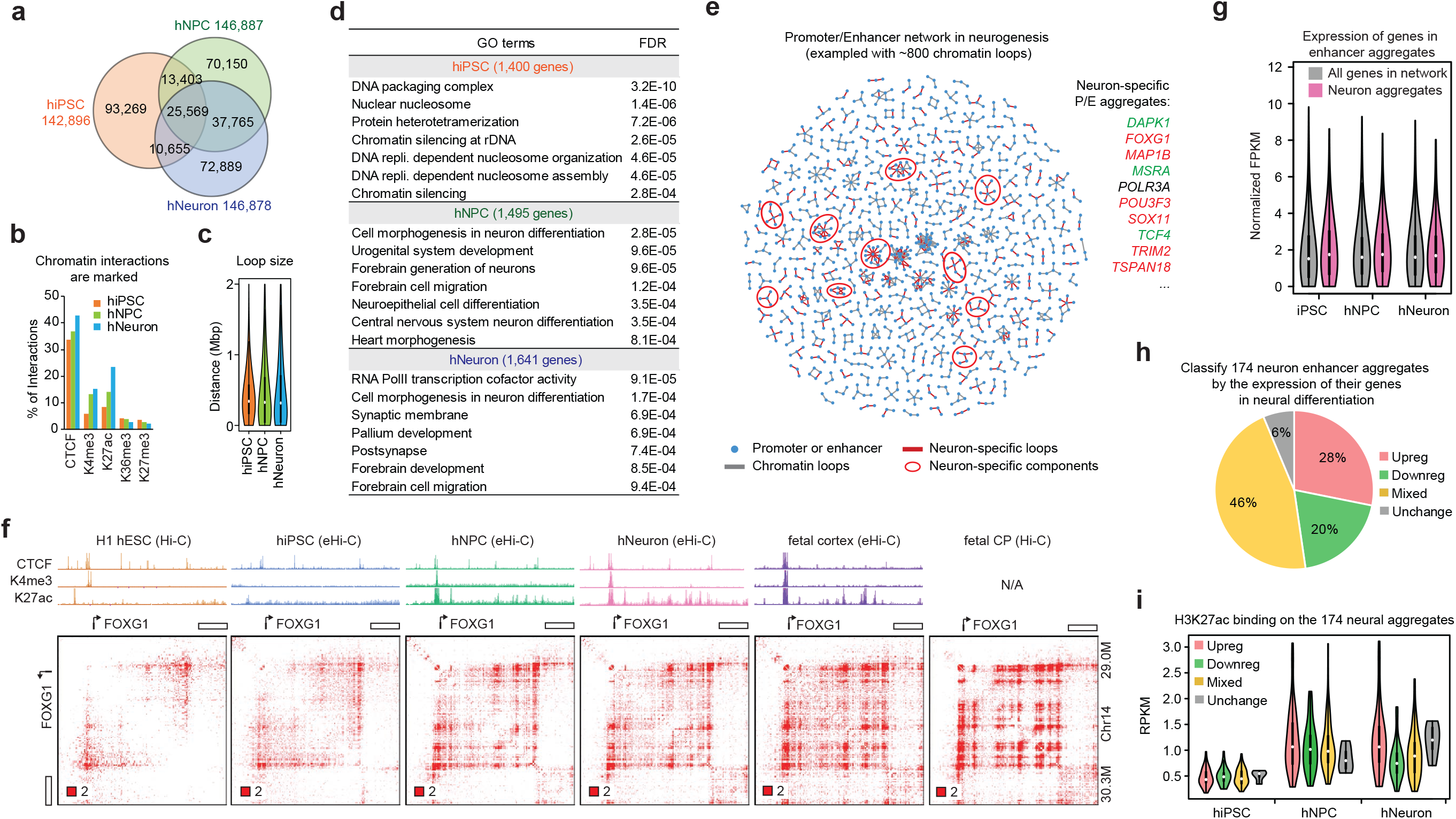
Chromatin loops in neural differentiation. **a**, Venn diagram showing the overlap between chromatin interactions from hiPSCs, hNPCs and hNeurons. **b**, Bar graph showing the percentage of chromatin interactions with various histone marks. **c**, Distance distribution of chromatin loops in three cell types. **d**, Top gene ontology terms from genes involved in top 3,000 chromatin loop pixels ranked by ratio. **e**, An exemplary promoter-enhancer network with ~800 strongest chromatin loops during neurogenesis. Neuron-specific network components can be identified as candidate neuronal enhancer aggregates. Genes in a few neural enhancer aggregates are listed on the right: red, upregulated in neural differentiation, green, downregulated. **f**, Formation of enhancer aggregate at the *FOXG1* locus during neural differentiation. **g**, Summary of gene expression in neural enhancer aggregates. **h**, Classification of neural enhancer aggregates based on their dynamic gene expression during differentiation. **i**, H3K27ac occupancy at different categories of neural enhancer aggregates.

We next constructed a network of 6,067 promoters and 11,453 enhancers using the aforementioned chromatin loops. The network includes 1,939 connected components (*i.e.* connected subnetworks); nearly one-third (603) of them are candidate enhancer aggregates (multi-node clusters with at least five edges, **Extended Data Fig. 10**). We used the ratio of each loop pixel to measure the loop strength semi-quantitatively, and identified 174 neural enhancer aggregates in which the chromatin loops are strengthened in hNeurons compared to hiPSCs (**Supplementary Methods, Supplementary Data 2**). As expected, the neural enhancer aggregates contain key neural genes, including *FOXG1, POU3F3*, *SOX11*, and *TCF4* (**Fig. 5e**). Independent Hi-C data from hESCs and primary brain tissues supported the gain of enhancer or promoter loops at these loci during neural differentiation (**Fig. 5f**, more examples in **Extended Data Fig. 11**). Interestingly, many of these enhancer aggregates are substantially strengthened in primary brain tissues, sometimes form striking grid-like patterns (**Fig. 5f**), suggesting that hNPCs and hNeurons are in a transition phase of genome rewiring; with enhancers continuing to aggregate and stabilize during neuronal maturation.

It is however surprising that the neural enhancer aggregates do not correlate with gene activation (**Fig. 5g**). Our RNA-seq data revealed that the 174 neural enhancer aggregates contain both up- and down-regulated genes during neurogenesis (**Extended Data Fig. 11, Supplementary Data 2**), although they clearly gain higher overall H3K27ac occupancy in hNPC or hNeuron than in hiPSCs (**Fig. 5h-i**). Furthermore, we also observed continuous enhancer aggregation at several gene-dense regions in which genes are already active in hESCs and hiPSCs (marked by H3K4me3 and H3K27ac); these genes can be either up- or down-regulated in hNPCs and hNeurons in a coordinated fashion (**Extended Data Fig. 12, Supplementary Data 2**). These results indicate that enhancer aggregation may not necessarily result in gene activation (see more discussion below).

### Chromatin loop outperforms eQTL in identifying GWAS target genes

Finally, we explored our dataset to investigate the genetics of brain disorders. We collected 6,556 lead GWAS SNPs reported for a number of cognitive traits or brain-related disorders (including intelligence, autism, schizophrenia, Alzheimer’s disease, *etc*.)^42^ and defined their linkage disequilibrium (LD) using the latest TOPMed data (**Supplementary Methods**). We next called 14,943 distal GWAS SNP-promoter pairs (*i.e.* the predicted promoter is outside of the GWAS LD) using chromatin loop data (**Supplementary Data 3**).

We defined tier 1 neural loop predictions as the SNP-promoter pairs supported by loops from >=2 of the six neural (e)Hi-C datasets. There are 4,421 tier1 pairs involving 2,173 SNPs and 1,439 genes (**Fig. 6a**). Similarly, we also defined tier 2 and 3 loop predictions, which are supported by only one or zero neural (e)Hi-C datasets. Additionally, we also predicted distal GWAS target genes (outside of LD) using the GTEx *cis-*eQTL data from 48 human tissues^43^, including 14 neural tissues (13 brain tissues and nerve tibial) (**Supplementary Data 3**). The overlap between loop and eQTL predictions is modest: 10.4%, 7.8%, and 7.5% of tier 1, 2, and 3 neural loop predictions are supported by neural eQTLs. However, non-neural eQTL data also have a similar trend (18.4%, 14.7% and 15.4% for tier 1-3 loop predictions, **Fig. 6a**), suggesting a lack of tissue specificity.

**Figure 6.**
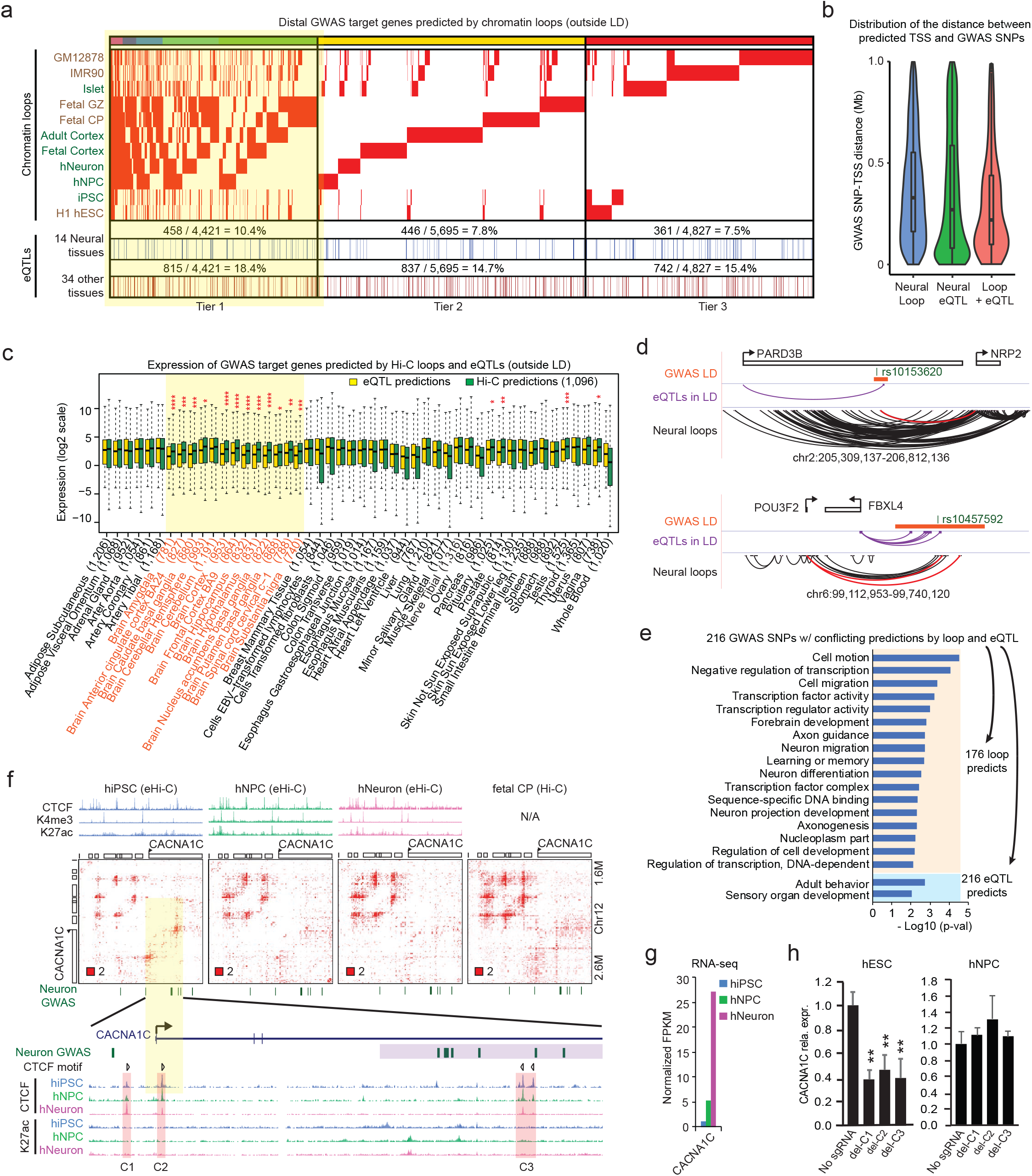
Chromatin loop outperforms eQTLs in explaining GWAS results. **a**, Heatmap showing the chromatin loop predicted GWAS target genes, and their overlap with GTEx eQTL data. Highlighted: Tier 1 neural predictions supported by at least two neural Hi-C datasets. **b**, Distance distribution of predicted GWAS SNP-TSS pairs, based on whether they are supported by loop, eQTL, or both. **c**, Wilcox rank sum tests were used to test if the loop predicted GWAS target genes are expressed at higher level than eQTL predictions in 48 GTEx tissues.*p<1e-2,**p<1e-3,***p<1e-4,****p<1e-5; Highlighted: 13 brain tissues. Numbers in parenthesis: genes predicted using eQTL data from each tissue. **d**, Two GWAS loci examples for which neural loop and eQTL make conflicting predictions. **e**, GO terms enriched in loop or eQTL predicted target genes, when the two methods make conflicting predictions. **f**, The *CACNA1C* GWAS locus is associated with an hiPSC-specific CTCF loop. Highlighted are the three CTCF occupied regions and the CTCF motif directionality. **g**, Expression of *CACNA1C* during neurogenesis using RNA-seq data. **h**, CTCF deletion downregulates *CACNA1C* in hESC but not NPC. Data are representative from > 3 independent experiments. Error bar: *s.d*. of three PCR replicates; * p < 0.05, ** p < 0.01 in *t* test.

We therefore systematically compared the performance of chromatin loop and eQTL data in explaining GWAS results. We focused on tier 1 loop predictions only within 1Mb, since GTEx only called *cis*-eQTLs in this window (**Fig. 6b**). In 12 of the 13 GTEx brain tissues, the expression levels of the 1,096 loop target genes are significantly higher than eQTL target genes, even though the two methods predicted comparable number of genes (**Fig. 6c**); such difference is only observed in 4 of the 35 non-brain tissues, indicating that chromatin loop predictions have better brain specificity than eQTL predictions.

We further focused on the 216 GWAS SNPs for which chromatin loops and brain eQTLs made conflicting prediction of target genes (**Supplementary Data 3**). **Fig. 6d** shows two such examples: one locus (rs10153620) associated with attention deficit hyperactivity disorder (ADHD)^44^, and the other locus (rs10457592) associated with major depression^45^. In both examples, chromatin loop predicted key neuronal genes (*NRP2* and *POU3F2*), while brain eQTLs predicted genes with unclear brain functions (*PARD3B* and *FBXL4*). Most importantly, we found an overall trend that chromatin loops outperform eQTLs in identifying genes with known brain functions. For all of the 216 GWAS SNPs, Hi-C predicted 176 target genes, which enriched dozens of GO terms related to neural functions and transcription regulation (**Fig. 6e**, **Supplementary Data 3**). In contrast, the eQTL target genes only enriched two relevant GO terms at a p < 0.01 level, highlighting the value of chromatin loop data in explaining disease genetics (**Fig. 6e**, see discussion).

Interestingly, although we frequently observed neural loops at known brain GWAS loci, such as *MEF2C*, *CTNND1, TRIO*, and *DRD2* (**Extended Data Fig. 13**), some loci lose chromatin loops during neural differentiation. The best example of this is the GWAS locus located in the third intron of *CACNA1C*, which is one of the strongest and best-replicated associations for schizophrenia (SCZ) and bipolar disorder (BD)^46^. Past studies on this locus in neurons or brain tissues suggested a transcription regulatory role, but the causative variants are still unknown^47–50^. Unexpectedly, we found a strong hiPSC-specific CTCF loop connecting the GWAS locus to the *CACNA1C* promoter; this loop weakens when the gene is upregulated during neurogenesis and in brain tissues, possibly due to transcription elongation^51^ (**Fig. 6f-g**). *CACNA1C* has a low but detectable expression in hESCs. To test if the CTCF loop is functional, we deleted the three corresponding CTCF binding sites and found that *CACNA1C* is downregulated only in hESCs but not in hNPCs (**Fig. 6h**). These results raised a possibility that the *CACNA1C* GWAS locus may affect gene expression and disease during early development instead of in mature neurons, which is consistent with a recent mouse study showing that *CACNA1C* affects psychological disorders during embryonic development instead of adult neurons^52^. This example highlighted the importance of examining looping dynamics and cautions against only using brain or neuron data to investigate disease genetics.

## Discussion

We made several unexpected observations in this study. Firstly, we found a few ultra-large loci showing recurrent compartment level memory of heterochromatin in skin-derived human iPSC lines. These results are in contrast to a previous landmark study showing nearly complete reprogramming of mouse pre-B, MEF, macrophage and neuron stem cells^31^ to iPSCs at compartment level. This discrepancy is possibly due to species difference and the fact that skin fibroblasts are at a more terminally differentiated stage. Since skin fibroblasts are a popular source of cells for human somatic cell reprogramming, these regions may serve as a benchmark for gauging skin derived hiPSC pluripotency.

Our improved normalization method allows direct recognition and systematic identification of enhancer-promoter loops and loop aggregates from Hi-C or eHi-C heatmaps at sub-TAD level, with much less interference from the local DNA packaging signal. In many examples, the promiscuous TAD blocks in raw heatmaps become discrete promoter or enhancer interactions after correction. These results suggest that promoters and enhancers form stable CTCF-independent interactions and are dominant contributors to intra-TAD signal.

It is also unexpected that differentiation-gained enhancer aggregates do not correlate with gene activation in neural differentiation, since the enhancer “phase separation” model was initially proposed as a mechanism for trans-activation^36^. Notably, past and recent studies have revealed many phase separation mechanisms that organize both euchromatin and heterochromatin.^53^ We therefore speculate that even at enhancers, different *trans*- factors (protein or RNA) may create chromatin contacts during cellular differentiation, which do not necessarily cause gene activation. More studies are required on a case-by-case basis to tease out the underlying mechanisms, and to investigate whether the newly gained DNA contacts have gene regulatory functions.

Chromatin loops and eQTLs are two independent methods to identify long-range *cis*-regulatory relationships. When studying the function of non-coding variants, it is becoming common practice to look for evidence from both chromatin loop and eQTL data. However, our study showed a limited consistency between the two methods in predicting GWAS target genes: only a small fraction of looped GWAS loci are also supported by eQTLs. One possible explanation for this discrepancy is the lack of statistical power in eQTL detection, *i.e.* many *cis*-regulatory variants may not pass statistical significance due to: (i) limited population size; (ii) low minor allele frequency (MAF). However, the sensitivity issue cannot explain why loop appears to be more accurate than eQTL when the two methods make conflicting predictions (**Fig.6d-e**). Furthermore, a recent large blood eQTL study reported that after increasing the sample size to > 30,000 donors, although many more *cis*-eQTLs could be identified, they were mostly short-range eQTLs near promoters and had a different genetic architecture from GWAS SNPs^54^. The limited success of eQTLs in GWAS study highlighted another potential possibility that eQTLs obtained from healthy tissues may not reflect the gene regulatory landscape from patients. For example, a SNP may only have subtle effects on looped target gene in healthy donors, but plays a more prominent role when the locus gains a disease-specific enhancer in patients; in this scenario, chromatin loop can identify the correct target genes but eQTL from normal tissues cannot. Therefore, our results indicated that high-quality Hi-C loops have a unique value in the study of disease genetics: we should treat loops and eQTLs as two distinct lines of biological evidence in explaining GWAS results, rather than two mutually confirmatory datasets.

## Supporting information

SuppData 1

SuppData 2

SuppData3

## Acknowledgement

We would like to thank K.M. Christian, B. Ren and J. Dekker for comments. D. Johnson and J. Schnoll for lab coordination and support. This work was supported by grants from SFARI (to G.M. & F.J.), Mt Sinai Health Care Foundation (OSA510113 to F.J., OSA510114 to Yan Li) National Institutes of Health (R01HG009658 to F.J., R01DK113185 to Yan Li, P01NS097206, U19MH106434 and R37NS047344 to H.S., R35NS097370, U19AI13110, R01MH105128 to G-l.M., and K01MH109772 to P.G.R.). We gratefully acknowledge the studies and participants who provided biological samples and data for TOPMed (https://www.nhlbiwgs.org/).

## Code Availability

The source code for the improved Hi-C normalization can be found in https://github.com/JinLabBioinfo/HiCorr.

