## Supplemental information for "Robust Hi-C chromatin loop maps in human neurogenesis and brain tissues at high-resolution"

### Supplementary Methods

#### 1. Sample preparations

Cell lines: We used human primary IMR90 fibroblasts to test eHi-C performance. IMR90 cells were grown as previously described^1^. After confluence, the cells were detached with trypsin and collected by spinning down at 900g for 5 minutes. Then the cells were fixed in 1% formaldehyde for 15 minutes at 37ºC, followed by 1/20 volume of 2.5M glycine at room temperature for 5 minutes to quench formaldehyde. The fixed cells were washed in PBS and pelleted before stored in -80ºC. We generated additional conventional Hi-C libraries for H1 hESCs (WiCell, #WA01) because published Hi-C data in H1 hESC are not deep enough to support the fragment resolution analysis. H1 cells were cultured on the hESC-qualified Matrigel (Corning, #354277) coated plates in mTeSR1 medium (StemCell Technologies, #05850) before harvest for Hi-C analysis. The cell fixation protocol is the same as IMR90 cells. Additionally, we purchased another 3 human iPS cell lines from ATCC, 2 skin fibroblast derived (ATCC ACS-1011 and ATCC ACS-1019) and 1 bone marrow derived (ATCC ACS-1026). Cells were cultured and maintain on matrigel and mTeSR1 same as H1 cells. The cell fixation protocol is the same as IMR90 cells.

Neurogenesis samples: The hiPSC line used have been previously extensively characterized, including expression of pluripotent markers, karyotyping, lack of transgene integration, demethylation of promoter regions of pluripotent genes, in vitro differentiation into cell types of three germ layers and teratoma formation^2^^,^^3^. We followed our previously established protocol for forebrain-specific neuronal differentiation^3^. Briefly, hiPSC colonies were lifted by 1 mg/ml collagenase and cultured in non-treated polystyrene plates with embryoid body (EB) medium consisting of 20% KOSR, 2 μM dorsomorphin (Tocris) and 2 μM A83-01 (Tocris) for 7 days with daily medium changes. The EBs were then attached on matrigel to develop organized rosette-like structure and maintained in neural induction medium (hNPC medium) with an equal mixture of DMEM/F12 and Neural basal medium, N2 supplement, B27 supplement, NEAA, 2mg ml−1 heparin and 2 μM cyclopamine (Stemgent) for 16 days with medium change every second day. The neural rosettes were harvested mechanically and transferred to low attachment plates (Corning) in hNPC medium to form neural spheres for 3 days. hiNPCs were expanded as monolayer in hNPC medium after dissociation of neural spheres by Accutase. For neuronal differentiation, monolayer hiNPCs were switched to Neurobasal medium with 10 ng ml−1 BDNF, 10 ng ml−1 GDNF, GlutaMaxTM (Gibco) and B27 supplement. Quantification of different cellular markers was performed by analyzing a minimum of 500 cells from at least 4 randomly chosen fields of fluorescent images with ImageJ software. The cell fixation protocol is the same as IMR90 cells.

Brain tissues: For brain tissue analysis, anterior temporal cortex was dissected from postmortem samples from three adults of European ancestry with no known psychiatric or neurological disorder (Dr Craig Stockmeier, University of Mississippi Medical Center). Cerebra from three fetal brains were obtained from the NIH NeuroBiobank (gestational age 17-19 weeks), and none were known to have anatomical or genomic disease (**Extended Data Table 1**). Samples were dry homogenized to a fine powder using a liquid nitrogen-cooled mortar and pestle. All samples were free from large structural variants (>100 kb) detectable using Illumina OmniExpress arrays. Genotypic sex matched phenotypic sex for all samples. For easy Hi-C, Pulverized tissue (~100 mg) was crosslinked with formaldehyde (1% final concentration) and the reaction was quenched using glycine (150 mM). We lysed samples on ice with brain tissue-specific lysis buffer (10 mM HEPES; pH 7.5, 10 mM KCl, 0.1 mM EDTA, 1 mM dithiothreitol, 0.5% Nonidet‐40 and protease inhibitor cocktail). Samples are Dounce homogenized before HindIII digestion.

Colon crypt tissues: Crypts were dissected from non-cancer colon mucosa. After removing from the patient, we first cut away non–colon mucosa as much as possible, such as muscles, blood vessels and fat. The tissue was then treated with cell dissociation solution to pop out crypts from surrounding mucosa tissue. The suspension was filtered through a cell strainer to remove remaining tissue pieces. Pelleted crypts were crosslinked in 1% formaldehyde followed by glycine quenching. The fixed crypts were used for eHi-C as described below.

#### 2. ChIPmentation

We used ChIPmentation^4^ to map histone modification and/or CTCF in different samples. Briefly, cells and tissues were fixed in 1% formaldehyde at room temperature for 15 minutes followed by glycine quenching. To isolate nuclei, we lysed brain tissues with a specific lysis buffer (10 mM HEPES; pH 7.5, 10 mM KCl, 0.1 mM EDTA, 1 mM dithiothreitol (DTT), 0.5% Nonidet‐40 and protease inhibitor cocktail) for 10 minutes at 4°C. For cell cultures, we used lysis buffer 1 (50 mM HEPES; pH 7.5, 140 mM NaCl, 1 mM EDTA, 10% glycerol, 0.5% Nonidet‐40, 0.25% Triton X-100 and protease inhibitor cocktail) for 10 minutes at 4°C. The collected nuclei were then washed with a lysis buffer II (200mM NaCl, 1mM EDTA pH8.0, 0.5mM EGTA pH8.0, 10mM Tris-Cl pH8.0 and protease inhibitor cocktail) for 20 minutes at room temperature. The nuclei were pelleted at 1,800g for 10 minutes at 4°C and then resuspended in lysis buffer III (10mM Tris-Cl pH8.0, 100mM NaCl, 1mM EDTA, 0.5mM EGTA, 0.1% Na-Deoxycholate, 0.5% N-lauroylsarcosine and protease inhibitor cocktail) for sonication. The chromatin was sheared for 10 cycles (15 seconds on and 45 seconds off at constant power 3) on Branson 450 sonifier. 20-50ug of chromatin was used for each H3K4me3 (Abcam, ab8580)/ H3k27Ac (Abcam, ab4729)/ H3K27me3 (07-449)/ H3K36me3 (ab9050)/ H3K9me3 (ab8898) pulldown and 100-150ug for each CTCF (Abcam, ab70303) pulldown. First, 11ul of Dynabeads M-280 (Life Technologies, Sheep Anti-Rabbit IgG, Cat# 11204D) was washed three times with 0.5mg/ml of BSA/PBS on ice and then incubated with designed antibody for at least 2 hours at 4°C. The beads/antibody complexes were then washed with BSA/PBS. The pulldown was done in binding buffer (1% Trixon-X 100, 0.1% Sodium Deoxycholate and protease inhibitor cocktail in 1X TE) by mixing the beads/antibody complexes and chromatin. After pulling down for overnight, the beads/antibody/chromatin complexes were washed with RIPA buffer (50mM HEPES pH8.0, 1% NP-40, 0.7% Sodium Deoxycholate, 0.5M LiCl, 1mM EDTA and protease inhibitor cocktail). The beads complexes were then subjected to ChIPmentation by incubating with homemade Tn5 transposase in tagmentation reaction buffer (10mM Tris-Cl pH8.0 and 5mM MgCl2) for 10 minutes at 37°C. To remove free DNA, beads were washed twice with 1x TE on ice. The pulldown DNA was recovered by reversing crosslink for overnight followed by PCRClean DX beads purification. To generate ChIP-seq libraries, PCR was applied to amplify the pulldown DNA with illumina nextera primers. Size selection was then done with PCRClean DX beads to choose the fragments ranging from 100bp to 1000bp.

#### 3. Easy Hi-C

##### 3.1. The overview of eHi-C design

In Hi-C, 5’ overhangs are created after restrictive DNA digestion (*e.g.* with *HindIII*) so that ligation junctions can be labeled with biotinylated nucleotides and eventually enriched in a pull-down step with streptavidin beads. However, this biotin-dependent strategy has several intrinsic limitations that prevents the application of Hi-C in rare tissue or small cell populations. First, the efficiency of biotin incorporation into DNA is usually ~20-30%, sometimes as low as 5%^5^. Therefore, a majority of ligation junctions cannot be recovered. Second, only a portion of labeled ligation junction products can be pulled-down after several washes, further lowering the recovery rate. Lastly, extensive washes are required in the biotin-pulldown procedure to effectively remove contamination of un-ligated DNA products, but this will significantly reduce the library complexity.

We reasoned that we might circumvent the limitations of Hi-C by using a biotin-free strategy to enrich ligation products, thus improving the assay efficiency. Inspired by the biotin-free strategies used in 4C^6^ and ELP^7^, we developed eHi-C, which only involves a series of enzymatic reactions to generate DNA libraries for the mapping of genome architecture (**Fig. 1a**). In this protocol, we begin with the *in situ* proximity ligation procedure and performed *HindIII* digestion and proximity ligation while keeping nuclei intact^8-10^. In eHi-C, *HindIII* digested chromatin are ligated without end repair, leading to *HindIII*-digestible junction products (**Fig. 1a**). After nuclear lysis and reverse crosslinking, the DNA are digested with more frequent 4-base cutter *DpnII* before self-ligation. DNA with *DpnII* restrictive overhangs on both ends, including ligation junction products, will form circles. We used exonuclease to remove DNA that failed to form circles, as well as contaminations from un-ligated ends and other linear DNA species. At last, we cut the circularized DNA again with *HindIII*; only re-linearized junction DNA will be sequenced (**Fig. 1a**).

The eHi-C method is essentially a genome-wide “all-to-all” version of 4C and also closely similar to ELP, another biotin-free genome-wide method developed several years ago to identify DNA contacts in fission yeast^7^. However, the design of ELP was flawed because it cannot remove contaminations from several species of non-junction DNA (**Supplementary Fig. 1a**). As a result, less than 4% of ELP reads represent proximity ligation events^7^. The eHi-C protocol solves this issue by introducing an exonuclease digestion step. Additionally, because all reads from ELP are next to *HindIII* sites, it cannot distinguish PCR duplicates from reproducible ligation events between the same pair of *HindIII* ends (**Extended Data Fig. 1b**). Our eHi-C method addresses this issue with a custom adapter with random barcode as a unique molecule index (UMI) (**Fig. 1a,** **Supplementary Fig. 1c**). We also used *in situ* ligation in eHi-C to improve the library quality (**Fig. 1a**). Taken together, we have significantly optimized the eHi-C strategy to obtain high quality libraries for ultra-deep sequencing from small-scale bio-samples, which is not feasible with the original ELP method.

Because there is no DNA loss in its protocol (**Fig. 1a**), eHi-C should have a higher recovery rate of ligation junction products than conventional Hi-C, which is important for the analyses of small cell populations. The only exception is the exonuclease digestion step: Ligation junction DNA may be digested if they fail to self-ligate (**Fig. 1a**). From a control experiment, we determined that the efficiency of the self-ligation reaction is high (~60%, **Supplementary Fig. 1d**).

##### 3.2. Easy Hi-C protocol

In this study, low-input eHi-C libraries were prepared in two settings. In the first scenario (“aliquot” setting), we started with 1 million IMR90 cells and go through the protocol described below and usually resulted in ~250-500ng DNA for library preparation (**Fig. 1a**). 10% or 20% of these DNA were used to generate library (0.1 or 0.2 million cells per library). In the second scenario (“mini” setting), we started the experiments with lysing 0.1 or 0.2 million cells following the same protocol as described below, except that all steps before library preparation were performed in 25% volume. Because the cell lysis and *HindIII* digestion conditions are different from the published *in situ* Hi-C protocol. We have made modifications in order to ensure nuclei integrity during ligation.

###### 3.2.1. *Cell lysis, HindIII digestion, and in situ ligation*

Cell pellet from ~1 million cells was lysed in 1ml cell lysis buffer (10mM Tris-Cl, pH7.5, 10mM NaCl, 0.2% NP-40, 1X proteinase inhibitor cocktail (Roche)) before incubating on ice for 15 minutes. If there is cell clump in the tube, we dounce the cells for 10 times every cycle for 3 cycles, with one-minute on ice between each cycle. After douncing, the nuclei were put on ice for another 5 minutes and then pelleted by centrifuging (2,500g for 5 minutes at 4ºC). The pellets were washed once in 1X Cutsmart buffer (NEB) before resuspension in 360ul 1X Cutsmart buffer. After resuspension, 40ul of 1% SDS were added (final 0.1%), and the tubes were incubated at 65ºC for 10 minutes. To quench the SDS, 44ul of 10% Triton X-100 (final 1%) was then added to each tube. For chromatin digestion, 400U *HindIII* (NEB, R3104M 100U/µl) were added to each tube followed by incubation at 37ºC for 4 hours. To ensure efficient digestion, another 400U of *HindIII* were added to each tube again for overnight digestion. On day 2, we digested the nuclei for another 4 hours by adding fresh *HindIII* enzyme (400U). After digestion, the enzyme was inactivated by adding 40ul of 10% SDS (final 1%) to each tube and incubation at 65ºC for 20 minutes. The digested products were then transferred to a new 15ml tube and mixed with 3.06ml 1.15X ligation buffer (75.9mM Tris-HCl, ph7.5, 5.75mM DTT, 5.75mM MgCl_2_ and 1.15mM ATP). 187ul 20% Triton X-100 was added to the mixture and incubated at 37ºC for 1 hour. For ligation, the products were then mixed with 30ul of T4 DNA ligase (Invitrogen, 15224-025, 1U/ul) and incubated at 16ºC overnight. After ligation, the tubes were put in room temperature for 30 minutes and the nuclei were pelleted by centrifuging at 2,500g for 5 minutes. The supernatant was discarded to remove the free DNA and only the nuclei pellets were kept. The nuclei pellet step is skipped in the “dilute” libraries in **Extended Data Table 2**. The nuclei pellets were then resuspended in 3.06ml of 1.15X ligation buffer and mixed with 40ul of 10% SDS and 187ul of 20% Triton X-100 for nuclear lysis.

###### 3.2.2. Reverse crosslinking, DpnII digestion and self-ligation

After nuclear lysis, the mixture was then reverse crosslinked at 65ºC overnight after adding 25ul of 20mg/ml proteinase K. DNA were purified with Phenol: Chloroform: Isoamyl Alcohol (25:24:1, Affymetrix) following standard protocol. ~2-3μg DNA are expected from 1M cells. The DNA was then digested with 50U *DpnII* (NEB, R0543L, 10U/μL) in a total volume of 100uL at 37ºC for 2 hours. After digestion, the enzyme was heat inactivated at 65ºC for 25 minutes. The mixture was first incubated with 0.5 volume of PCRClean DX beads (Aline Biosciences) at room temperature for 10 minutes before harvesting the supernatant according to vendor’s protocol. The supernatant was then incubated with 2 volumes of PCRClean DX beads at room temperature for 10 minutes. DNA on the beads was then harvested in 300ul nuclease free water. The two-step bead purification results in DNA with a size range ~100-1,000bp. The DNA products were then mixed with 200ul of 5X ligation buffer, 5U T4 DNA ligase (Invitrogen, 15224-025, 1U/ul) and water to a total volume of 1ml. Self-ligation was done by incubating the tubes at 16ºC overnight.

###### 3.2.3. Exonuclease digestion and DNA circle re-linearization.

The self-ligated DNA were purified again with Phenol: Chloroform: Isoamyl alcohol and digested with 6U of lambda exonuclease (NEB, M0262S) in 200μL volume at 37ºC for 30 minutes. The exonuclease was then inactivated by incubating at 65ºC for 20 minutes. Resulting DNA were purified with 2 volumes of PCRClean DX beads as described above. For DNA circle re-linearization, bead bound DNA were eluted and digested with 20U *HindIII* again at 37ºC for 2 hours in 150μL volume. The *HindIII* enzyme was inactivated at 65ºC for 20 minutes, and the DNA was purified with 2 volume PCRClean DX beads for another time as described above. In the end, bead-bound DNA was eluted in 50ul nuclease free water. From 1M cells, we expect 250-500ng DNA in the end.

###### 3.2.4. Library preparation.

We took ~10-20% of re-linearized DNA (~50ng) for library generation following Illumina TruSeq protocol. Briefly, the DNA was first end repaired using End-it kit (Epicentre, #ER0720). The end-repaired DNA was then A tailed with Klenow fragment (3′–5′ exo–; NEB, #M0212L) and purified with PCRClean DX beads. Bead bound DNA were eluted in 20μL water and then reduced to 4μL using Speedvac at 50ºC. The 4ul DNA product was mixed with 5ul of 2X quick ligase buffer, 1ul of 1:10 diluted annealed adapter and 0.5ul of Quick DNA T4 ligase (NEB, #M2200L). The ligation was done by incubating at room temperature for 15 minutes and the enzyme was then inactivated by incubating at 65ºC for 10 minutes. DNA was then purified with 1.8 volume of DX beads as described above. Elution was done in 14ul nuclease free water. After checking eHi-C libraries quality, we only needed to sequence less than 1 million reads on MiSeq (Illumina). Because the proportion of PCR duplicates from low-depth sequencing is very low, we used TruSeq indexed adapters (Illumina) without UMI barcode. To deep sequence an eHi-C library, we used custom TruSeq adapter in which the index is replaced by a 6 base random sequence. The custom adapter was generated by annealing the following two oligos:

Universal oligo –

AATGATACGGCGACCACCGAGATCTACACTCTTTCCCTACACGACGCTCTTCCGATC*T

UMI oligo --

/5Phos/GATCGGAAGAGCACACGTCTGAACTCCAGTCACNNNNNNATCTCGTATGCCGTCTTCTGCTT*G

###### 3.2.5. PCR amplification of DNA libraries.

To amplify the DNA libraries, we mixed 13ul adapter ligated DNA with 1ul of 20uM oligo C (AATGATACGGCGACCACCGAGATCTACAC), 1ul of 20uM oligo D (CAAGCAGAAGACGGCATACGAGAT) and 15ul of 2X KAPA HiFi Hotstart ready mix (Kapa Biosystems, #KK2602). And the PCR amplification was done as follows: denature at 98ºC for 45 seconds, cycle at 98ºC for 15 seconds, 60ºC for 30 seconds, 72ºC for 30 seconds, and we did 5 cycles at first for estimating the total cycle number needed, and then further extension at 72ºC for 5 minutes. The products were then purified using 1.8 volume of PCRClean DX beads (Aline Biosciences, #C-1003-50) to remove primer contamination as described above. And the DNA was eluted in 20ul nuclease free water. And library quantification was done following the protocol of Illumina library quantification kit (KAPA Biosystems, KK4824). PCR was done again in 50μL volume for a target final concentration ~20-40nM (usually ~3-4 additional cycles). The generated libraries were then subjected to sequencing.

##### 3.3. The overview of eHi-C performance

We tested eHi-C in low-input setting with ~0.1-0.2 million human primary lung fibroblast IMR90 cells and used low- or high-depth sequencing to evaluate the library quality (**Extended Data Table 2**). As expected, averagely 95% of eHi-C reads begin with digested *HindIII* restrictive sequence AGCTT, indicating that nearly all reads are from re-linearized *HindIII*-digestible DNA circles. When one eHi-C library from 0.1 million cells is deep-sequenced to 150 million mapped read pairs, the percentage of PCR duplicates is lower than the published IMR90 Hi-C libraries prepared with 100 times more (10 million) cells^1^ (**Extended Data Table 2-3**), indicating a significantly improved library complexity.

We also compared the sources of errors in Hi-C and eHi-C libraries^1^^,^^5^. In conventional Hi-C, read pairs falling into the same *HindIII* fragments are considered invalid, and the major type of invalid reads are “dangling reads” originated from non-ligation DNA. In contrast, the only type of invalid pairs from eHi-C are self-circles, all the other types of invalid pairs are removed by exonuclease treatment (**Supplementary Fig. 1e**).

While eHi-C avoids several types of common false reads found in Hi-C, it has a drawback of getting false reads from undigested *HindIII* sites, which can be computationally filtered as back-to-back read pairs next to the same restrictive sites (**Extended Data Fig. 1e-f**). After data filtering, we found that the yield of *cis*-contacts from eHi-C libraries, especially the ones prepared with *in situ* ligation procedure, is better than most of the published *HindIII*-based Hi-C libraries prepared with ~10-25 million cells (**Extended Data Fig. 1g-h, Extended Data Table 2-3**). Importantly, the contact heatmaps from Hi-C and eHi-C data are identical showing the same component A/B^11^ and TAD^12^ structures (**Extended Data** **Fig. 1i-j**). All these results demonstrated that eHi-C is a reliable alternative to Hi-C and can correctly identify 3D genome features from small cell populations.

##### 3.4. Easy Hi-C data pre-processing for QC and performance analysis

Note: The data filtering step of deep Hi-C and eHi-C data for fragment level analysis is slightly different from the performance analysis here. Please refer to **section 5.2** for details.

###### 3.4.1. Alignment and removing PCR duplications.

Published IMR90 Hi-C data are used in this study to compare with eHi-C. The accession numbers of Hi-C data are listed in **Extended Data Table 3**. All the sequencing data are mapped to human reference genome hg19 using Bowtie. For Hi-C, the two ends of paired-end (PE) reads were mapped independently using the first 36 bases of each read. PCR duplications were defined as PE reads with both ends mapped to the same locations. For eHi-C, because nearly all the mappable reads start with HindIII sequence AGCTT, we trimmed the first 5 bases from every read, took the next 36 bases, and added the 6-base sequence AAGCTT to the 5’ of every read before mapping using the whole 42 bases. Some MiSeq runs were performed with reads shorter than 41 bases, and the full-length reads will be used in those cases. After mapping, we further filtered the reads requiring the positions of both ends to be exactly at the HindIII cutting sites. The deep sequenced eHi-C libraries were prepared with UMI adapter, PCR duplications were defined as identical PE reads also with the same UMI barcode. The eHi-C libraries sequenced on MiSeq were not intended for deep sequencing and therefore were prepared without UMI barcode. We assume no PCR duplication in MiSeq libraries because the sequencing depth is very low.

###### 3.4.2. Conventional Hi-C data filtering and QC analysis

After removing PCR duplications, we analyzed the library quality by classifying the reads into different categories. In both Hi-C and eHi-C, the percentage of trans- contacts can be easily calculated by counting the number of reads with two ends on different chromosomes (listed in **Extended Data Table 2-3**). For cis- reads in Hi-C data, we first discard the reads with both ends mapped to the same HindIII fragments as invalid pairs. Dangling ends are defined as “inward” pairs among the invalid pairs (**Supplementary Fig. 1e**) and the percentages are listed in **Extended Data Table 3**. The rest of the invalid pairs are classified into “other false” category.

All rest read pairs represent two different HindIII fragments in *cis*. Since cut-and-ligation events are expected to generate reads within 500bp upstream of HindIII cutting sites due to the size selection (“+” strand reads should be within 500bp upstream of a HindIII site, and “-“ strand reads should be within 500bp downstream a HindIII site), we only keep read pairs with both ends satisfying this criteria. The other pairs are also classified into “other false” category in **Extended Data Table 3**. We next split all the remaining reads into three classes based on their strand orientations (“same-strand”, “inward”, or “outward”) (**Supplementary Fig. 1e**). We have previously shown that although theoretically “same-strand” reads should be twice as many as “inward” or “outward” reads, in reality more “inward” or “outward” reads can be observed due to incomplete digestion of chromatin^1^. We therefore estimate the total number of real cis-contact as twice the number of valid “same-strand” pairs (**Extended Data Table 3**).

###### 3.4.3. eHi-C data filtering and QC analysis

For eHi-C library, the only type of invalid cis- pairs are self-circles with two ends within the same HindIII fragment facing each other (**Supplementary Fig. 1e**). Similar to Hi-C, we also computed the total number of real *cis*-contact as twice the number of valid “same-strand” pairs. Reads from undigested HindIII sites are back-to-back read pairs next to the same HindIII sites facing away from each other (**Supplementary Fig. 1f**).

##### 3.5. Compare the bias structure of Hi-C and eHi-C

*Summary*: We analyzed the intrinsic biases that may affect the eHi-C experimental procedure. As expected, both Hi-C and eHi-C show a decay of contact frequency with increasing distance (**Supplementary Fig. 2a**). The contact frequencies involving very small *HindIII* restriction fragments (< 200bp) are low in both Hi-C and eHi-C libraries, because the small fragments are less likely to be sheared or digested (see **Supplementary Methods**), or due to the spatial hindrance for small fragments to ligate (**Supplementary Fig. 2b**)^13^. The eHi-C has an overall better performance capturing ligation between small-sized (~200bp-1kb) fragments (**Supplementary Fig. 2b-d**), presumably because *DpnII* can digest small *HindIII* fragments effectively as long as the restrictive sites are present. Furthermore, the profile of distance decay at short range is affected by the length of the two HindIII fragments (**Supplementary Fig. 2c-d**), indicating an interaction between the three parameters. Intriguingly, the GC-bias profile in eHi-C library is opposite to what was observed for conventional Hi-C^13^ (**Supplementary Fig. 2e**). We speculate that this might be because both ends of the eHi-C library start with a fixed *HindIII* restrictive sequence (AGCTT). Therefore, the GC-bias in eHi-C reflects the efficiency of DNA polymerase elongation after it has already gone through first few bases during PCR amplification or sequencing. Finally, as expected, eHi-C libraries are also constrained by the size selection of ligation products (**Supplementary Fig. 2f**). These analyses provide a basis for the eHi-C data normalization and computational inference of DNA contacts.

*Methods*: To plot the decay of contact with distance (**Supplementary Fig. 2a**), we only used “same-strand” *cis*- contact reads. For any given distance $L$, we found all HindIII fragment pairs with gap distance between $0.9*L$ and $1.1*L$, and computed the average contact frequency among them. We normalized these numbers by dividing them by the average contacts from all the intra-chromosome HindIII fragment pairs. For length bias (**Supplementary Fig. 2b**), we divided all the HindIII fragments into 40 equal-sized groups and computed the average *trans-* contact frequency for each pair of groups, and enrichment values were calculated by normalizing to the global average. Similarly, we also plotted the GC bias (**Supplementary Fig. 2e**) using *trans-* data. We divided all the HindIII ends into 20 equal-sized groups by GC content. For Hi-C, the GC content was computed using the 200bp near each HindIII end. For eHi-C, the GC content was computed for the region between a HindIII end and the nearest DpnII site.

#### 4. Compartment level Hi-C or eHi-C data analysis

##### 4.1. Calling compartments from Hi-C or eHi-C data

We performed compartment level analysis following the method described previously^11^. We divide the genome into 250kb bins and generate the contact matrices between bins for each chromosome. We next normalize the matrix $M$ by genome distance. For every interaction value $x_{i,j}$ ($i$ is the row number, $j$ is the column number) in matrix$M$, let the distance for this interaction be$L_{|j-i|}$, and we calculated the average of all interaction values with the same distance$avg\left( \sum_{L_{\left| j-i \right|}} x \right).$ Thus, the normalized matrix$M^{'}$ is:

$x_{i,j}^{'}= {x_{i,j}}/{avg\left( \sum_{L_{\left| j-i \right|}} x \right)}$.

We next generated the correlation matrix $M^{''}=cor(M^{'})$, in which each element $x_{i,j}^{''}$ is the Pearson’s correlation coefficient for two vectors $x_{i,*}^{'}$and $x_{j,*}^{'}$from $M^{'}$, representing the similarity of two bins’ interaction pattern. The principal component analysis on the correlation matrix then assigns the genome into two compartments depending on whether the PC1 of a bin is negative or positive value. We used the H3K4me3 data in each cell type to determine the compartment A and B (More H3K4me3 peaks: compartment A; fewer H3K4me3 peaks: compartment B). Since H3K4me3 data for the fetal CP and GZ are not available, we used the H3K4me3 data from fetal cortex instead.

##### 4.2. Identifying regions with different neighborhood profiles, or differentially compartmentalized regions (DCRs)

To identify DCRs, we defined a similarity score to describe how similar the interaction patterns of the same bin $i$ between cell type A and cell type B are. Only *cis* data are used.

$$s_{i}^{A,B}=cor(x_{i,A}^{''},x_{i,B}^{''} )$$

Because $s_{i}^{A,B}\in[-1,1]$, we first do data transformation $x=(s+1)/2$, then used Beta distribution to model the similarity score.

$f\left( x \right)= \frac{x^{\alpha-1}{(1-x)}^{\beta-1}}{B(\alpha,\beta)}$ $0\leq x\leq1;\alpha,\beta>0$

$B\left( \alpha,\beta\right)$is the Beta function; $\alpha,\beta$ are the shape parameter to describe the Beta distribution. We computed the p-value to pick up the bins with significantly different interaction patterns between two cell types.

$p=Prob\left( X< x \right|\alpha, \beta)$.

#### 5. Fragment-resolution Hi-C or eHi-C data analysis

##### 5.1. Determine the sequencing depth required for fragment-level analysis

The highest possible resolution of Hi-C analysis is between individual restrictive fragments (fragment level). Depending on the restrictive enzyme used, the theoretically best resolution for Hi-C is 2 kb (with 6-cutter, *e.g.* *HindIII*) or 128 bp (with 4-cutter, *e.g. DpnII*). However, the feasibility to achieve high resolution also depends on the sequencing depth. Here we propose a rule-of-thumb to determine the sequencing depth requirement for high-resolution analysis.

There are ~350,000 *HindIII* fragments in human genome (we merge fragments < 5 kb in to neighboring fragments, ~7kb resolution), and therefore ~65 billion possible fragment pairs. With ~1 billion total contacts, the average reads number of a fragment pair is only 0.015. Therefore, genome-wide fragment level Hi-C analysis is not possible with billion-scale sequencing depth due to the lack of statistical power. On the other hand, within a short range (such as ~1-2 Mb), data density is high enough so that most fragment pairs have non-zero values. According to our experience, the density of *cis-* data is ~20 fold higher than *trans*-; and the *cis*- data density within 2Mb is ~30 fold higher than over 2Mb (**Extended Data Table 4**).

Analyses of the ~350,000 *HindIII* fragments in human genome has an average resolution of 5-10 kb. There are ~3.5 billion possible fragment pairs in *cis*, and ~100 million possible pairs with the 2 Mb window. ***In order to determine the minimum sequencing depth, we required the average expected frequency to be > 2 between all fragment pairs within 2 Mb.*** The purpose is to prevent too many zeros in the contact matrices. This translates to a requirement of at least 200 million *cis*- contacts within 2 Mb (or mid-range contacts) after data filtering.

It should be noted that the mid-range contacts is not evenly distributed within the 2 Mb window. In the example of GM12878 cells, with the global average value in 2 Mb being 2, the average contact number decreases when the distance increases, *e.g.* 6 (100 kb), 2 (500 kb), 1 (1 Mb), and 0.4 (2 Mb). Therefore, unless ~2~-5 times more data above minimum are generated, we still expect a suboptimal performance for the range between 1 Mb and 2 Mb. In a typical Hi-C experiment, ~40-80% of all *cis*- contacts are within 2 Mb. Therefore, ~300-500 million filtered *cis*- contacts are required for fragment level analysis within 2 Mb. Depending on the *cis-* / *trans-* ratio of the Hi-C experiments, the minimum number of contacts (*cis* and *trans*) after filtering should be ~0.5-1 billion (**Extended Data Table 4**).

The same rule also applies to Hi-C data with 4-cutter, which theoretically may achieve finer resolution. For 1kb resolution within 2Mb window (7-fold finer), a minimum of ~25 billion contacts (0.5 X 7^2^ billion) is required. To our knowledge, the densest published dataset is the *in situ* Hi-C data in GM12878 (4.9 billion total contacts)^10^, which is roughly enough for 1 kb resolution in 1 Mb window, or 2kb resolution in 2Mb window. Taken together, sequencing depth, not the choice of cutter, is the bottleneck for kilobase-scale resolution Hi-C analysis due to the cost-effectiveness limitation of current sequencing technology.

##### 5.2. Hi-C and eHi-C data filtering for fragment level analysis

This step is largely the same as described in **3.4.2-3.4.3** with additional data filtering at the fragment level. Specifically, for Hi-C data, we keep all “same-strand” reads, discard all “inward” data for fragment pairs with the size of gap less than 1kb, and discard all “outward” data for fragment pairs with gap size less than 25kb, as reported previously^1^. For eHi-C, we also keep all “same-strand” reads, but discard all “inward” data for fragment pairs with the size of gap less than 25kb, and discard all “outward” data for fragment pairs with gap size less than 1kb. We used different rules in eHi-C because strand-directions in eHi-C and Hi-C are opposite (**Supplementary Fig. 1e**). For example, undigested *HindIII* sites cause “inward” reads in Hi-C but “outward” reads in eHi-C.

##### 5.3. Fragment-resolution Hi-C analysis to identify cis- looping interactions

This part describes the method to analyze *cis-* Hi-C data within 2Mb window at fragment resolution. The eHi-C data analysis follows the same idea but is slightly different (**section 5.4**). We have previously reported a fragment level Hi-C data analysis to model the significance of ligation product enrichment between any pairs of HindIII fragments^1^ based on a previous systematic study of biases in Hi-C data.^13^ The pipeline includes a normalization step that estimates expected frequencies between any two fragments after correcting several explicit Hi-C biases, a negative binomial model to assess the statistical significance, and a peak-calling step identifying significant fragment pairs as DNA loops. In this study, we included an additional factor in the normalization step to correct an implicit “visibility” factor, which can correct unknown sources of biases and improve the normalization results (**5.3.1-5.3.3**). We still used a negative-binomial model to compute the p values for each fragment pairs (**5.3.4**). Finally, we devised a balanced loop-calling method which reduces biases by considering both enrichment ratio and p-values (**5.3.5**).

###### 5.3.1. A model to estimate expected frequencies between two HindIII fragments

In Hi-C, every *HindIII* fragment has two ends that can form ligation junction with other fragments, and the two ends of the same fragment may have different local mappability and GC content values. We therefore analyze the two ends of a fragment differently. Note that if two ends $i$ and $j$ belong to the same *HindIII* fragments, they will have the same length and distance parameters, but different GC-content and mappability parameters. The goal of this normalization step is to estimate $\mu_{i,j}$, the expected number of reads between two ends $i$ and $j$. We have developed a new model to compute $\mu_{i,j}$, which corrects both known and unknown sources of Hi-C biases.

$$\mu_{i,j}=m_{i}*m_{j}*F_{i,j}^{gc}*L_{i,j}*V_{i}*V_{j}$$

In this equation, $m_{i}$ and $m_{j}$ are the mappability of end $i$ and end $j$. $F_{i,j}^{gc}$ is a correction factor for GC-bias. $L_{i,j}$ is the expected *cis-*contact frequency between end $i$ and end $j$ if both ends are 100% mappable. The explicit correction of factors $m_{i}$, $m_{j}$, $F_{i,j}^{gc}$ and $L_{i,j}$ are the same as described previously^1^. We introduce two additional factors, $V_{i}$ and $V_{j}$, for the “visibility” of the two ends. The correction of visibility corrects unknown sources of biases implicitly.

To further explain this model: (1) Mappability bias originates from the sequence alignment step, the mappability of two fragments are independent from each other, and independent from all other sources of biases. (2) The computation of $L_{i,j}$ corrects biases from distance and the length of the two fragments. These three parameters are interacting factors affecting the proximity ligation in Hi-C protocol, which need to be corrected using the joint function (**Supplementary Fig. 2c**). (3) The GC contents of the two fragments are likely interacting factors, which also need to be corrected using joint function ($F_{i,j}^{gc}$). On the other hand, as Yaffe et al. pointed out^13^, the GC content of the two ends introduce bias mainly through affecting PCR efficiency during the library preparation, which is an independent step from the proximity ligation in Hi-C protocol. Therefore, we assume that the correcting factors in $F_{i,j}^{gc}$ and $L_{i,j}$ are independent from each other. (4) After correcting the aforementioned explicit biases, we assume that the implicit visibility factors $V_{i}$ and $V_{j}$ are additional independent sources of biases that are also independent from each other. Biologically, $V_{i}$ and $V_{j}$ may be understood as the concentration of the two ends in the Hi-C protocol. For example, *HindIII* sites at open chromatin are more likely to be digested by restrictive enzyme. Another possibility is that there might be unannotated copy number variants for a fragment. (5) Theoretically, the mappability biases can be corrected during visibility correction. An alternative model is: $\mu_{i,j}=F_{i,j}^{gc}*L_{i,j}*V_{i}*V_{j}$, in which $V_{i}$ and $V_{j}$ incorporate $m_{i}$ and $m_{j}$ as implicit bias sources. Here, we still correct mappability explicitly even though the difference between two models are trivial.

###### 5.3.2. Correcting known sources of biases with explicit approach

This step is largely the same as described previously^1^. Firstly, local fragment mappability is expected to have a linear effect on the expected ligation frequency^13^. We used a real value $m_{i}$ (ranges from 0 to 1) to represent the mappability of fragment $i$ at forward or reverse strand (representing the two ends of the restriction fragment). To calculate the mappability of a fragment, we generated 36-base pseudo-reads every 9 bases within 500 bases from the end of fragment $i$, and then use bowtie to determine the fraction of uniquely mapped pseudo-reads.

It has been reported that ligation product processing and sequencing may be biased due to local GC content 200bp near restrictive cutting site^13^. We therefore corrected this bias by adjusting $\mu_{i,j}$ according to the local GC content of the two fragments. We split all the ends in to 20 equal-size groups according to their GC contents, and calculated two-dimensional GC-bias matrices (for the fold enrichment of average read counts between groups) using *trans*- Hi-C data. We corrected GC-bias in *cis*- Hi-C data with the GC-bias matrices.

To correct biases from end size and distance, we sorted all the ends based on the length of their corresponding *HindIII* fragments, and divided all the ends into 20 equal size groups. We define the distance between two ends being the size of the gap between their corresponding fragments, and set up 400 groups for distance within the range ~0-2Mb, or one group per 5kb distance. Therefore, group 1 has gap size ~0-5kb; group 2 has gap size ~5-10kb; group 3 is ~10-15kb, *etc*. Because when we do the data filtering, we remove “inward” reads between end pairs with gap size < 1kb, in order to be consistent, we further split group 1 into two new groups with gap size ~0-1kb and gap size ~1-5kb. Therefore, there are total 401 groups based on distance.

Let $G_{i}^{len}$ and $G_{j}^{len}$ be the group assignment of ends $i$ and $j$ based on length; $G_{i,j}^{dist}$ be the group assignment for the pair of end $i$ and $j$ based on the distance between the two ends; $G_{i}^{gc}$and $G_{j}^{gc}$ be the group assignment for the ends $i$ and $j$ based on GC content of its two ends; and $x_{i,j}$ be the observed paired-end reads count between ends $i$ and $j$.

We used the following equation to estimate $L_{i,j}$

$$L_{i,j}=(\sum_{k,l} \frac{x_{k,l}}{m_{k}*m_{l}})/(\sum_{k,l} 1)$$

For $\forall\{k,l\}$ satisfying:

${G_{k}^{len}=G}_{i}^{len}, {G_{l}^{len}=G}_{j}^{len}, {G_{k,l}^{dist}=G}_{i,j}^{dist}, and chr\left( k \right)=chr\left( l \right), m_{k}>0.2,m_{l}>0.2$ (Minimum mappability values of 0.2 are set to avoid division-by-zero errors). Therefore, this is a joint function of two size parameters and the distance parameter. There are 16,040 groups in total with different combination of fragment size and distance.

$F_{i,j}^{gc}$ is a correction factor for GC-bias, which can be computed with *trans-* Hi-C data using the following equation:

$$F_{i,j}^{gc}=\frac{(\sum_{k,l} \frac{x_{k,l}}{m_{k}*m_{l}})/(\sum_{k,l} 1)}{(\sum_{u,v} \frac{x_{u,v}}{m_{u}*m_{v}})/(\sum_{u,v} 1)}$$

For $\forall\{k,l\}$ satisfying:

$$G_{k}^{gc}=G_{i}^{gc},G_{l}^{gc}=G_{j}^{gc}, chr\left( k \right)\neq chr\left( l \right),m_{k}>0.2, m_{l}>0.2$$

And for $\forall\{u,v\}$ satisfying:

$$chr\left( k \right)\neq chr\left( l \right), m_{u}>0.2,m_{v}>0.2$$

In this equation, the denominator is the average frequency of all *trans*- fragment pairs; and the numerator is the average frequency of a subset of those fragment pairs after stratifying GC-content. Note $chr\left( i \right)$ is the chromosome where fragment $i$ is in. The same equation was also used to correct *trans-* Hi-C data except requiring $chr\left( k \right)\neq chr\left( l \right)$.

###### 5.3.3. Implicitly correcting unknown biases hidden in “visibility”

We computed visibility for every *HindIII* end using *trans-* Hi-C data. Since known sources of biases are corrected explicitly for *cis*- data normalization in 2Mb, we need to remove the known biases while calculating visibility factor. The following equation is used to compute $V_{i}$:

$$V_{i}=\frac{\sum_{k} \frac{x_{i,k}}{m_{i}*m_{k}*F_{i,k}^{gc}*F_{i,k}^{len}}}{(\sum_{u,v} \frac{x_{i,k}}{m_{u}*m_{v}*F_{u,v}^{gc}*F_{u,v}^{len}})/(\sum_{u} 1)}$$

For $\forall\{k\}$ satisfying: $chr\left( k \right)\neq chr\left( i \right), m_{k}>0.2, m_{i}>0.2$;

And for $\forall\{u,v\}$ satisfying: $chr\left( u \right)\neq chr\left( v \right), m_{u}>0.2,m_{v}>0.2$.

This equation counts the total *trans-* reads for a *HindIII* end (after correcting the known bias including mappability, GC content and fragment length), and computes its correction factor by dividing with the average count of all the ends. $F^{gc}$ is the same correction factor for the GC-bias computed in **5.3**.**2**. $F^{len}$ is a correction factor for *HindIII* fragment length calculated with *trans*- data:

$$F_{i,j}^{len}=\frac{(\sum_{k,l} \frac{x_{k,l}}{m_{k}*m_{l}})/(\sum_{k,l} 1)}{(\sum_{u,v} \frac{x_{u,v}}{m_{u}*m_{v}})/(\sum_{u,v} 1)}$$

For $\forall\{k,l\}$ satisfying: $G_{k}^{len}=G_{i}^{len},G_{l}^{len}=G_{j}^{len}, chr\left( k \right)\neq chr\left( l \right),m_{k}>0.2, m_{l}>0.2$;

And for $\forall\{u,v\}$ satisfying: $chr\left( u \right)\neq chr\left( v \right), m_{u}>0.2,m_{v}>0.2$.

Finally, after estimating the $\mu$ values for all the ends, we can sum all end-specific values to obtain expected Hi-C read counts for the whole fragment. The fragment-specific $\mu$ values are the Poisson parameter between fragments.

###### 5.3.4. Use negative binomial model to compute the significance of pixels

Two classes of loop calling methods, looking for either “global enrichment” or “local enrichment”, have been developed to identify *cis*- chromatin interactions from Hi-C data. However, the identified loops from these methods only partially overlapped^14^. This is mainly due to the interference from high background signal at short range, reflected by the strong signal along the diagonal in raw contact matrices. “Global enrichment” methods are highly sensitive to Hi-C data normalization because under- or over- correction of Hi-C biases will lead to a large number of false positives or false negatives. On the other hand, the alternative “local enrichment” performs better identifying discrete peak summits with low surrounding signal, but loses its power when surrounding background signal is high, such as at short-range.

We have previously shown that the Hi-C reads count $X_{i,j}$ between two fragments $i$ and $j$ can be modeled by negative binomial distribution^1^:

$$X_{i,j}\sim NB(r_{i,j}=\frac{\mu_{i,j}}{\beta-1}, p=\frac{\beta-1}{\beta})$$

This distribution has mean $\mu_{i,j}$ and variance $\beta*\mu_{i,j}$, in which $\beta$ is a constant number. To estimate $\beta$, we first selected 20 $\mu$ values spanning the range of all $\mu_{i,j}$, then we for each of the selected 20 $\mu$ value, we took all pairs with expected values between $0.99*\mu$ and $1.01*\mu$ (this typically includes at least 100,000 fragment pairs), and then plotted the variance within each group against their expected reads count. Therefore, $\beta$ is the slope value between variance and mean estimated from linear regression analysis. For each dataset, $\beta$ needs to be re-estimated. We can therefore calculate p-value using negative binomial distribution for any pair of fragments $p_{i,j}=P\left( X_{i,j}\geq x_{i,j} | \mu_{i,j}, \beta\right)$ reflecting the significance of enrichment. Importantly, negative binomial distribution has additive properties when $p$ is constant: read counts between any two groups of fragments can be modeled by $X_{i\in I,j\in J}\sim NB(r_{i\in I,j\in J}, p)$, in which $I$ and $J$ are two disjoint subsets of restriction fragments, and $r_{i\in I,j\in J}=\sum_{i\in I, j\in J} r_{i,j}=\frac{1}{\beta-1}\sum_{i\in I, j\in J} \mu_{i,j}$ is dependent on the sum of expected random collision frequency between two groups of fragments. This additive property is convenient because we can quickly determine the parameters for statistical tests when neighboring *HindIII* fragments are merged. Using this model, we can calculate the p value for any fragment pair $i$ and $j$: $p_{i,j}=Prob\left( X_{i,j}> x_{i,j} \right|\mu_{i,j}, \beta)$.

###### 5.3.5. Looping calling and visualization in ratio heatmaps

We computed the enrichment ratio of each pixel and used the value to draw the ratio heatmaps.

$e_{i,j}=(x_{i,j}+d)/{(\mu}_{i,j}+d)$,

In this equation, $d$ is a dummy number to prevent large ratio when $\mu_{i,j}$ is very small.

The loop calling procedure identifies red pixels as chromatin interactions. Using p-value alone for loop calling is biased toward short-range, because the data density at short-range is high, a pixel may achieve statistical significance even with modest enrichment. It actually makes better sense to call loops using enrichment ratios. However, using a ratio cutoff is biased toward long-range because when $\mu_{i,j}$ is very small due to the low data density, the ratio can be very big but lacks statistical significance. If the dummy number is too small, the ratio heatmaps have many red noisy pixels at long-range.

We devised a method to address this problem by adjusting the dummy number. For any pixel, the ratio decreases with increasing $d$, but its p-value does not change. Therefore, if we use a two-fold cutoff, there will be fewer positive pixels when dummy number is higher. We picked the minimum dummy number so that every pixel passed the two-fold cutoff have p-value < 0.001. The dummy numbers are 6 (H1 hESC), 10 (IMR90, fetal CP, fetal GZ, and adult cortex), 7 (GM12878), 13 (hiPSC, hNPC, hNeuron, and fetal cortex). These dummy numbers are also used to compute the ratios when we draw the ratio heatmaps. In the ratio heatmaps, we used a default color scale so that the pixels with over two-fold enrichment are in brightest red.

##### 5.4. Fragment-resolution eHi-C analysis to identify cis- looping interactions

There are some important differences between Hi-C and eHi-C data normalization. Firstly, eHi-C read from a *HindIII* end is completely predictable (**Supplementary Fig. 1**). Therefore, the mappability of a *HindIII* end is only 0 or 1. We therefore first filtered out data from all the 0 mappability ends. Furthermore, if a *HindIII* fragment does not have *DpnII* sites, it should not generate ligation reads because we used *DpnII* to fragment the DNA. We therefore next removed all the reads from such fragments and excluded these fragments from further analysis. After this additional data filtering, the resulting model does not involve mappability anymore. As discussed in **section 3.5**, eHi-C reads are restricted by the size of DNA circles from the ligation product, we therefore need an additional parameter to model DNA circle size.

$$\mu_{i,j}=F_{i,j}^{gc}*{F_{i,j}^{cir}*L}_{i,j}*V_{i}*V_{j}$$

In this equation, everything else is the same as Hi-C analysis except that $F_{i,j}^{cir}$ is a correction factor for the size of ligation product of two ends. Let ${len}_{i}^{HD}$ be the length form a *HindIII* end $i$ to its nearest upstream *DpnII* site, ${len}_{i,j}^{cir}={len}_{i}^{HD}+{len}_{j}^{HD}$.

The following equations are used for eHi-C analysis:

$L_{i,j}=mean(x_{k,l})$,

For $\forall\{k,l\}$ satisfying: ${G_{k}^{len}=G}_{i}^{len}, {G_{l}^{len}=G}_{j}^{len}, {G_{k,l}^{dist}=G}_{i,j}^{dist}, chr\left( k \right)=chr\left( l \right)$

$$F_{i,j}^{gc}=\frac{mean(x_{k,l})}{mean(x_{u,v})}$$

For $\forall\{k,l\}$ satisfying: $G_{k}^{gc}=G_{i}^{gc},G_{l}^{gc}=G_{j}^{gc}, chr\left( k \right)\neq chr\left( l \right)$, and

for $\forall\{u,v\}$ satisfying: $chr\left( u \right)\neq chr\left( v \right)$

$F_{i,j}^{cir}=\frac{mean(x_{k,l})}{mean(x_{u,v})}$

For $\forall\{k,l\}$ satisfying: ${len}_{k,l}^{cir}={len}_{i,j}^{cir}, chr\left( k \right)\neq chr\left( l \right)$, and

for $\forall\{u,v\}$ satisfying: $chr\left( u \right)\neq chr\left( v \right)$

$$V_{i}=\frac{\sum_{k} \frac{x_{i,k}}{F_{i,k}^{gc}*F_{i,k}^{len}*F_{i,k}^{cir}}}{(\sum_{u,v} \frac{x_{i,k}}{F_{u,v}^{gc}*F_{u,v}^{len}*F_{u,v}^{cir}})/(\sum_{u} 1)}$$

For $\forall\{k\}$ satisfying: $chr\left( k \right)\neq chr\left( i \right)$, and

for $\forall\{u,v\}$ satisfying: $chr\left( u \right)\neq chr\left( v \right)$.

$$F_{i,j}^{len}=\frac{mean(x_{k,l})}{mean(x_{u,v})}$$

For $\forall\{k,l\}$ satisfying: $G_{k}^{len}=G_{i}^{len},G_{l}^{len}=G_{j}^{len}, chr\left( k \right)\neq chr\left( l \right)$;

And for $\forall\{u,v\}$ satisfying: $chr\left( u \right)\neq chr\left( v \right)$.

##### 5.5. Loop calling reproducibility

Assess the reproducibility of our loop calling method requires independent datasets with adequate sequencing depth. As mentioned in **5.1**, we need ~200 million mid-range contacts (within 2Mb) for fragment-level loop calling. Therefore, we performed reproducibility analysis after splitting datasets with ~400 million mid-range contacts or more. To summarize, inadequate sequencing depth and batch variation are the two major causes for lower reproducibility; our peak caller consistently achieves Jaccard Index ~0.3 with 60~150K mid-range (< 2Mb) loop calls at ~10kb resolution, which outperforms all major published methods according to a recent comparative analysis.^14^

###### 5.5.1. Loop call reproducibility in GM12878 cells

The GM12878 Hi-C dataset has 5 biological replicates from two different labs with ~385 million total mid-range contacts (**Extended Data Table 4**). We therefore split the 5 replicates into two subsets with roughly equal mid-range contacts (199M and 187M) and compare the reproducibility of chromatin loop callings (**Extended Data Table 5**). Using the same peak calling method described above, the two subsets identified 65K and 84K chromatin loops with 28K overlapping (*Jaccard index 0.23*) (**Extended Data Figure 8a**). As a comparison, our hNPC and hNeuron datasets has 322 million and 302 million mid-range contacts respectively (**Extended Data Table 4**; each identified over 146K loops with 63K overlapping (*Jaccard index 0.275*) (**Figure 5**) despite being different cell types,

Two main reasons cause the non-overlap subset-specific loops in GM12878 cells. Firstly, our loop caller requires p values < 0.001, and ratio > 2 after dummy number adjustment (see **5.3.5**), but when sequencing depth is not adequate, loops from one subset may not pass significant test due to low read numbers. We found that non-overlap loops from one subset usually still have enrichment signal although they do not pass the cutoffs. For example, among the 56,564 loops identified from subset 2 but not subset 1, in subset 1 data 37,106 (66%) have ratios > 1.5, and 43,515 (77%) have p values < 0.05; only 692 (1.2%) do not have any enrichment signal (**Extended Data Figure 8b**). Due to this reason, we always identify more loops when data from subsets are pooled together; the pooled data identity all the overlapped loops and over 80% of the subset-specific non-overlap loops (**Extended Data Figure 8a**). We concluded that inadequate sequencing depth is a major reason for non-reproduced loops, and therefore always use pooled data when multiple biological replicates are available.

The other cause for the subset-specific loops is the batch variation. The five GM12878 replicates have different QC metrics (*trans*/*cis* ratio, proportion of mid-range contacts), and the variation between different labs may be bigger than replicates from the same lab due to the difference with cell culture, chromatin preparation, or Hi-C protocol. We therefore performed another analysis with fetal brain Hi-C data, which has smaller batch variance and probably better for the reproducibility assessment purpose.

###### 5.5.2. Loop call reproducibility in fetal brain

The fetal brain Hi-C dataset is generated by the same lab with a total of ~471 million mid-range contacts from 6 Hi-C experiments, including 3 cortical plate (CP) and 3 germinal zone (GZ) cortex samples (**Extended Data Table 6**). The sequencing depth and QC metrics of the 6 samples are quite even (**Extended Data Table 6**). Although we treated CP and GZ samples separately in all follow-up analyses, the similarity between the two samples are very high, most likely reflecting the fact that CP and GZ are two spatially close regions of brain cortex. At compartment level, CP and GZ show highest similarity (**Figure 1**). Our method identified 138K and 141K loops from CP and GZ sample, with 71K overlapping (*Jaccard index 0.35*) (**Extended Data Figure 8c**). After pooling CP and GZ data together, we called 244,586 loops covering 99.8% of the overlap loops between CP and GZ, and 78% of non-overlap loops.

Given the high reproducibility between CP and GZ data, we also tried to group this dataset into three subsets (every subset has one CP and one GZ); each subset has 150~160 million mid-range contacts (**Extended Data Table 6**). Again, the three subsets identified similar number of chromatin loops (114K, 116K and 119K), the overlap between any two subsets is 54~58K loops, with pairwise *Jaccard Index between 0.30 and 0.34*. 59~64% of chromatin loops from any subset can be called in at least another subset (**Extended Data Figure 8d**, left panel). Again, the pooled dataset can recover nearly 80% of all loops identified from the subset analysis, including 60~70% of the subset-specific loops (**Extended Data Figure 8d**, right panel).

###### 5.5.3. Reproducible neural chromatin loops among 6 neural samples

Finally, we compared the loops identified from 6 neural samples (hNPC, hNeuron, fetal cortex, adult cortex, CP and GZ), and postulated that a meta-analysis of these heterogeneous samples may improve both sensitivity and accuracy, even though the variation between samples may also reflect the tissue- or cell-type specificity. We identified 165K loops that are observed in at least 2 samples, which are considered credible neural loops (**Extended Data Figure 8h**). As expected, this number is higher than loops identified from any sample alone; averagely ~60% of loops from any single dataset are credible neural loops level.

#### 6.  Other data analysis methods

ChIP-seq: ChIP-seq data were mapped to human reference genome hg19 using Bowtie. The first 36 bases of each read were used for mapping. We only use non-redundant reads to eliminate possible duplicates from biased PCR amplification. We used MACS^15^ with default parameters to call ChIP-seq peaks.

Network analysis: For network analysis of neuron differentiation chromatin loops, we took all fragments containing TSSs, and all fragments containing H3K27ac peaks in hiPSC, hNPC or hNeuron. All chromatin loops in the three cell types are used to construct the network. Each fragment is a node and every chromatin loop is an edge. We built the network with *NetworkX*^16^ and visualized with *Cytoscape*^17^. The network in **Fig. 6c** is drawn using only a portion of top interactions (~800) based on enrichment ratios. The resulting network is divided into hundreds of components and the smallest component is two node and one edge. We defined 603 multi-node components (with at least 5 edges) as candidates of enhancer-promoter aggregates. We call neuron-specific component if the average ratio of all its edges in hNeuron is >1.5 fold higher than the average ratio in hiPSC. 174 components satisfied these criteria.

Gene Ontology analysis: For GO analysis, we used RefSeq genes as the background genes downloaded from UCSC table browser. We downloaded the complete gene sets (function categories) from MSigDB (Molecular Signatures Database, version 5.2) from GSEA website (http://software.broadinstitute.org/gsea). We used one-tailed binomial test to calculate p-values of enrichment of any function categories. We used the R package *qvalue* to estimate q-values and FDR for the p-values. We used a cutoff FDR < 0.05 in the analysis.

GWAS SNP, eQTL, and LD analyses: We compiled lists of GWAS SNPs in neuronal relevant disease and diabetes/obesity relevant disease from the NHGRI-EBI GWAS catalog^18^ (**Supplementary Data 3**). The eQTL data of 44 tissues were downloaded from GTEx portal.^19^ We calculated linkage disequilibrium (LD) for all pairs of genetic variants within 1Mb, among individuals with global Europe ancestry estimate ≥0.8 in TOPMed freeze5b samples. The global ancestry estimates were derived from local ancestry estimates from RFMix^20^ using data from the Human Genome Diversity Project (HGDP)^21^ as the reference panel with seven populations, namely Sub-Saharan Africa, Central and South Asia, East Asia, Native America, Oceania, and West Asia and North Africa (Middle East). Global ancestry for each TOPMed individual is defined as the mean local ancestry across all HGDP SNPs. We defined the LD of a GWAS SNP being the region that every SNP inside has D’ > 0.8 with the lead SNP. Consequently, the median size of LD’s is ~150kb. A bigger LD should be more inclusive with potential causal SNPs, which is probably beneficial for the study of SNPs from highly heterogeneous sources provide by GWAS catalog. Additionally, bigger LD also make it more likely that the defined outside-LD SNP-gene pairs (loop or eQTL) represent distal regulatory relationship.

Predicting GWAS target genes with chromatin loop or eQTL data: For any GWAS lead SNP, we define a loop target gene if its TSS loop to the GWAS LD. Similarly, because eQTL data are in the format of SNP-gene pairs, we also predict a GWAS SNP’s eQTL target gene if the eQTL data link a SNP in the LD to the TSS. Note that in this study, we only focused on predicted genes with TSS outside of the GWAS LD. Additionally, since the GTEx only called cis-eQTLs within 1Mb, we only used chromatin loops in this window for fair comparison.

#### 7. Data availability

Data for eHi-C protocol optimization (in IMR90 cells) are available at NCBI GEO with accession number GSE89324. Raw and/or processed eHi-C and ChIPmentation data in hiPSC, hNPC and hNeuron are available at NCBI GEO with accession number GSE115407. Newly generated Hi-C data in hESCs are also included in GSE115407. ChIP-seq and eHi-C from fetal or adult brain cortex are available at NCBI GEO with accession number GSE116825. This study also re-analyzed published Hi-C data and ChIP-seq data. The accession numbers of raw data are listed in **Supplementary Table 7-8**.

#### 8. CRISPR experiments

Generating doxycycline inducible Cas9 expressing hESC line (DI-Cas9-H9)

The pBlue-AAVS1-Puro-Cas9-M2rtTA-AAVS1 HITI donor plasmid was constructed by ligating the HindIII restricted Puro-Cas9-M2rtTA fragment cut out from the Puro-Cas9-M2rtTA plasmid to the pBlue-AAVS1-AAVS1 vector linearized with HindIII. To construct the Puro-Cas9-M2rtTA plasmid, CAG-M2rtTA-pA sequence was amplified from Neo-M2rtTA plasmid and subcloned into the Puro-Cas9 plasmid linearized with MfeI and MluI. To construct the pBlue-AAVS1-AAVS1 plasmid, a pair of oligos for AAVS1 gRNA targeting sequence (g-AAVS1-F: TCACCAATCCTGTCCCTAGGTTTA; g-AAVS1-R: CTAGGGACAGGATTGGTGACGGTG) were annealed and ligated to the pBlue vector linearized with XhoI and NotI. H9 cell line was maintained on Matrigel (Corning, 354277, Bedford, MA, USA) in mTeSR1 (STEMCELL Technologies, 85850/05850, Vancouver, BC, Canada). Cells were cultured at 37 °C in a humidified atmosphere with 5% CO_2_ in air. hESCs were passaged with TrypLE (Gibco, 12604-021, Grand Island, NY, USA). H9 cells were transfected using electrotransfection (1 pulse, 300 V, 4 ms, BTX). A total of 25μg plasmid (donor : Cas9 : gAAVS1RNA = 3 : 3 : 2) was used in each electroporation. Around 4~9 million cells were resuspended with 500μL PBS in 0.4 cm cuvette. Two days later, 0.5μg/mL puromycin was used to treat cells for 3 days. Cells were allowed to grow visible colonies for about 10 days, and then the colonies were picked into 96-well plate. Colonies were expanded and identified by PCR and sequencing (5-F: GGTTAATGTGGCTCTGGTT; 5-R: CTTGTACTCGGTCATCTCG; 3-F: TGACGGTTCACTAAACGAG; 3-R: AGAGGTTCTGGCAAGGAG).

Deleting CTCF sites in ESCs and NPCs with sgRNAs-CARGO

We made CARGO^22^ constructs whenever we need to transfect multiple sgRNAs into the same cell. With CARGO system, we could assemble 4-10 sgRNAs simultaneously into one plasmid following the protocol described by Gu *et al*.^22^ The CARGO plasmids are gifts from the laboratory of Joanna Wysocka. All sgRNAs were designed on CCTop-CRISPR/Cas9 target online predictor (https://crispr.cos.uni-heidelberg.de/) and manually picked. For CARGO, *(n+1)* pairs of oligos are necessary to assemble *n* sgRNAs. The CARGO oligo sequences are listed in **Extended Data Table 9**. We deleted three CTCF-containing regions at *CACNA1C* locus (C1~C3). Successful deletion was verified with PCR. The primer used for C1 (Product length wt: 616 bp, del: 471-518 bp; fwd: ACAGGATGCTATGGGACACC; rev: AGGGAGGAGGAAGAAATGGA); C2 (Product length wt: 786 bp, del: 531-603 bp; fwd: CCTGGGGTGTTGAGAGAGAA; rev: ATTCACCCAAAAGGCTTCCT); C3 (Product length wt: 9,358 bp, del: 550-600 bp; fwd: TGAGCCCAAAGGCACTAGAC; rev: TACCCAGAACAGGCACTTCC).

DI-Cas9-H9 cells were maintained in mTeSR1 medium (STEMCELL technologies, #85850) on matrigel. Cells were detached and suspended to single cells by Accutase (Fisher, #A1110501). CARGO vector transfection was done following the manufacturer’s instruction of Amaxa 2b nucleofector, using Kit 1 (Lonza, Human stem cell nucleofector Kit 1, #VPH-5012) and program B-16. After 24 hours recovery, cells were treated with 1µg/mL of Doxycycline to induce Cas9 expression for 48 hours before harvesting. The hNPCs were differentiated as described above and seeded at 170k cells per cm^2^. Transfection was done following the manufacturer’s instruction of Amaxa 4D nucleofector. Briefly, cells were treated with Accutase to make single cell suspension and then pelleted at 110g for 5min. P3 primary cell 4D-nucleofector X kit L (Lonza, #V4XP-3024) was applied combining program CU-133. After 24 hours recovery, cells were treated with 1µg/mL of Doxycycline to induce Cas9 expression for 48 hours before harvesting for DNA and RNA samples.

Construct dCas9-KRAB-puro for CRISPRi assay

EF1-dcas9-KRAB was PCR amplified from Lenti-dCas9-KRAB-blast (Addgene, #89564) with primers (F: CCTTTTGCTCACATGTGCTAGCTGCAAAGATGGATAAAG, R: AACTTTGCGTTTCTTTTTCGGAACTGATGATTTGAT); T2A-puro was PCR amplified from the LentiCRISPRv2 plasmid (Addgene, #98654) (F: AAGAAACGCAAAGTTGGATCCGGCGCAACAAACTTC, R: CGAGCTCTAGGAATTCTCAGGCACCGGGCTTGCG). The two PCR products were assembled into px332-original plasmid (gifts from the laboratory of Joanna Wysocka^22^) between PciI and EcoRI sites by In-Fusion HD cloning (TAKARA, #639648).

CRISPRi enhancer inhibition in GM12878 cells with sgRNAs-CARGO

We constructed CARGO vectors containing multiple sgRNAs as described above. GM12878 cells were maintained in RPMI1640 with 15% FBS. GM12878 cells were seeded in fresh medium at 350k cells per ml the day before nucleofection. 4 million cells were used for each nucleofection. First, cells were pelleted at 90g for 5min and then resuspended in 100ul of nucleofection reagent (SF cell line 4D-Nucleofector X kit, Lonza, #V4XC-2024) together with 5-7ug designed plasmids. The nucleofection was done on a 4D lonza nucleofector using program CM-137. Puromycin selection was done at 3µg/mL for 48 hours after letting the cell recovering for 24 hours post transfection. Cells were then harvested for RNA extraction, or fixed with 1% formaldehyde. We performed H3K27ac ChIP-qPCR using ChIP-mentation protocol described before. 10% of chromatin was saved as input control. The qPCR and ChIP-qPCR primers used are listed in **Extended Data Table 9**.

3C-qPCR

To confirm whether deletion of CTCF at the CACNA1C locus would lead to loss of chromatin loops, we did 3C assay in hESCs. We followed the protocol as previously described^23^. First, H9 cells harboring CTCF deletion were generated as above by nucleofection and fixed for 3C assay. Briefly, Cells were permeabilized in a lysis buffer (10mM Tris-Cl, pH8.0, 10mM NaCl, 0.2% NP-40 and 1X proteinase inhibitor cocktail), and nuclei were collected by centrifuging at 2500g for 5min. The nuclei were then digested with HindIII-HF (NEB, #R3104M), 400U for 5 million cells at 37 °C overnight. After inactivation of HindIII, the proximity ligation was done with T4 DNA ligase (Invitrogen, #15224-025) at 16°C for overnight. Chromatins were then reverse-linked by proteinase K and purified by phenol: chloroform. Two BAC clones (RP11-265G12 and RP11-698B23) cover the studied region were applied as genomic background control. Equal moles of the DNA from two BACs were mixed together and used to generate the control template following the protocol. Primers designed for 3C-qPCR are listed **Extended Data Table 9**.

#### Supplementary Data 1. The list of reprograming and neuronal differentiation DCRs.

#### Supplementary Data 2. Promoter-enhancer loop network in neurogenesis.

#### Supplementary Data 3. Analyses of neurological GWAS SNPs.

### Extended Data Tables

#### Extended Data Table 1. Brain cortex samples.

Adult cortex tissue

| **ID** | **Ancestry** | **Sex** | **Region** | **Age** | **Post mortem interval** | **Cause of death** |
| --- | --- | --- | --- | --- | --- | --- |
| ME_49 | European | F | anterior temporal cortex | 65yr | 26hrs | Cardiovascular disease |
| ME_50 | European | M | anterior temporal cortex | 37yr | 17hrs | acute hemorrhagic pancreatitis due to choledocholithiasis |
| ME_51 | European | M | anterior temporal cortex | 50yr | 22hrs | Atherosclerotic cardiovascular disease |

Fetal cortex tissue

| **ID** | **Ancestry** | **Sex** | **Region** | **Gestational Age** |
| --- | --- | --- | --- | --- |
| ME_45 | African American | F | Cerebrum | 19 weeks |
| ME_46 | African American | M | Cerebrum | 19 weeks |
| ME_47 | African American | M | Cerebrum | 19 weeks |

#### Extended Data Table 2. QC metrics of eHi-C datasets in IMR90 cells.

| **Library ID** | **eHi-C conditions and cell numbers** | **Uniquely mapped pairs** | **Non-redundant pairs** | **% of PCR dups** | **% of non-redundant pairs** | | | |
| --- | --- | --- | --- | --- | --- | --- | --- | --- |
|  |  |  |  |  | **Self-circle** | **Un-digested HindIII sites** | ***Trans*-contact** | ***Cis*-contact** |
| eHiC1 | Dilut-0.1M^1^ | 2,062,303 | NA | NA | 27.4% | 24.6% | 18.0% | 30.0% |
| eHiC2 | Dilut-0.1M^1^ | 3,021,213 | NA | NA | 28.9% | 26.7% | 16.6% | 27.8% |
| eHiC3 | Dilut-0.1M^2^ | 1,757,103 | NA | NA | 23.7% | 28.7% | 17.7% | 30.0% |
| eHiC4 | Dilut-0.2M^1^ | 93,757 | NA | NA | 24.6% | 28.9% | 17.1% | 29.5% |
| eHiC5 | Dilut-0.2M^1^ | 1,166,089 | NA | NA | 14.4% | 37.0% | 14.4% | 34.2% |
| eHiC6 | Dilut-0.2M^2^ | 907,455 | NA | NA | 34.6% | 15.8% | 24.6% | 24.9% |
| eHiC7 | Insitu-0.1M^1^ | 1,044,280 | NA | NA | 6.1% | 27.3% | 27.5% | 39.1% |
| eHiC8 | Insitu-0.1M^1^ | 905,865 | NA | NA | 7.0% | 23.1% | 31.5% | 38.4% |
| eHiC9 | Insitu-0.1M^2^ | 514,929 | NA | NA | 8.6% | 37.4% | 13.8% | 40.3% |
| eHiC10 | Insitu-0.1M^2^ (deep) | 154,781,478 | 137,347,461 | 11.3% | 13.1% | 33.2% | 14.1% | 39.6% |

^1^ These libraries were prepared with “aliquot” setting starting with 1 million cells. 10% or 20% of the re-linearized junction products were used for library preparation.

^2^ These libraries were prepared with “mini” setting starting with 0.1 or 0.2 million cells (see **Method**).

#### Extended Data Table 3. QC metrics of HindIII-based Hi-C datasets in IMR90 cells.

| **IMR90 Hi-C data**  **(10M cells)** | **Uniquely mapped pairs** | **Non-redundant pairs** | **% of PCR dups** | **% of non-redundant pairs** | | | |
| --- | --- | --- | --- | --- | --- | --- | --- |
|  |  |  |  | **Dangling** | **Other false** | ***Trans*-contact** | ***Cis*-contact** |
| GSM1055800 | 257,464,146 | 185,505,290 | 27.9% | 28.0% | 18.9% | 23.0% | 30.1% |
| GSM1055801 | 275,155,361 | 221,966,824 | 19.3% | 11.8% | 15.0% | 42.6% | 30.6% |
| GSM1154021 | 392,648,087 | 225,360,969 | 42.6% | 49.0% | 16.2% | 22.7% | 12.2% |
| GSM1154022 | 355,763,158 | 163,466,864 | 54.1% | 47.7% | 18.9% | 19.9% | 13.5% |
| GSM1154023 | 154,952,639 | 127,019,197 | 18.0% | 47.4% | 19.4% | 17.4% | 15.8% |
| GSM1154024 | 132,335,077 | 95,775,615 | 27.6% | 36.8% | 17.6% | 24.7% | 20.9% |
| GSM1055802 | 469,381,194 | 202,542,797 | 56.8% | 10.1% | 21.1% | 38.1% | 30.7% |
| GSM1055803 | 381,403,983 | 304,548,370 | 20.2% | 7.2% | 12.9% | 40.8% | 39.1% |
| GSM1154025 | 297,550,433 | 126,467,235 | 57.5% | 26.6% | 14.6% | 32.6% | 26.3% |
| GSM1154026 | 395,911,149 | 159,726,476 | 59.7% | 25.2% | 14.9% | 31.9% | 28.0% |
| GSM1154027 | 104,518,964 | 66,798,106 | 36.1% | 30.9% | 17.3% | 32.2% | 19.6% |
| GSM1154028 | 142,486,775 | 107,980,643 | 24.2% | 27.3% | 15.5% | 29.8% | 27.3% |
| GSM1055804 | 173,877,711 | 149,674,888 | 13.9% | 8.4% | 13.2% | 37.0% | 41.5% |

#### Extended Data Table 4. Summary of Hi-C and easy Hi-C data

|  | **Uniquely mapped pairs** | **Non-redundant** | **# of *trans*-contacts*** | **# of *cis*-contacts*** | **cis- contacts within 2Mb** |
| --- | --- | --- | --- | --- | --- |
| H1 hESC (Hi-C) | 2,667,429,980 | 1,465,769,303 | 205,496,318 | 359,606,047 | 225,077,032 |
| IMR90 (Hi-C) | 3,209,573,496 | 2,030,138,702 | 495,001,524 | 515,971,257 | 221,928,827 |
| GM12878 (Hi-C) | 4,317,877,124 | 3,232,821,732 | 1,096,434,895 | 780,569,204 | 385,152,495 |
| Fetal CP (Hi-C) | 1,446,206,862 | 1,016,026,079 | 450,478,146 | 376,215,499 | 233,082,782 |
| Fetal GZ (Hi-C) | 1,500,816,734 | 1,084,012,945 | 447,691,778 | 389,188,891 | 237,626,547 |
| hiPSC (eHi-C) | 1,594,118,699 | 1,388,527,795 | 200,139,674 | 458,590,294 | 346,564,143 |
| hNPC (eHi-C) | 2,334,966,485 | 1,858,876,027 | 256,835,352 | 500,586,974 | 322,232,584 |
| hNeuron (eHi-C) | 2,421,746,422 | 1,945,779,270 | 222,351,344 | 463,353,815 | 301,875,870 |
| Fetal cortex (eHi-C) | 3,496,691,999 | 2,976,838,880 | 1,282,013,776 | 672,030,711 | 301,592,969 |
| Adult Cortex (eHi-C) | 2,496,925,693 | 2,262,697,027 | 778,496,996 | 723,320,137 | 328,268,152 |
| Colon crypt (eHi-C) | 2,423,323,097 | 1,911,640,712 | 753,963,268 | 342,371,777 | 113,404,289 |
| hiPSC-skin2 (ATCC ACS-1019, eHi-C) | 242,765,498 | 192,508,184 | 24,708,400 | 69,845,399 | 54,904,924 |
| hiPSC-skin3 (ATCC ACS-1011, eHi-C) | 239,804,071 | 192,800,291 | 19,964,703 | 62,508,289 | 49,542,162 |
| hiPSC-BM (ATCC ACS-1026, eHi-C) | 219,158,775 | 189,397,194 | 30,801,221 | 64,771,680 | 49,035,715 |

|  | **% of PCR dups** | **% of non-redundant pairs** | | | |
| --- | --- | --- | --- | --- | --- |
|  |  | **Dangling** | **Other false** | ***Trans*-contact** | ***Cis*-contact** |
| H1 hESC (Hi-C) | 45.0% | 32.2% | 26.8% | 14.9% | 26.1% |
| IMR90 (Hi-C) | 36.7% | 26.4% | 23.8% | 24.4% | 25.4% |
| GM12878 (Hi-C) | 25.1% | 18.1% | 23.9% | 33.9% | 24.1% |
| Fetal CP (Hi-C) | 29.7% | 2.2% | 16.5% | 44.3% | 37.0% |
| Fetal GZ (Hi-C) | 27.8% | 5.6% | 17.2% | 41.3% | 35.9% |

| **Library ID** | **% of PCR dups** | **% of non-redundant pairs** | | | |
| --- | --- | --- | --- | --- | --- |
|  |  | **Self-circle** | **Un-digested HindIII sites** | ***Trans*-contact** | ***Cis*-contact** |
| hiPSC (eHi-C) | 12.9% | 14.9% | 37.7% | 14.4% | 33.0% |
| hNPC (eHi-C) | 20.4% | 6.7% | 52.6% | 13.8% | 26.9% |
| hNeuron (eHi-C) | 19.7% | 7.0% | 57.8% | 11.4% | 23.8% |
| Fetal cortex (eHi-C) | 14.9% | 11.2% | 27.1% | 43.1% | 22.6% |
| Adult Cortex (eHi-C) | 9.4% | 14.1% | 16.8% | 34.4% | 32.0% |
| Colon crypt (eHi-C) | 21.1% | 21.2% | 21.5% | 39.4% | 17.9% |
| hiPSC-skin2 (eHi-C) | 20.7% | 4.8% | 42.9% | 13.1% | 39.2% |
| hiPSC-skin3 (eHi-C) | 19.6% | 4.8% | 49.6% | 10.5% | 35.1% |
| hiPSC-BM (eHi-C) | 13.6% | 7.0% | 39.1% | 16.7% | 37.2% |

#### Extended Data Table 5. Summary of GM12878 replicates and subset splitting

|  | **Uniquely mapped pairs** | **Non-redundant** | **# of *trans*-contacts*** | **# of *cis*-contacts*** | **cis- contacts within 2Mb** |
| --- | --- | --- | --- | --- | --- |
| Rep 1,4,5 (subset 1): 198,509,994 mid-range contacts | | | | | |
| GM12878 rep1 | 269,468,591 | 258,018,532 | 133,780,331 | 66,623,776 | 27,340,514 |
| GM12878 rep4 | 800,714,585 | 625,645,239 | 159,080,816 | 102,123,857 | 45,010,220 |
| GM12878 rep5 | 1,210,464,417 | 1,056,069,146 | 267,517,537 | 228,587,471 | 126,159,260 |
| Rep 2,3 (subset 2): 186,642,501 mid-range contacts | | | | | |
| GM12878 rep2 | 252,427,530 | 237,825,546 | 121,699,948 | 60,326,559 | 24,954,926 |
| GM12878 rep3 | 1,784,802,001 | 1,055,263,269 | 414,356,263 | 322,907,541 | 161,687,575 |

#### Extended Data Table 6. Summary of fetal brain replicates and subset splitting

|  | **Uniquely mapped pairs** | **Non-redundant** | **# of *trans*-contacts*** | **# of *cis*-contacts*** | **cis- contacts within 2Mb** |
| --- | --- | --- | --- | --- | --- |
| CP1 and GZ1 (subset 1): 151,102,172 mid-range contacts | | | | | |
| Fetal CP1 | 532,714,556 | 399,044,047 | 205,237,535 | 136,768,462 | 82,414,240 |
| Fetal GZ1 | 540,875,893 | 413,818,572 | 207,617,983 | 122,071,144 | 68,687,932 |
| CP2 and GZ2 (subset 2): 160,099,765 mid-range contacts | | | | | |
| Fetal CP2 | 521,072,823 | 340,507,492 | 148,709,892 | 125,368,513 | 76,105,460 |
| Fetal GZ2 | 547,006,557 | 357,837,719 | 152,901,659 | 129,669,202 | 83,994,305 |
| CP3 and GZ3 (subset 3): 160,669,832 mid-range contacts | | | | | |
| Fetal CP3 | 392,419,483 | 276,474,540 | 96,948,999 | 114,078,524 | 75,129,170 |
| Fetal GZ3 | 412,934,284 | 312,356,654 | 87,595,874 | 137,448,545 | 85,540,662 |

#### Extended Data Table 7. Accession number of published Hi-C data used in this study

|  | **Replicates** | **Accession Number**** | **Reference** |
| --- | --- | --- | --- |
| H1 hESC | H1 (hESC) Rep1 | GEO:GSM1267196 | Dixon et al.^24^ |
|  | H1 (hESC) Rep2 | GEO:GSM1267197 |  |
|  | H1 (hESC) Rep3 | This study | This study |
|  | H1 (hESC) Rep4 | This study |  |
|  | H1 (hESC) Rep5 | This study |  |
|  | H1 (hESC) Rep6 | This study |  |
| IMR90 | IMR90 Rep1 | GEO:GSM1055800 | Jin et al.^1^ |
|  | IMR90 Rep2 | GEO:GSM1055801 |  |
|  | IMR90 Rep3 | GEO:GSM1154021 |  |
|  | IMR90 Rep4 | GEO:GSM1154022 |  |
|  | IMR90 Rep5 | GEO:GSM1154023 |  |
|  | IMR90 Rep6 | GEO:GSM1154024 |  |
|  | IMR90+TNF Rep1 | GEO:GSM1055802 |  |
|  | IMR90+TNF Rep2 | GEO:GSM1055803 |  |
|  | IMR90+TNF Rep3 | GEO:GSM1154025 |  |
|  | IMR90+TNF Rep4 | GEO:GSM1154026 |  |
|  | IMR90+TNF Rep5 | GEO:GSM1154027 |  |
|  | IMR90+TNF Rep6 | GEO:GSM1154028 |  |
| GM12878 | GM12878 rep1 | GEO:GSM1181867 | Selvaraj et al.^25^ |
|  | GM12878 rep2 | GEO:GSM1181868 |  |
|  | GM12878 rep3 | GEO:GSM1551583 | Rao et al.^10^ |
|  | GM12878 rep4 | GEO:GSM1551584 |  |
|  | GM12878 rep5 | GEO:GSM1551586 |  |
| Fetal CP | CP rep1 | GEO:GSM2054564 | Won et al.^26^ |
|  | CP rep2 | GEO:GSM2054565 |  |
|  | CP rep3 | GEO:GSM2054566 |  |
| Fetal GZ | GZ rep1 | GEO:GSM2054567 |  |
|  | GZ rep2 | GEO:GSM2054568 |  |
|  | GZ rep3 | GEO:GSM2054569 |  |
| Lympho-derived hiPSC | N/A | ArrayExpress:  E-MTAB-6014 | Montefiori et al.^27^ |

#### Extended Data Table 8. Accession numbers of ChIP-seq data used in this study

|  | **H1** | **IMR90** | **GM12878** |
| --- | --- | --- | --- |
| Input | GSE16256 | GSM1055808 | GSM733742 |
| CTCF | GSM733672 | GSM1055825 | GSM733752 |
| H3K4me1 | GSE16256 | GSM1055814 | GSM733772 |
| H3K4me3 | GSE16256 | GSM1055816 | GSM733708 |
| H3K9me3 | GSE16256 | GSE16256 | GSM733664 |
| H3K27ac | GSE16256 | GSM1055818 | GSM733771 |
| H3K27me3 | GSE16256 | GSE16256 | GSM733758 |
| H3K36me3 | GSE16256 | GSM1055820 | GSM733679 |

|  | H3K4me3 | H3K9me3 |
| --- | --- | --- |
| iPSC 6.9 | GSM706075 | GSM706080 |
| iPSC 19.11 | GSM706074 | GSM706079 |
| iPSC 20b | GSM772844 | GSM772842 |
| iPSC 18a | GSM773029 | GSM773023 |
| iPSC 15b | GSM537687 | GSM537691 |
| Skin fibroblast | [GSM817235](http://www.ncbi.nlm.nih.gov/geo/query/acc.cgi?acc=GSM817235) | [GSM817236](http://www.ncbi.nlm.nih.gov/geo/query/acc.cgi?acc=GSM817236) |

#### Extended Data Table 9. Sequences and primers.

| sgRNAs for CTCF deletion at *CACNA1C* locus | |
| --- | --- |
| C1 | TCGCGGTTGCGGCTCTGAAC |
|  | AATCCAACTTTGCGGGACAG |
|  | CACCTTCCAAGTCCTGAAGG |
|  | ACAGGAAAACGGCCTCCTTC |
| C2 | ATTTGTTCAGCCTTGCATCG |
|  | TTTCGAGGCCCTGCTCATCT |
|  | CCTGGTGATGAAAGGTGTAG |
|  | TTAGAGCTTGTGACCTGTTG |
| C3 | CTCCTTCCCTCACAGCTCAT |
|  | TGTGAGGGAAGGAGTGGTTT |
|  | TCTCTTTCTCGACTGCTCTT |
|  | GCAGGACACCAGGAGAGGCA |
| sgRNAs for CRISPRi in GM12878 cells | |
| E1 | CCTGGTCT TGTA ATTCACAG |
|  | ATTACCCC ATGA ATTGAAAG |
|  | GCTGTATA GTGT ATGCTCAG |
|  | CATGGGGA ATTA ATTTATTT |
| E2 | ATACTTTAAGCCTCTCATAA |
|  | GAAAGTCCCAGAATGACTGC |
|  | CACTGGCCAAGAGTTCACAA |
|  | AAAATTACCTAAACCAACTG |
|  | TGAGGAAGCAGGCTTTCACC |
|  | CCAGTGGTTTATCATGCTGA |
|  | CTACAGGTAGTGCCAAATCG |
|  | GTTGTGAAAGTTTCCTTGAG |
| E3 | CCTTTGAACATGCTCTATTT |
|  | ACACATAACTGGATTTTTTG |
|  | GGCCTGCTGAACATTGACAA |
|  | CTTCTCCCTCCAAAGGCATC |
|  | CCATGTAGGTGGGATAGTCT |
|  | GAGGTGACCATCGTACACTG |
|  | AGGAATGGAAGGGTTGGTTG |
|  | ACTGAATCTTGAAACATTAC |
| E4 | AGTGGCGACTTCTTGCAACT |
|  | AACTTTCAGTCGGCTGGAGG |
|  | AGGAGGGGGGAGAAGGGGGG |
|  | GCCCGGGCTCGGCCGCGCAC |
|  | CTAGTGCACAGCCTGCTTTT |
|  | TGGGGGTGTCGGCAGACCAC |
|  | GCAATGAAAAGACATGATTA |
|  | AGAACCATGTACTTGTTCTT |
|  | ACACTGAAATCAGAAACCTG |
|  | GGTCTCCTGCAGTACAAGAG |

| CARGO oligos for CTCF deletion at *CACNA1C* locus | | |
| --- | --- | --- |
| C1 | 1-F | ACCG TCGCGGTT GCGG TTGTCTTCGAAGACAA CATGTG |
|  | 1-R | TAAA CACATGTTGTCTTCGAAGACAACCGCAACCGCGA |
|  | 2-F | ACCG AATCCAAC TTTG TTGTCTTCGAAGACAA GCGG CTCTGAAC G |
|  | 2-R | TAAA CGTTCAGAGCCGCTTGTCTTCGAAGACAACAAAGTTGGATT |
|  | 3-F | ACCG CACCTTCC AAGT TTGTCTTCGAAGACAA TTTG CGGGACAG G |
|  | 3-R | TAAA CCTGTCCCGCAAATTGTCTTCGAAGACAAACTTGGAAGGTG |
|  | 4-F | ACCG ACAGGAA AACG TTGTCTTCGAAGACAA AAGT CCTGAAGG G |
|  | 4-R | TAAA CCCTTCAGGACTTTTGTCTTCGAAGACAACGTTTTCCTGT |
|  | 5-F | ACCG T CTAG TTGTCTTCGAAGACAA AACG GCCTCCTTC G |
|  | 5-R | TAAA CGAAGGAGGCCGTTTTGTCTTCGAAGACAACTAGA |
| C2 | 1-F | ACCG ATTTGTTC AGCC TTGTCTTCGAAGACAA CATGTG |
|  | 1-R | TAAA CACATGTTGTCTTCGAAGACAAGGCTGAACAAAT |
|  | 2-F | ACCG TTTCGAGG CCCT TTGTCTTCGAAGACAA AGCC TTGCATCG G |
|  | 2-R | TAAA CCGATGCAAGGCTTTGTCTTCGAAGACAAAGGGCCTCGAAA |
|  | 3-F | ACCG CCTGGTGA TGAA TTGTCTTCGAAGACAA CCCT GCTCATCT G |
|  | 3-R | TAAA CAGATGAGCAGGGTTGTCTTCGAAGACAATTCATCACCAGG |
|  | 4-F | ACCG TTAGAGCT TGTG TTGTCTTCGAAGACAA TGAA AGGTGTAG G |
|  | 4-R | TAAA CCTACACCTTTCATTGTCTTCGAAGACAACACAAGCTCTAA |
|  | 5-F | ACCG T CTAG TTGTCTTCGAAGACAA TGTG ACCTGTTG G |
|  | 5-R | TAAA CCAACAGGTCACATTGTCTTCGAAGACAACTAGA |
| C3 | 1-F | ACCG CAAGGGCC CCTG TTGTCTTCGAAGACAA CATGTG |
|  | 1-R | TAAA CACATGTTGTCTTCGAAGACAACAGGGGCCCTTG |
|  | 2-F | ACCG GTAGGCCC TTGA TTGTCTTCGAAGACAA CCTG GCTTCTCA G |
|  | 2-R | TAAA CTGAGAAGCCAGGTTGTCTTCGAAGACAATCAAGGGCCTAC |
|  | 3-F | ACCG TCTCTTTC TCGA TTGTCTTCGAAGACAA TTGA GAAGCCAG G |
|  | 3-R | TAAA CCTGGCTTCTCAATTGTCTTCGAAGACAATCGAGAAAGAGA |
|  | 4-F | ACCG GCAGGACA CCAG TTGTCTTCGAAGACAA TCGA CTGCTCTT G |
|  | 4-R | TAAA CAAGAGCAGTCGATTGTCTTCGAAGACAACTGGTGTCCTGC |
|  | 5-F | ACCG T CTAG TTGTCTTCGAAGACAA CCAG GAGAGGCA G |
|  | 5-R | TAAA CTGCCTCTCCTGGTTGTCTTCGAAGACAACTAGA |
| CARGO oligos for CRISPRi in GM12878 cells | | |
| E1 | 1-F | ACCG CCTGGTCT TGTA TTGTCTTCGAAGACAA CATGTG |
|  | 1-R | TAAA CACATGTTGTCTTCGAAGACAATACAAGACCAGG |
|  | 2-F | ACCG ATTACCCC ATGA TTGTCTTCGAAGACAA TGTA ATTCACAG G |
|  | 2-R | TAAA CCTGTGAATTACATTGTCTTCGAAGACAATCATGGGGTAAT |
|  | 3-F | ACCG GCTGTATA GTGT TTGTCTTCGAAGACAA ATGA ATTGAAAG G |
|  | 3-R | TAAA CCTTTCAATTCATTTGTCTTCGAAGACAAACACTATACAGC |
|  | 4-F | ACCG CATGGGGA ATTA TTGTCTTCGAAGACAA GTGT ATGCTCAG G |
|  | 4-R | TAAA CCTGAGCATACACTTGTCTTCGAAGACAATAATTCCCCATG |
|  | 5-F | ACCG TCTAG TTGTCTTCGAAGACAA ATTA ATTTATTT G |
|  | 5-R | TAAA CAAATAAATTAATTTGTCTTCGAAGACAACTAGA |
| E2 | 1-F | ACCG ATACTTTA AGCC TTGTCTTCGAAGACAA CATGTG |
|  | 1-R | TAAA CACATGTTGTCTTCGAAGACAAGGCTTAAAGTAT |
|  | 2-F | ACCG GAAAGTCC CAGA TTGTCTTCGAAGACAA AGCC TCTCATAA G |
|  | 2-R | TAAA CTTATGAGAGGCTTTGTCTTCGAAGACAATCTGGGACTTTC |
|  | 3-F | ACCG CACTGGCC AAGA TTGTCTTCGAAGACAA CAGA ATGACTGC G |
|  | 3-R | TAAA CGCAGTCATTCTGTTGTCTTCGAAGACAATCTTGGCCAGTG |
|  | 4-F | ACCG AAAATTAC CTAA TTGTCTTCGAAGACAA AAGA GTTCACAA G |
|  | 4-R | TAAA CTTGTGAACTCTTTTGTCTTCGAAGACAATTAGGTAATTTT |
|  | 5-F | ACCG TGAGGAAG CAGG TTGTCTTCGAAGACAA CTAA ACCAACTG G |
|  | 5-R | TAAA CCAGTTGGTTTAGTTGTCTTCGAAGACAACCTGCTTCCTCA |
|  | 6-F | ACCG CCAGTGGT TTAT TTGTCTTCGAAGACAA CAGG CTTTCACC G |
|  | 6-R | TAAA CGGTGAAAGCCTGTTGTCTTCGAAGACAAATAAACCACTGG |
|  | 7-F | ACCG CTACAGGT AGTG TTGTCTTCGAAGACAA TTAT CATGCTGA G |
|  | 7-R | TAAA CTCAGCATGATAATTGTCTTCGAAGACAACACTACCTGTAG |
|  | 8-F | ACCGGTTGTGAA AGTT TTGTCTTCGAAGACAA AGTG CCAAATCG G |
|  | 8-R | TAAA CCGATTTGGCACTTTGTCTTCGAAGACAAAACTTTCACAAC |
|  | 9-F | ACCG TCTAG TTGTCTTCGAAGACAA AGTT TCCTTGAG G |
|  | 9-R | TAAA CCTCAAGGAAACTTTGTCTTCGAAGACAACTAGA |
| E3 | 1-F | ACCG CCTTTGAA CATG TTGTCTTCGAAGACAA CATGTG |
|  | 1-R | TAAA CACATGTTGTCTTCGAAGACAACATGTTCAAAGG |
|  | 2-F | ACCG ACACATAA CTGG TTGTCTTCGAAGACAA CATG CTCTATTT G |
|  | 2-R | TAAA CAAATAGAGCATGTTGTCTTCGAAGACAACCAGTTATGTGT |
|  | 3-F | ACCG GGCCTGCT GAAC TTGTCTTCGAAGACAA CTGG ATTTTTTG G |
|  | 3-R | TAAA CCAAAAAATCCAGTTGTCTTCGAAGACAAGTTCAGCAGGCC |
|  | 4-F | ACCG CTTCTCCC TCCA TTGTCTTCGAAGACAA GAAC ATTGACAA G |
|  | 4-R | TAAA CTTGTCAATGTTCTTGTCTTCGAAGACAATGGAGGGAGAAG |
|  | 5-F | ACCG CCATGTAG GTGG TTGTCTTCGAAGACAA TCCA AAGGCATC G |
|  | 5-R | TAAA CGATGCCTTTGGATTGTCTTCGAAGACAACCACCTACATGG |
|  | 6-F | ACCG GAGGTGACC ATCG TTGTCTTCGAAGACAA GTGG GATAGTCT G |
|  | 6-R | TAAA CAGACTATCCCACTTGTCTTCGAAGACAACGATGGTCACCTC |
|  | 7-F | ACCG AGGAATGG AAGG TTGTCTTCGAAGACAA ATCG TACACTG G |
|  | 7-R | TAAA CCAGTGTACGATTTGTCTTCGAAGACAACCTTCCATTCCT |
|  | 8-F | ACCG ACTGAATC TTGA TTGTCTTCGAAGACAA AAGG GTTGGTTG G |
|  | 8-R | TAAA CCAACCAACCCTTTTGTCTTCGAAGACAATCAAGATTCAGT |
|  | 9-F | ACCG TCTAG TTGTCTTCGAAGACAA TTGA AACATTAC G |
|  | 9-R | TAAA CGTAATGTTTCAATTGTCTTCGAAGACAACTAGA |
| E4 | 1-F | ACCG AGTGGCGA CTTC TTGTCTTCGAAGACAA CATGTG |
|  | 1-R | TAAA CACATGTTGTCTTCGAAGACAAGAAGTCGCCACT |
|  | 2-F | ACCG AACTTTCA GTCG TTGTCTTCGAAGACAA CTTC TTGCAACT G |
|  | 2-R | TAAA CAGTTGCAAGAAGTTGTCTTCGAAGACAACGACTGAAAGTT |
|  | 3-F | ACCG AGGAGGGG GGAG TTGTCTTCGAAGACAA GTCG GCTGGAGG G |
|  | 3-R | TAAA CCCTCCAGCCGACTTGTCTTCGAAGACAACTCCCCCCTCCT |
|  | 4-F | ACCG GCCCGGGC TCGG TTGTCTTCGAAGACAA GGAG AAGGGGGG G |
|  | 4-R | TAAA CCCCCCCTTCTCCTTGTCTTCGAAGACAACCGAGCCCGGGC |
|  | 5-F | ACCG CTAGTGCA CAGC TTGTCTTCGAAGACAA TCGG CCGCGCAC G |
|  | 5-R | TAAA CGTGCGCGGCCGATTGTCTTCGAAGACAAGCTGTGCACTAG |
|  | 6-F | ACCG TGGGGG TGTC TTGTCTTCGAAGACAA CAGC CTGCTTTT G |
|  | 6-R | TAAA CAAAAGCAGGCTGTTGTCTTCGAAGACAAGACACCCCCA |
|  | 7-F | ACCG GCAATGAA AAGA TTGTCTTCGAAGACAA TGTC GGCAGACCAC G |
|  | 7-R | TAAA CGTGGTCTGCCGACATTGTCTTCGAAGACAATCTTTTCATTGC |
|  | 8-F | ACCG AGAACCAT GTAC TTGTCTTCGAAGACAA AAGA CATGATTA G |
|  | 8-R | TAAA CTAATCATGTCTTTTGTCTTCGAAGACAAGTACATGGTTCT |
|  | 9-F | ACCG ACACTGAA ATCA TTGTCTTCGAAGACAA GTAC TTGTTCTT G |
|  | 9-R | TAAA CAAGAACAAGTACTTGTCTTCGAAGACAATGATTTCAGTGT |
|  | 10-F | ACCG GGTCTCC TGCA TTGTCTTCGAAGACAA ATCA GAAACCTG G |
|  | 10-R | TAAA CCAGGTTTCTGATTTGTCTTCGAAGACAATGCAGGAGACC |
|  | 11-F | ACCG TCTAG TTGTCTTCGAAGACAA TGCA GTACAAGAG G |
|  | 11-R | TAAA CCTCTTGTACTGCATTGTCTTCGAAGACAACTAGA |

| RT-qPCR primers | | | | |
| --- | --- | --- | --- | --- |
| LINC00158 | F | | TGAGCAGAACACATGCACAA | |
|  | R | | CGACGACCGCTTTCTTAAAC | |
| MIR155HG | F | | GCAGGTTTTGGCTTGTTCAT | |
|  | R | | AAAACGTTGCCAGACAATCC | |
| MRPL39 | F | | GCTGTCACCGACAGAATTGA | |
|  | R | | CCAGAGCCAGAATGGACTTC | |
| JAM2 | F | | TGCTCTGAGTGGAACTGTGG | |
|  | R | | CACCTGCGATATCCAACAGA | |
| ATP5J | F | | CTGGAGGACCTGTTGATGCT | |
|  | R | | TGGGGTTTTTCGATGACTTC | |
| GABPA | F | | AAGTGACAAGATGGGCTGCT | |
|  | R | | CCGAAATGTTGAGTGTGGTG | |
| CACNA1C | F | | AGTCCGTCAACACCGAAAAC | |
|  | R | | CCAGTTGGGCTGGTTGTAGT | |
| ChIP-qPCR primers | | | | |
| E1 | F | | GAAACGCACAGCTGGAATAA | |
|  | R | | CCAGGCTAATGATGCTGTGA | |
| E2 | F | | CTTTGATTGGGGTGCTCTGT | |
|  | R | | GAAAGAAGGGCCATATTCCA | |
| E3 | F | | TTCCCCAGGGATGCTTTAAT | |
|  | R | | TTCAGCTCACGAAGACATGC | |
| E4 | F | | CCACCCAGTTGCAAGAAGTC | |
|  | R | | CACGCCGTGTACTTTCCTG | |
| 3C-qPCR primers | | | | |
| Fragment | | Strand | | sequence |
| * chr12:2391693-2395405 | | + | | GAGCTTTGAAACAAGACTGGCC |
| Chr12:2140738-2146957 | | + | | AACcctcacttctcagagcatg |
| Chr12:2147580-2149974 | | + | | tgacagaaaattccatctttgTGGt |
| Chr12:2150113-2155869 | | + | | GCTGAAAATCACAAGGCTAGTCA |
| Chr12:2155869-2174302 | | + | | CTGTGCAGAGCTGGAATGTTG |
| Chr12:2174302-2176247 | | + | | CGAGGTCTGGAAGCATGAGTTA |
| * Anchor primer | | | | |
