## Supplemental figures for "Robust Hi-C chromatin loop maps in human neurogenesis and brain tissues at high-resolution"

### Extended Data Figure 1

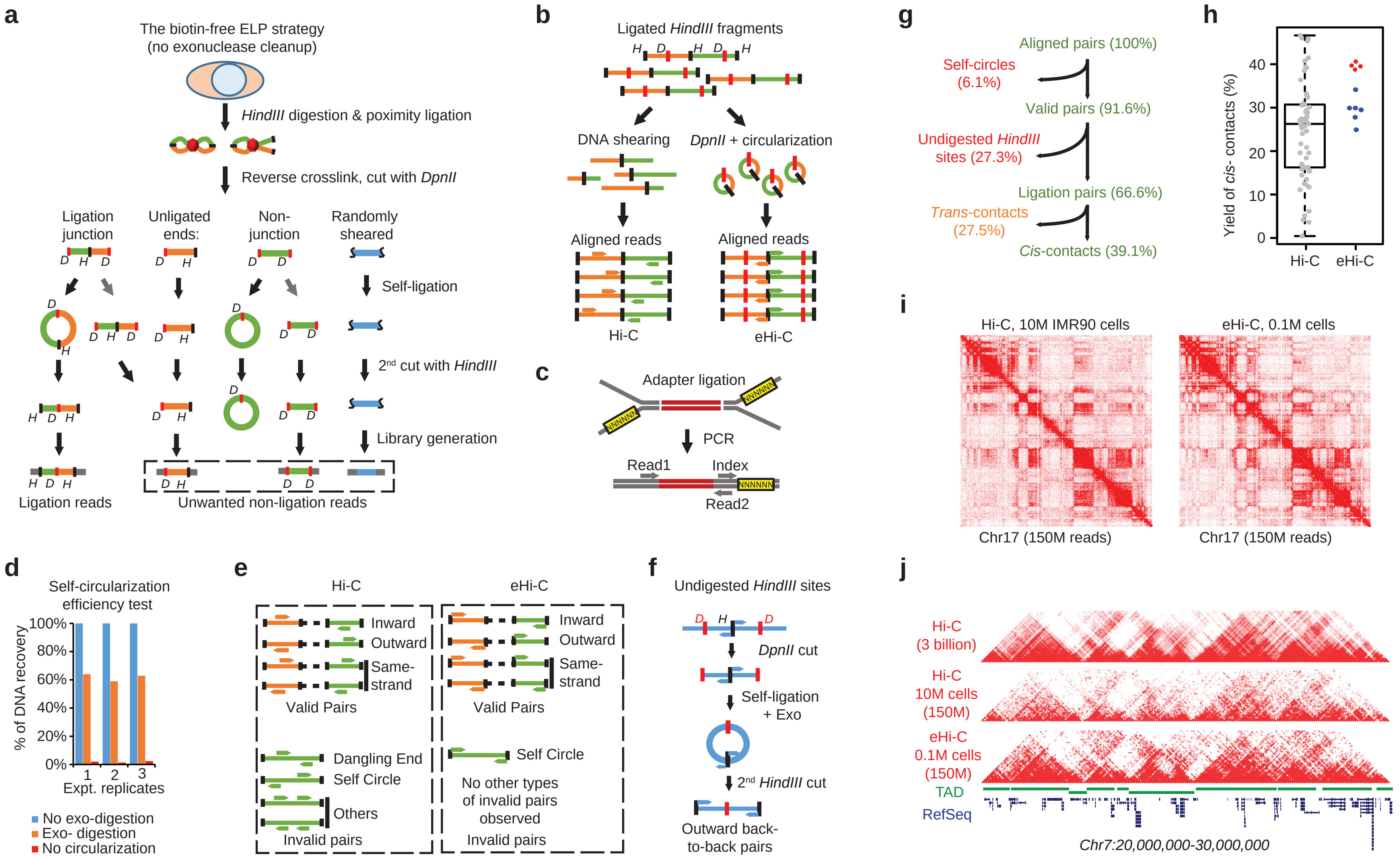

**Extended Data Figure 1. Source of errors from eHi-C protocol.**

**a,** The scheme shows the different types of unwanted non-ligation reads (in dashed box) when the exonuclease cleanup step is omitted from eHi-C protocol (which is equivalent to the previously published ELP method[1]). All these non-ligation reads can be effectively removed by exonuclease cleanup. **b**, For reproducible ligation DNA products between the same HindIII ends, (Left) Hi-C will generate different paired-end reads due to random DNA shearing. (Right) In eHi-C, these ligation events will form the same DNA circles, resulting in identical paired-end reads. **c**, We used a custom adapter with 6 random bases as a unique molecule index (UMI) to distinguish PCR duplicates. **d**, Test the efficiency of self-ligation. Naked genomic DNA were first digested with DpnII. Self-ligation reactions (2.5µg DNA in 1mL to mimic the eHi-C condition) were then performed before λ-exonuclease treatment. The efficiency of self-ligation was measured by the percentage of remaining DNA after exonuclease digestion (orange). No exonuclease controls are set at 100% (blue). DNA with no self-ligation step are completely digested (red). Results from 3 independent assays are shown. **e**, Valid Hi-C or eHi-C reads involve two *HindIII* fragments can be further classified based on strand orientation. The invalid Hi-C reads include "dangling end", "self-circle" and "others" based on strand orientation, while the only type of invalid eHi-C reads is self-circles (right). **f**, Undigested HindIII sites are one major source of errors in eHi-C, which are read pairs mapped back-to-back at the same HindIII site. **g**, Data filtering results of one exemplary eHi-C library generated from 0.1M cells. **h**, Compare the yield of *cis*-contact reads between 57 published Hi-C libraries and 10 eHi-C libraries. The 4 red spots are eHi-C libraries prepared under *in situ* ligation condition. **i**, Heatmaps of contact matrices (chr17) from Hi-C and eHi-C at 250kb resolution. **j**, Heatmaps of contact matrices from Hi-C and eHi-C at 50kb resolution. The top track is drawn using a published IMR90 Hi-C dataset with ~3 billion reads[2]. A track of TAD structures is plotted in green.

### Extended Data Figure 2

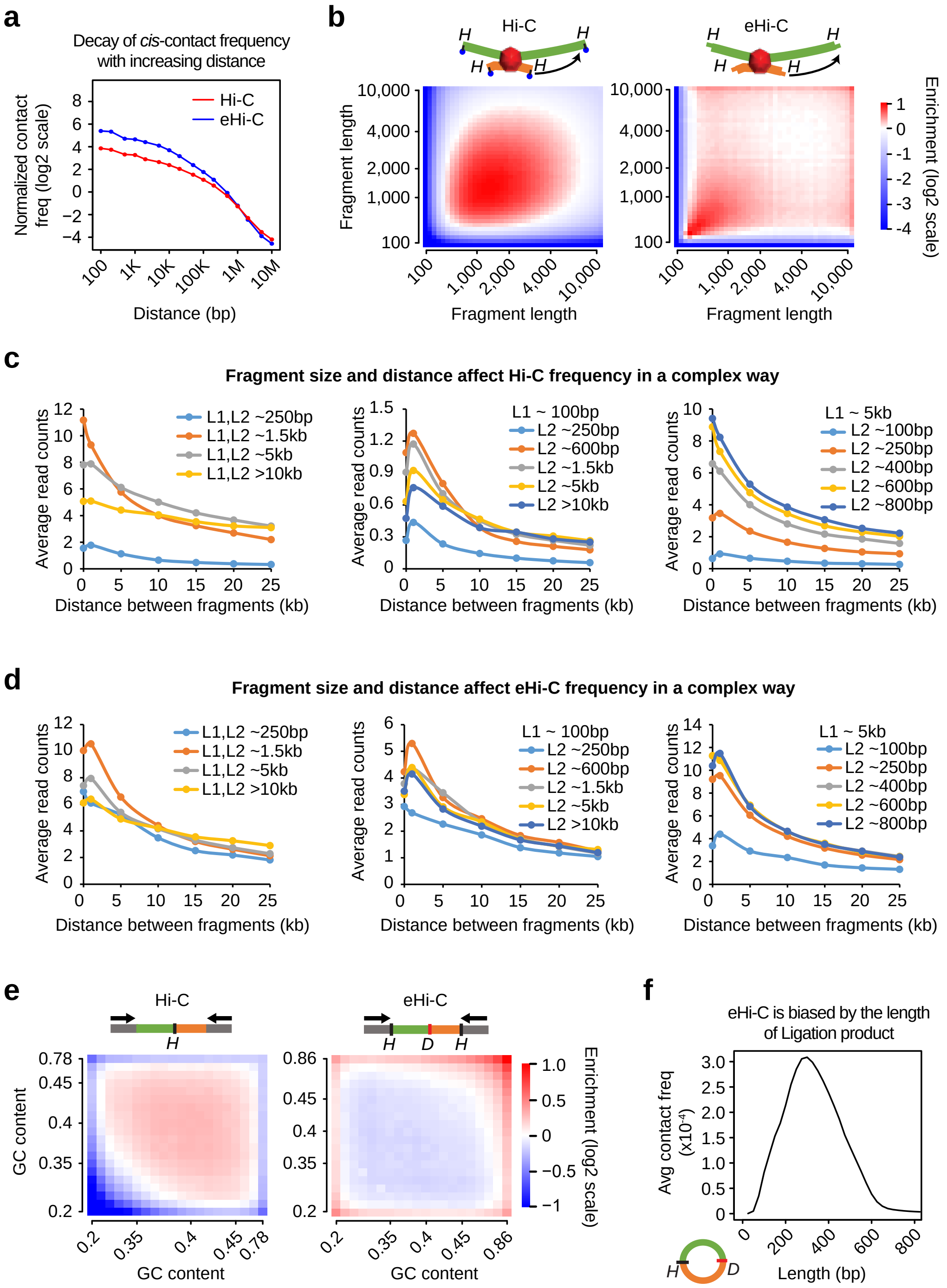

**Extended Data Figure 2. Systematical biases in eHi-C experiments.**

**a**, Curves showing the decay of *cis*-contact with an increasing distance between two *HindIII* restrictive fragments. Only "same-strand" reads (see **Methods**) were used to plot the curves. **b**, Compare the bias from *HindIII* fragment length in Hi-C (left) and eHi-C (right) libraries. All the fragments are binned into 40 equal-sized groups, and the enrichment of *trans* reads between any two groups are plotted as heatmaps. The enrichment value is the ratio between actual read counts and the global average for any two groups. **c**, Curves plot how the distance decay profile changes when the length of *HindIII* fragments are different in H1 hESC Hi-C data. **d**, The same analyses as **c** are performed with iPSC eHi-C data. **e**, *HindIII* ends are binned into 20 groups based on GC content, and the enrichment of *trans*- reads are also plotted as heatmaps. For Hi-C (left) we used the GC content in the 200bp region upstream the *HindIII* site, and for eHi-C, we used the GC content of the region between the *HindIII* and its nearest *DpnII* site. **f**, Curve shows the average contact frequency from eHi-C against the length of ligation junction products forming DNA circles.

### Extended Data Figure 3

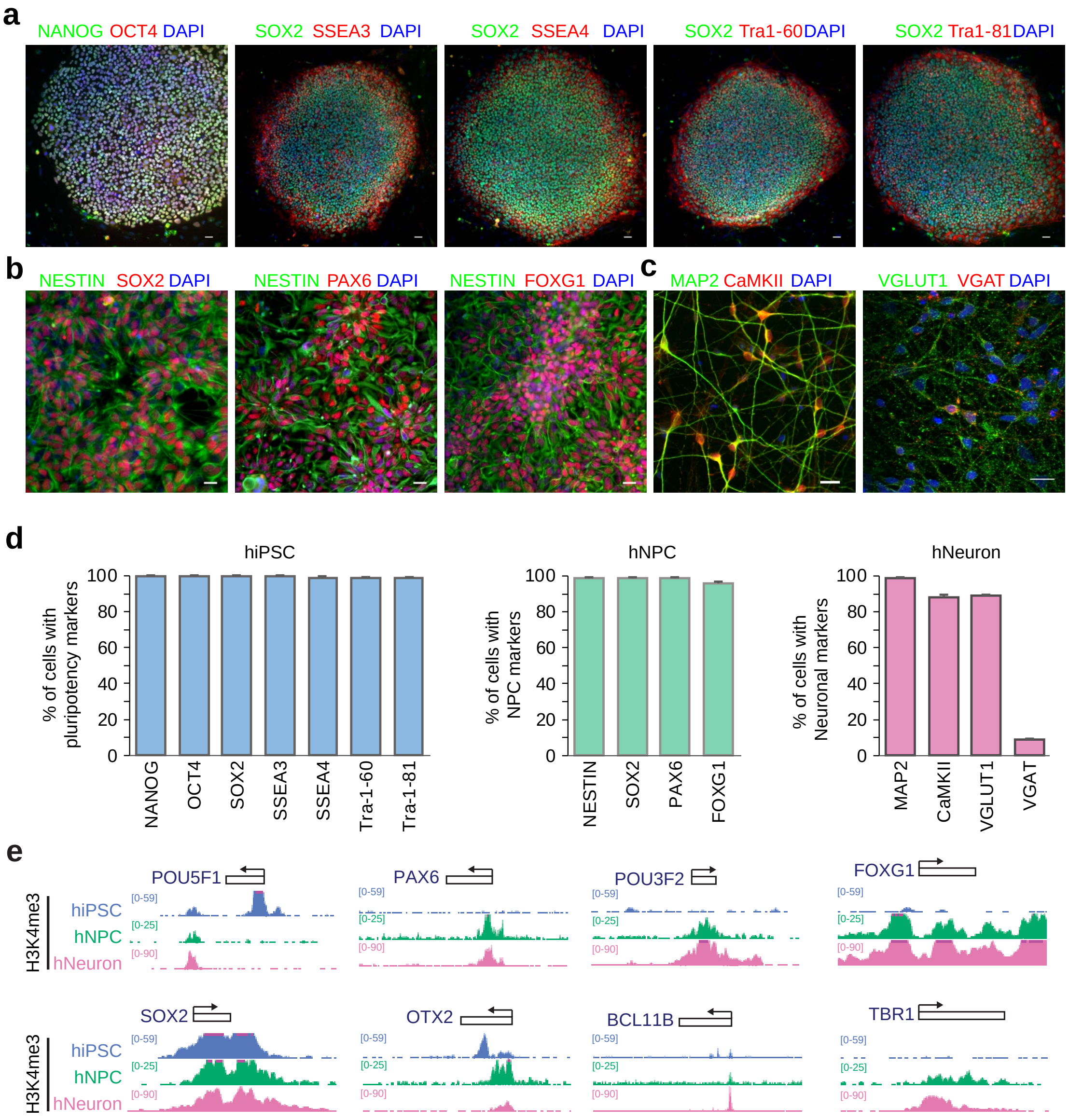

**Extended Data Figure 3. hiPSC characterization and forebrain-specific neural differentiation.**

**a**, Confocal images of pluripotency marker immunostaining of hiPSCs. **b**, Confocal images of neural progenitor marker immunostaining of hNPCs. **c**, Confocal images of immunostaining of MAP2AB, CAMKII, VGLUT1 and VGAT. Scale bars, 20 μm. **d**, Quantification of cells with pluripotency, NPC and neuronal markers from flurorescent images. Values represent mean+/-SEM. **e**, H3K4me3 ChIPmentation results at representative marker genes.

### Extended Data Figure 4

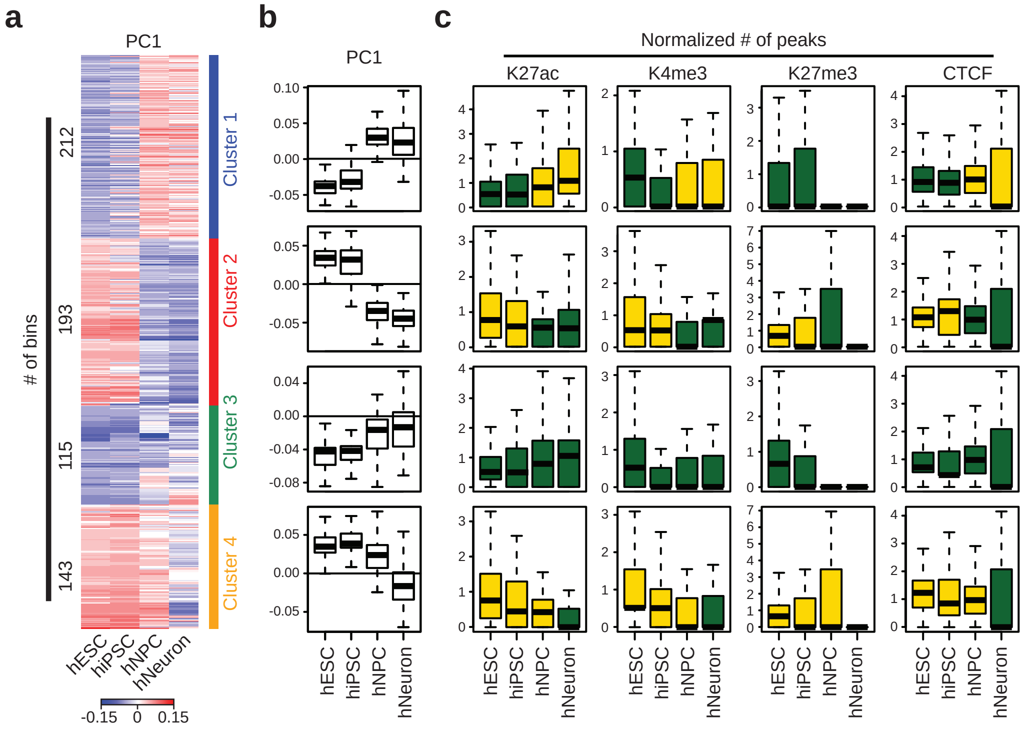

**Extended Data Figure 4. Genome compartment changes in neuronal differentiation.**

**a**, After excluding reprogramming DCRs, the 663 DCRs (bins) during neuronal differentiation are classified into 4 clusters based on PC1 value representing different patterns of compartment switching. **b**, Box plots summarize the PC1 values of each DCR cluster. **c**, Box plots showing normalized numbers of histone mark peaks from DCRs in each cluster.

### Extended Data Figure 5

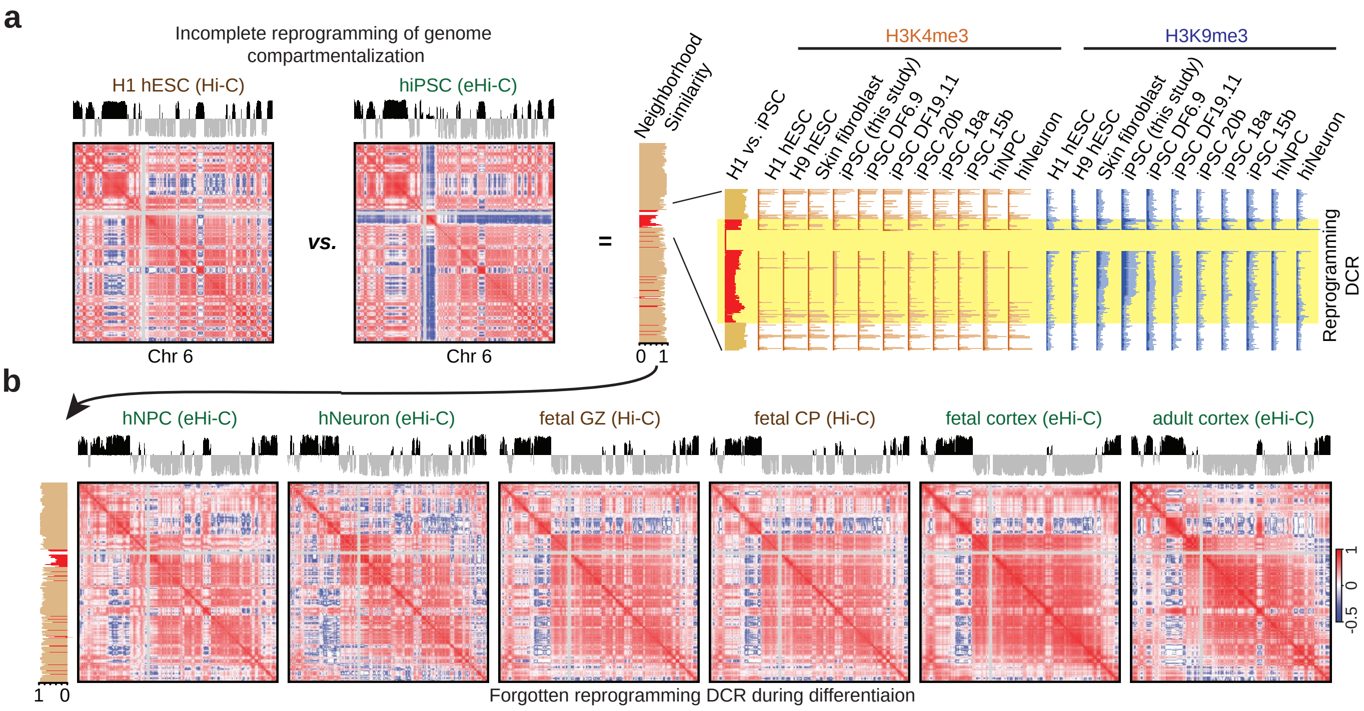

**Extended Data Figure 5. An ultra-large locus in chr6 with 3D memory of somatic heterochromatin in hiPSC reprogramming.**
**a**, Similar to **Fig. 2a-b**, another example of reprogramming DCR in chr6. **b**, The hiPSC specific architectural difference is largely removed after differentiation.

### Extended Data Figure 6

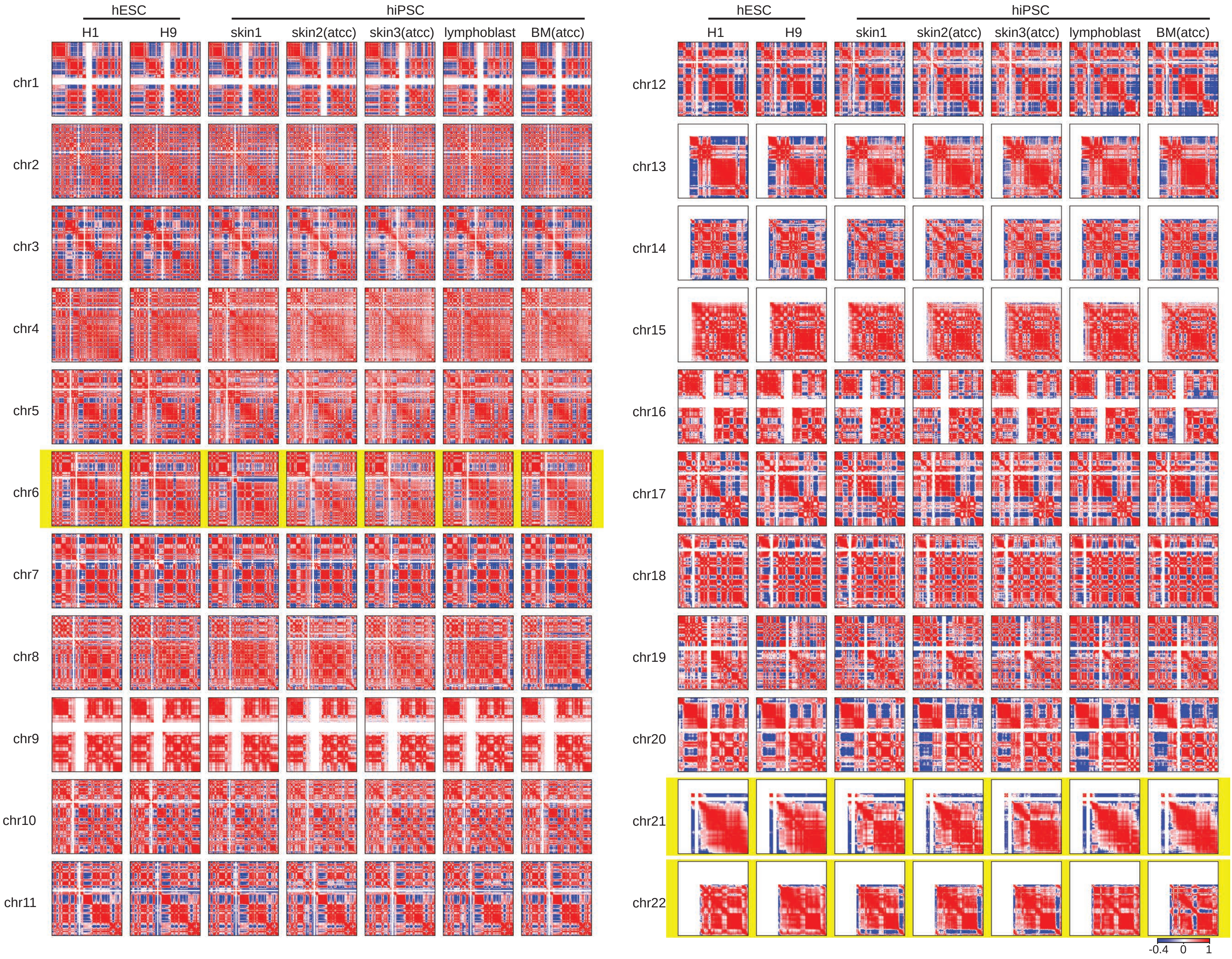

**Extended Data Figure 6. Compare the genome compartmentalization in hESCs and hiPSCs.** For each human chromosome, we show the correlation matrix heatmaps of two ES cells (H1 and H9), three skin derived hiPSCs (skin1, generated in this study; skin2, ATCC ACS-1019; skin3 ATCC ACS-1011), a published lymphoblast derived hiPSC[3], and a bone marrow derived hiPSC (ATCC ACS-1026). The three chromosomes (chr6, chr21, chr22) containing ultra-large loci with recurrent 3D memory of somatic heterochromatin are highlighted.

#### Extended Data Figure 7

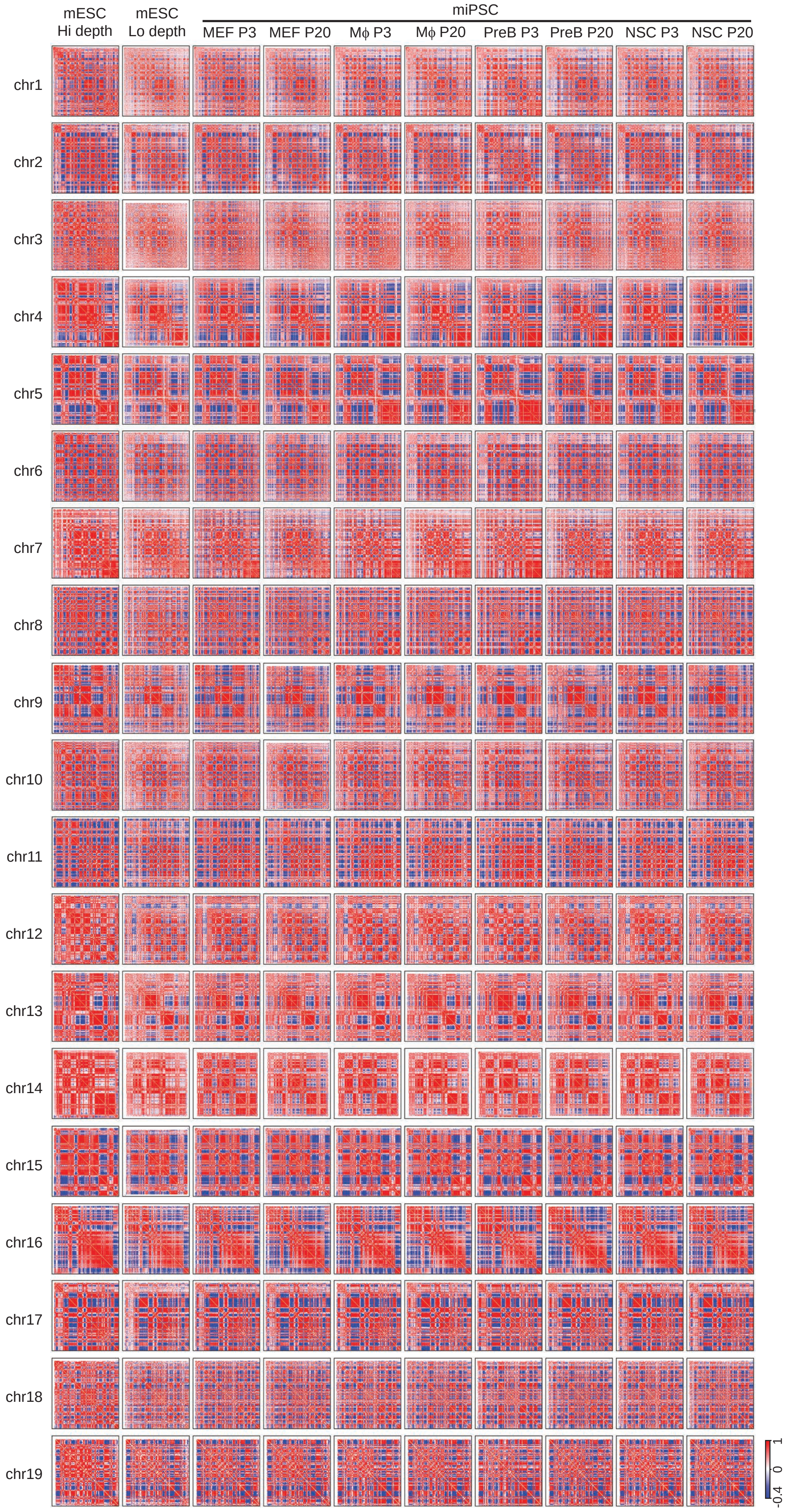

**Extended Data Figure 7. Compare the genome compartmentalization of mESC and miPSCs.** For each mouse chromosome, we show the correlation matrix heatmaps of two mESC and eight miPSCs. Low depth mESC and all miPSC heatmaps are from the reanalysis of the data in a previous publication[4]. High depth mESC data are an independent dataset[5]. There is no evidence of somatic 3D genome memory at compartment level.

### Extended Data Figure 8

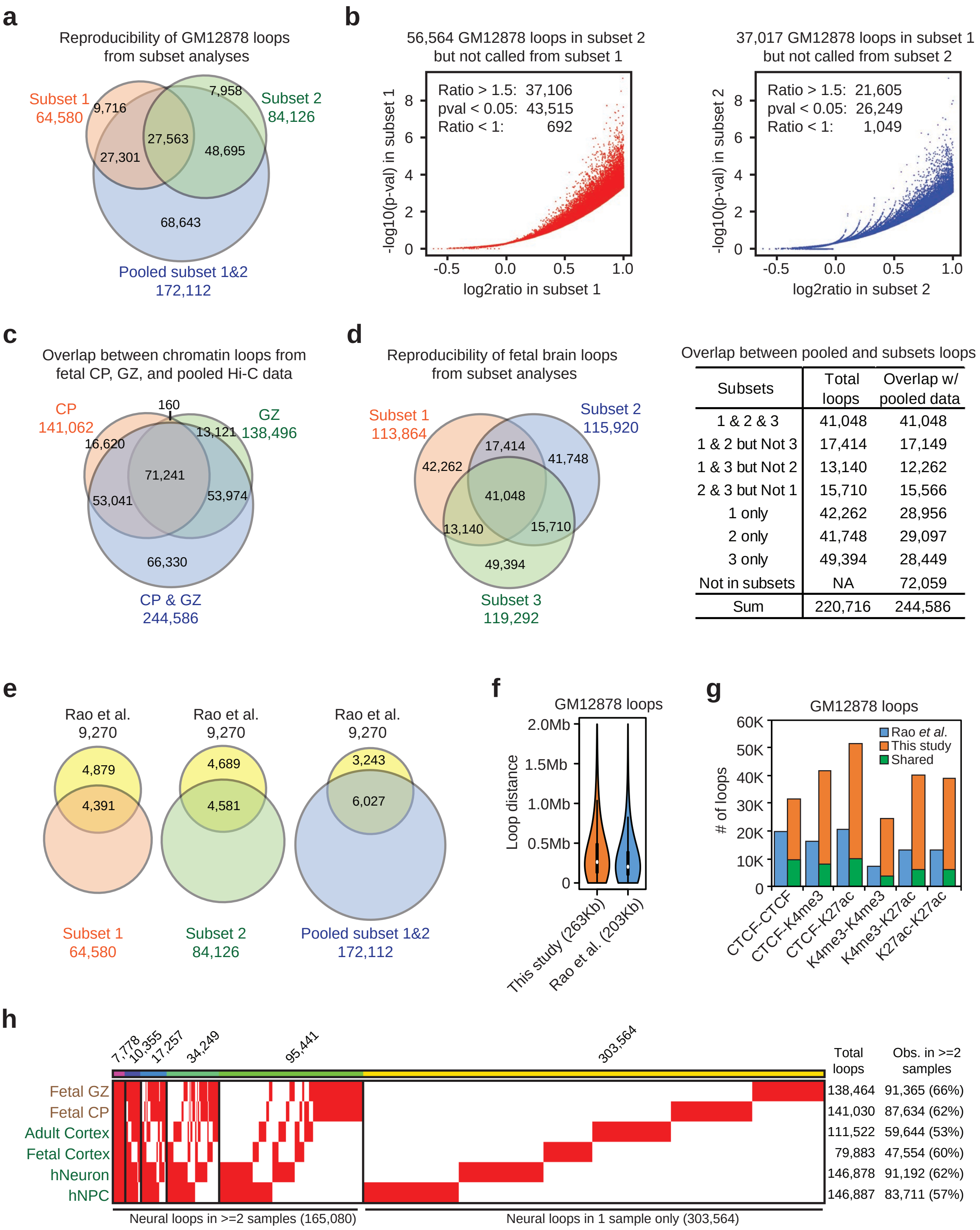


| Subsets | Total loops | Overlap w/ pooled data |
|---|---|---|
| 1 & 2 & 3 | 41,048 | 41,048 |
| 1 & 2 but Not 3 | 17,414 | 17,149 |
| 1 & 3 but Not 2 | 13,140 | 12,262 |
| 2 & 3 but Not 1 | 15,710 | 15,566 |
| 1 only | 42,262 | 28,956 |
| 2 only | 41,748 | 29,097 |
| 3 only | 49,394 | 28,449 |
| Not in subsets | NA | 72,059 |
| Sum | 220,716 | 244,586 |

**Extended Data Figure 8. Reproducibility of chromatin loop calls.**

**a**, Venn diagram show the overlap of loop calls (ratio > 2 and $p < 0.001$, **Supplementary Method**) between two subsets of GM12878 data (involving different biological replicates from different labs, **Extended Data Table 4**), and the pooled datasets. **b**, Left: For all loops called from subset 2 but not subset 1 GM12878 Hi-C data, the scatter plot show their ratios and p values in subset 1. The results shows that a majority of these non-reproducible loops actually have positive loop, but not significant enough to be called. Right: Scatter plot similar to left showing the results of non-reproducible loops in subset 1. **c**, The overlap between cortex CP and GZ loops, and the loops after pooling CP and GZ data together. Note CP and GZ are two very close cortex regions and the data are generated by the same lab[6]. **d**, Left: The overlap of loop calls after split the CP and GZ data into 3 subsets, each with 1 CP and 1 GZ replicate (**Extended Data Table 5**). Right: The overlap between loops in the three-way subset analysis and loops called after pooling the CP and GZ data together. Note that the pooled datasets capture nearly all the reproducible loops from subset analyses. **e**, The performance of our loop calls (6-cutter Hi-C) in recovering the loops called by Rao *et al*. (4-cutter in situ Hi-C) in GM12878 cells. **f**, Compare the loop distance distribution of the loop calls by our method and Rao et al. in GM12878 cells. **g**, The overlap of our loop calls and Rao *et al*. in GM12878 cells on *cis*-regulatory elements. Note that we call loops in the format of pixels, while Rao *et al*. loops in the format of a circle area with center and radius. To reconcile this, we compared our loop pixels with all the pixels covered by the Rao *et al*. loop circle area. Blue boxes: all pixels from Rao *et al*. loop circle areas; green: shared pixels between our method and Rao *et al*.; orange: new pixels called only in this study. Comparing the orange and green boxes, the fold increases at H3K4me3 and H3K27ac marked regions (>5 fold) are much higher than at CTCF sites, suggesting that our method called more enhancer and promoter looping. **h**, The overlap between loops from the 6 neural (e)Hi-C datasets. ~60% of loops from any neural dataset are reproduced by at least two samples. These are regarded as high-confidence neural loops in our analysis.

### Extended Data Figure 9

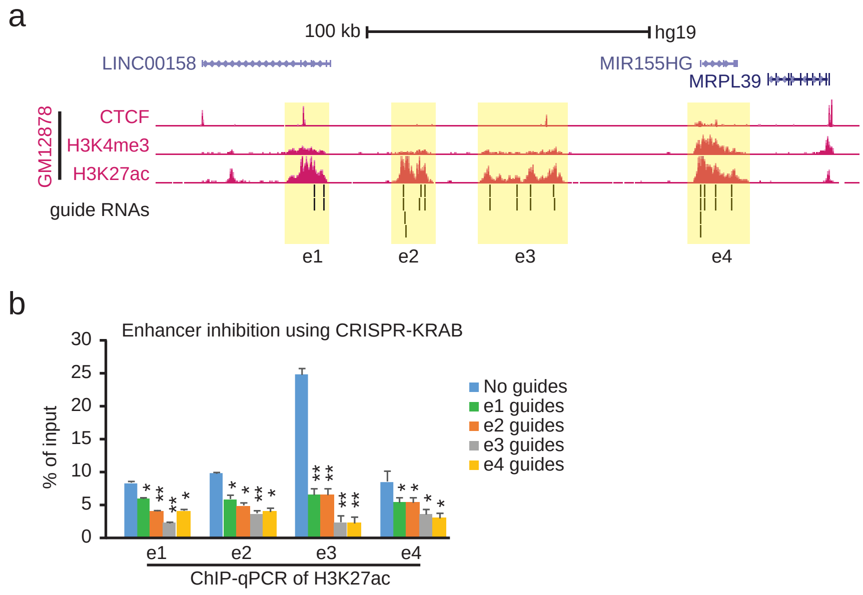

**Extended Data Figure 9. CRISPR inhibition of a GM12878 specific enhancer aggregate.**
**a**, Browser tracks showing the GM12878 ChIP-seq data and the locations of guide RNAs for the enhancer inhibition. **b**, ChIP-qPCR results showing the loss of H3K27ac occupancy after inhibiting each of enhancers.

### Extended Data Figure 10

**a**

Chromatin loops at cis-regulatory elements

| Chromatin loops | hiPSC | hNPC | hNeuron |
|---|---|---|---|
| w/ CTCF | 11,406 | 15,233 | 22,001 |
| w/ promoters | 5,102 | 7,748 | 10,393 |
| w/ enhancers | 3,619 | 7,533 | 13,689 |

**b**

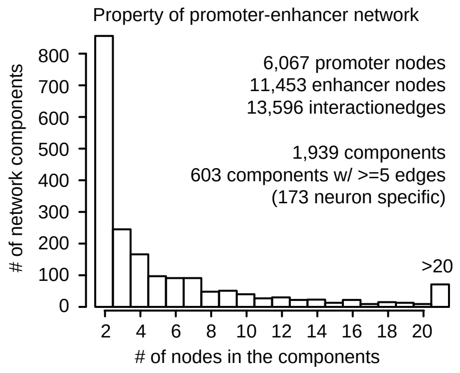

**Extended Data Figure 10. Summary of chromatin loops and network in neural differentiation.**

**a**, Summary of chromatin loop numbers overlapping different types of *cis*-regulatory elements in neural differentiation. **b**, Summary of components in our enhancer-promoter looping network analyses.

### Extended Data Figure 11

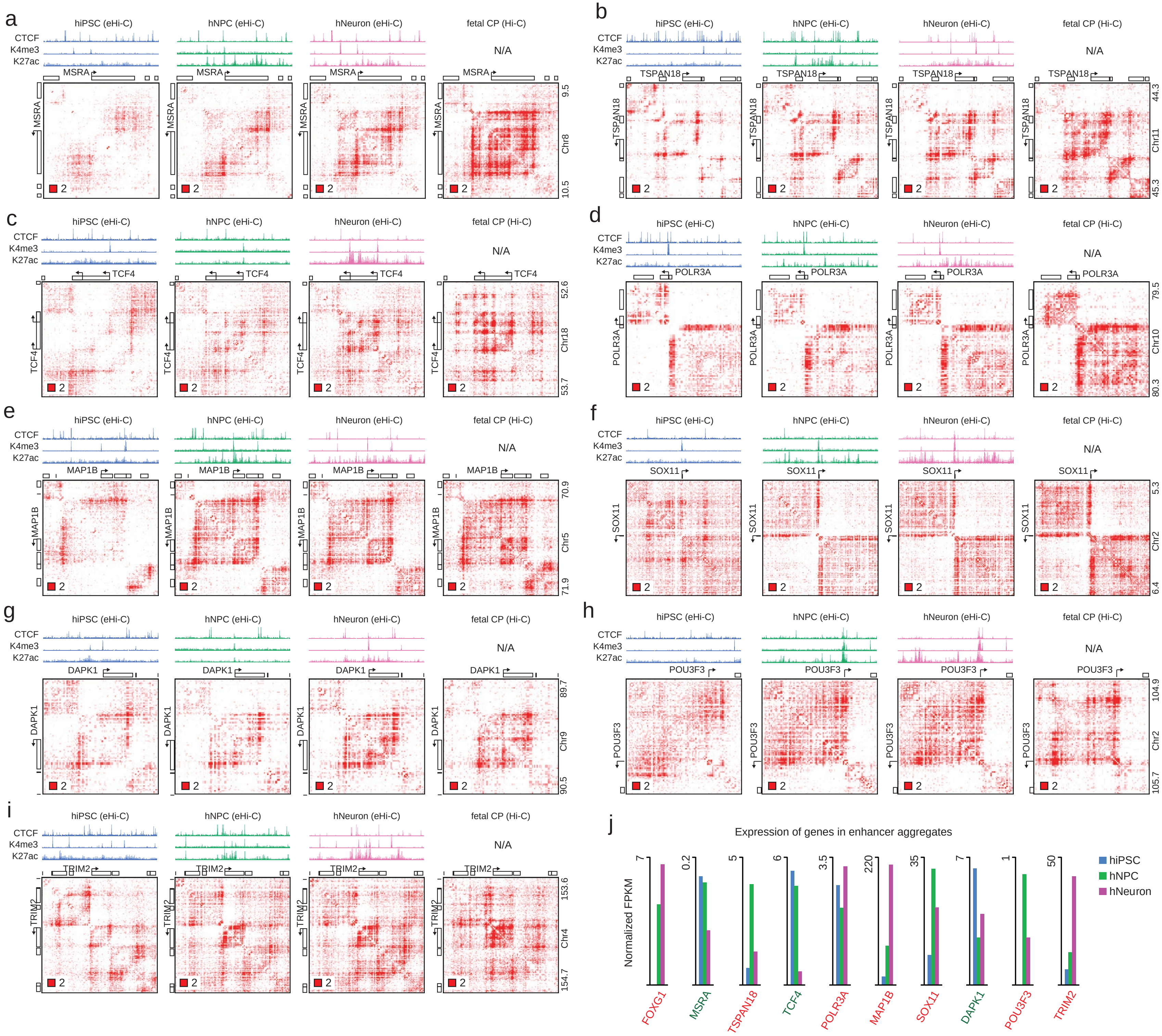

**Extended Data Figure 11. Example loci that gain enhancer aggregates in neural differentiation.**

**a-i**, Examples that gain chromatin loops during neural differentiation; heatmaps in hiPSC, hNPC, hNeuron and fetal CP are shown. **j**, RNA-seq expression data of key neural genes in these regions, including *FOXG1* gene shown in **Fig. 5f**. Genes in red are upregulated and in green are downregulated.

### Extended Data Figure 12

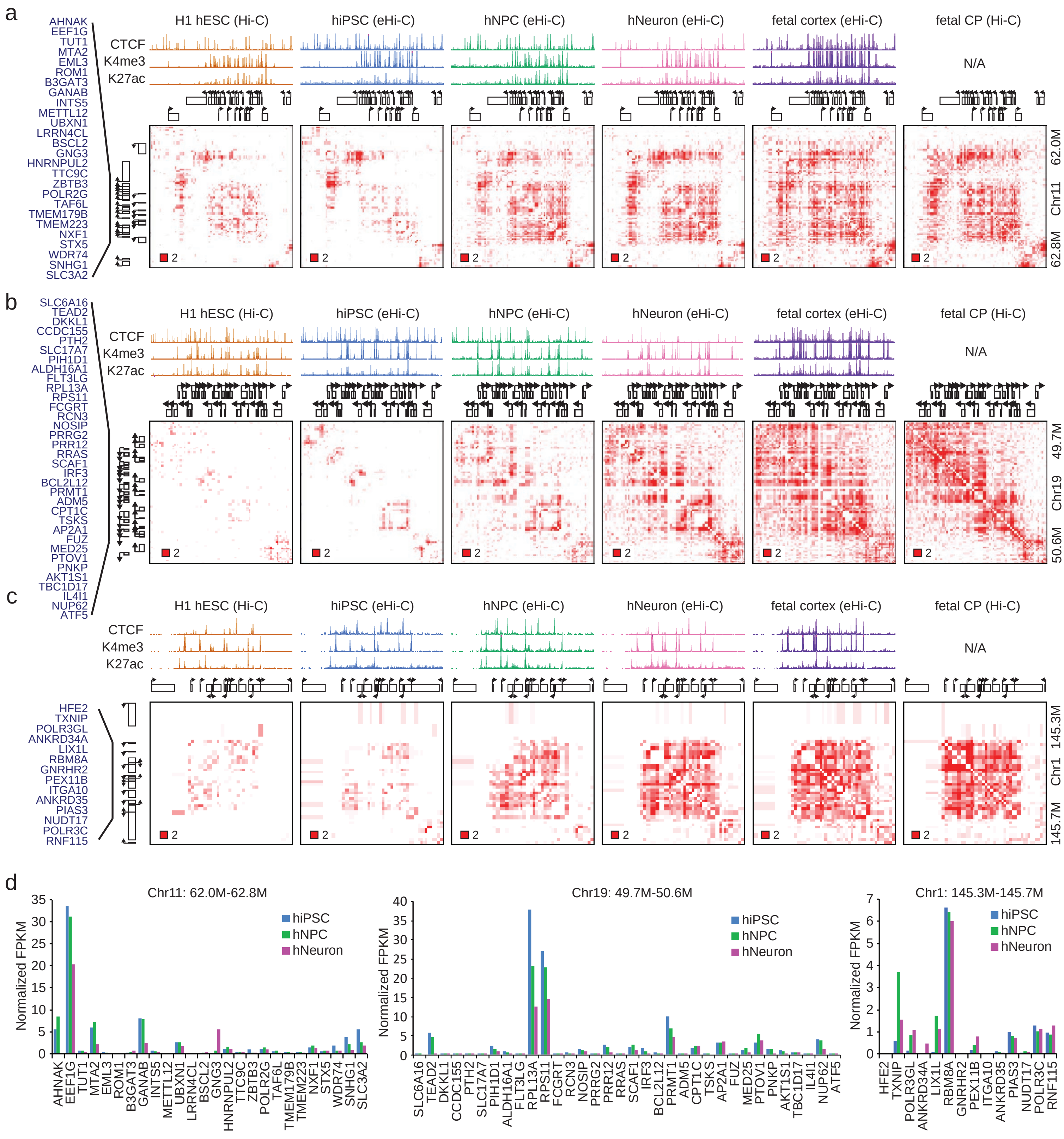

**Extended Data Figure 12. Genes in neural enhancer aggregates can be both up- and down-regulated in neural differentiation.**

**a-c**, Example enhancer aggregate regions that involve many promoters and enhancers. Gene names in the aggregated region are listed on the right. **d**, RNA-seq data of the genes in these enhancer aggregations demonstrated the coordinated gene downregulation in example **a** and **b**, and coordinated gene upregulation in example **c**.

### Extended Data Figure 13

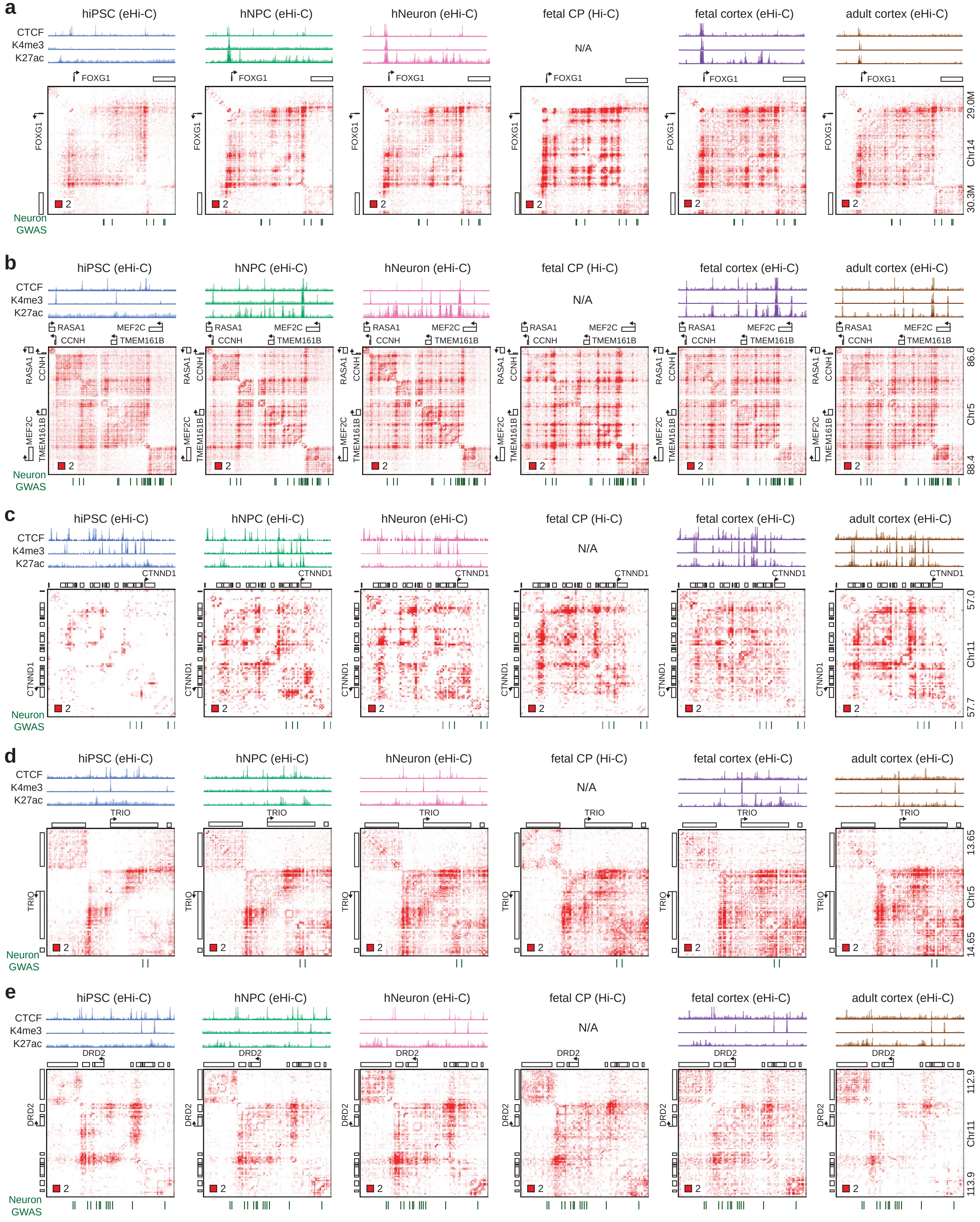

**Extended Data Figure 13. Examples of several known neural disease risk loci.**
For these examples, heatmaps of hiPSC, hNPC, hNeuron, fetal CP, fetal and adult cortex are shown to demonstrate the overall agreements between differentiated neurons and primary tissues. Note some examples gain new DNA contacts in differentiation, while some examples have pre-existing DNA contacts.

#### Extended Data Figure 14

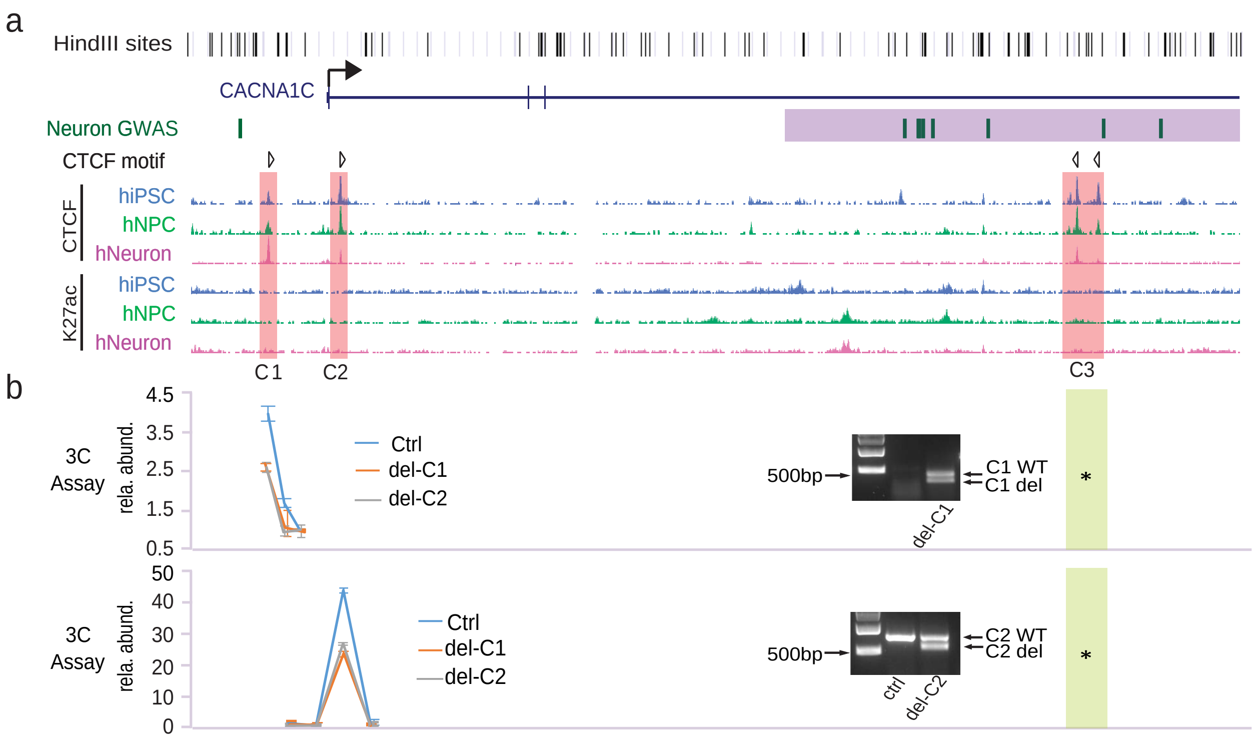

**Extended Data Figure 14. CTCF sites are responsible for the DNA loop at CACNA1C locus in hESC.**

**a**, Genomic features of *CACNA1C* locus. HindIII site track is shown as reference for 3C assays. C1, C2 and C3 are the CTCF sites with the motif direction shown above. **b**, 3C assays in H9 hESC cells. C1 and C2 sites were deleted respectively in H9 cells, and a nucleofection with no-sgRNA was used as control. Deletion efficiency was shown as in the gel figures. 3C assay was done in three biological replicates. The relative PCR abundance was calculated to a nearby region which shows no interaction with C3 region from Hi-C. The anchor fragment is highlighted in yellow. Error bars: *s.d.* from the three replicates.
